# Models of archaic admixture and recent history from two-locus statistics

**DOI:** 10.1101/489401

**Authors:** Aaron P. Ragsdale, Simon Gravel

## Abstract

We learn about population history and underlying evolutionary biology through patterns of genetic polymorphism. Many approaches to reconstruct evolutionary histories focus on a limited number of informative statistics describing distributions of allele frequencies or patterns of linkage disequilibrium. We show that many commonly used statistics are part of a broad family of two-locus moments whose expectation can be computed jointly and rapidly under a wide range of scenarios, including complex multi-population demographies with continuous migration and admixture events. A full inspection of these statistics reveals that widely used models of human history fail to predict simple patterns of linkage disequilibrium. To jointly capture the information contained in classical and novel statistics, we implemented a tractable likelihood-based inference framework for demographic history. Using this approach, we show that human evolutionary models that include archaic admixture in Africa, Asia, and Europe provide a much better description of patterns of genetic diversity across the human genome. We estimate that an unidentified, deeply diverged population admixed with modern humans within Africa both before and after the split of African and Eurasian populations, contributing 4 *-* 8% genetic ancestry to individuals in world-wide populations.

**Author Summary:** Throughout human history, populations have expanded and contracted, split and merged, and exchanged migrants. Because these events affected genetic diversity, we can learn about human history by comparing predictions from evolutionary models to genetic data. Here, we show how to rapidly compute such predictions for a wide range of diversity measures within and across populations under complex demographic scenarios. While widely used models of human history accurately predict common measures of diversity, we show that they strongly underestimate the co-occurence of low frequency mutations within human populations in Asia, Europe, and Africa. Models allowing for archaic admixture, the relatively recent mixing of human populations with deeply diverged human lineages, resolve this discrepancy. We use such models to infer demographic models that include both recent and ancient features of human history. We recover the well-characterized admixture of Neanderthals in Eurasian populations, as well as admixture from an as-yet unknown diverged human population within Africa, further suggesting that admixture with deeply diverged lineages occurred multiple times in human history. By simultaneously testing model predictions for a broad range of diversity statistics, we can assess the robustness of common evolutionary models, identify missing historical events, and build more informed models of human demography.

## Introduction

The study of genetic diversity in human populations has shed light on the origins of our species and our spread across the globe. With the growing abundance of sequencing data from contemporary and ancient humans, coupled with archaeological evidence and detailed models of human demography, we continue to refine our understanding of our intricate history. Accurate demographic models also serve as a statistical foundation for the identification of loci under natural selection and the design of biomedical and association studies.

Whole-genome sequencing data are high dimensional and noisy. In order to make inferences of history and biology, we rely on summary statistics of variation across the entire genome and in many sequenced individuals. One such statistic that is commonly used for demographic inference is the distribution of SNP allele frequencies in one or more populations, called the sample or allele frequency spectrum (AFS) (Marth *et al.*, 2004; Gutenkunst *et al.*, 2009; Jouganous *et al.*, 2017; Kamm *et al.*, 2017). AFS-based inference has proven to be a powerful inference approach, yet it assumes independence between SNPs and therefore ignores information contained in correlations between neighboring linked loci, which is also referred to as linkage disequilibrium (LD).

Measures of LD are also informative about demographic history, mutation, recombination, and selection. A separate class of inference methods leverage observed LD across the genome to infer local recombination rates (McVean *et al.*, 2004; Auton and McVean, 2007; Chan *et al.*, 2012; Kamm *et al.*, 2016) and demographic history (Li and Durbin, 2011; Loh *et al.*, 2013; Schiffels and Durbin, 2014; Rogers, 2014).

While two-locus statistics have been extensively studied (Hill and Robertson, 1966, 1968; Karlin and McGregor, 1968; Ohta and Kimura, 1969a,b; Golding, 1984; Ethier and Griffiths, 1990; Hudson, 2001; McVean, 2002; Song and Song, 2007), most of this work focused on a single population at equilibrium demography, precluding their application to realistic demographic scenarios. Recently, approaches for computing the full two-locus sampling distribution for a single population with non-equilibrium demography were developed via the coalescent (Kamm *et al.*, 2016) or a numerical solution to the diffusion approximation (Ragsdale and Gutenkunst, 2017), allowing for more robust inference of fine-scale recombination rates and single population demographic history. However, there remain significant limitations. Computing the full two-locus haplotype frequency spectrum is computationally expensive, hindering its application to inference problems that require a large number of function evaluations. Alternatively, computationally efficient low-order equations for specific LD statistics have been proposed (Hill and Robertson, 1968; Rogers, 2014), but these have seen limited application and only to single populations.

In this article, we show that the moment system of Hill and Robertson (1968) can be expanded to compute a large family of one-and two-locus statistics with flexible recombination, population size history, and mutation models. Additionally, we show that the system can be extended to multiple populations with continuous migration and discrete admixture events, and that low order statistics can be accurately and efficiently computed for tens of populations with complex demography.

We use this moment system together with likelihood-based optimization to infer multi-population demographic histories. We reexamine how well widely used models of human demographic history recover observed patterns of polymorphism, and find that these models underestimate LD among low frequency variants in each population, sometimes by a large amount. The inclusion of admixture from deeply diverged lineages in both Eurasian and African populations resolves these differences, and we infer an archaic lineage contributed 6 ∼*-*8% genetic ancestry in two populations in Africa. By jointly modeling a wide range of summary statistics across human populations, we can reveal important aspects of our history that are hidden from traditional analyses using individual statistics.

## Models and methods

To compute a large set of summary statistics for genetic data, we use mathematical properties of the Wright-Fisher model that are related to the look-down model of Donnelly and Kurtz (1999a,b). To illustrate this process, we first build intuition through familiar equations from population genetics and then explain how these fit within a larger hierarchy of tractable models.

In this section, we therefore begin with evolution equations for heterozygosity and the frequency spectrum, then turn to recursions for low-order LD statistics and show that the classical Hill-Robertson (Hill and Robertson, 1968) system for *D*^2^ can be extended to arbitrary moments of *D*, multiple populations, and even the full sampling distribution of two-locus haplotypes. Mathematical details and expanded discussion for each result are given in the Appendix. Throughout this article, we assume that human populations can be described, approximately, by a finite number of randomly mating populations. We also assume an infinite-sites model in the main text and describe a reversible mutation model in Appendix S1.1.3.

### Motivation: Single site statistics and the allele frequency spectrum

The most basic measure of diversity is expected heterozygosity 𝔼[*H*], or the expected number of differences between two haploid copies of the genome. Given 𝔼[*H*] at time *t*, population size *N* (*t*) and mutation rate *u*, Wright (1931) showed that enumerating all distinct ways to choose parents among two lineages leads to a recursion for 𝔼[*H*],

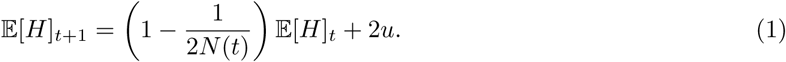

To leading order in 1*/N* and *u*, two copies of the genome are different if their parents were distinct (which has probability 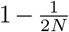) and carried different alleles (which has probability 𝔼[*H*]_*t*_), or if there was a mutation along one of two lineages (which has probability 2*u*).

Heterozygosity is a low-order statistic: we require only two copies of the genome to estimate 𝔼[*H*] genome-wide. More samples provide additional information that can be encoded in the sample AFS Φ_*n*_, the distribution of allele counts within a sample of size *n*. Specifically, Φ_*n*_(*i*) is the number (or proportion) of loci where the derived allele is observed in *i* copies out of *n* samples.

A standard forward approach to compute Φ_*n*_ involves numerically solving the partial differential equation for the distribution of allele frequencies in the full population and then sampling from this distribution for the given sample size *n* (e.g. Gutenkunst *et al.* (2009)). By enumerating mutation events and parental copying probabilities in a sample of size *n*, Jouganous *et al.* (2017) showed that Equation 1 can be generalized to a recursion for {Φ_*n*_(*i*)}_*i*=0,…,*n*_ (Figure 1). 𝔼[*H*] can be seen as a special case equal to Φ_2_(1), the *i* = 1 bin in the size *n* = 2 frequency spectrum. These recursions can also be derived as moment equations for the diffusion approximation (Evans *et al.*, 2007; Živković *et al.*, 2015; Jouganous *et al.*, 2017).

**Figure 1:**
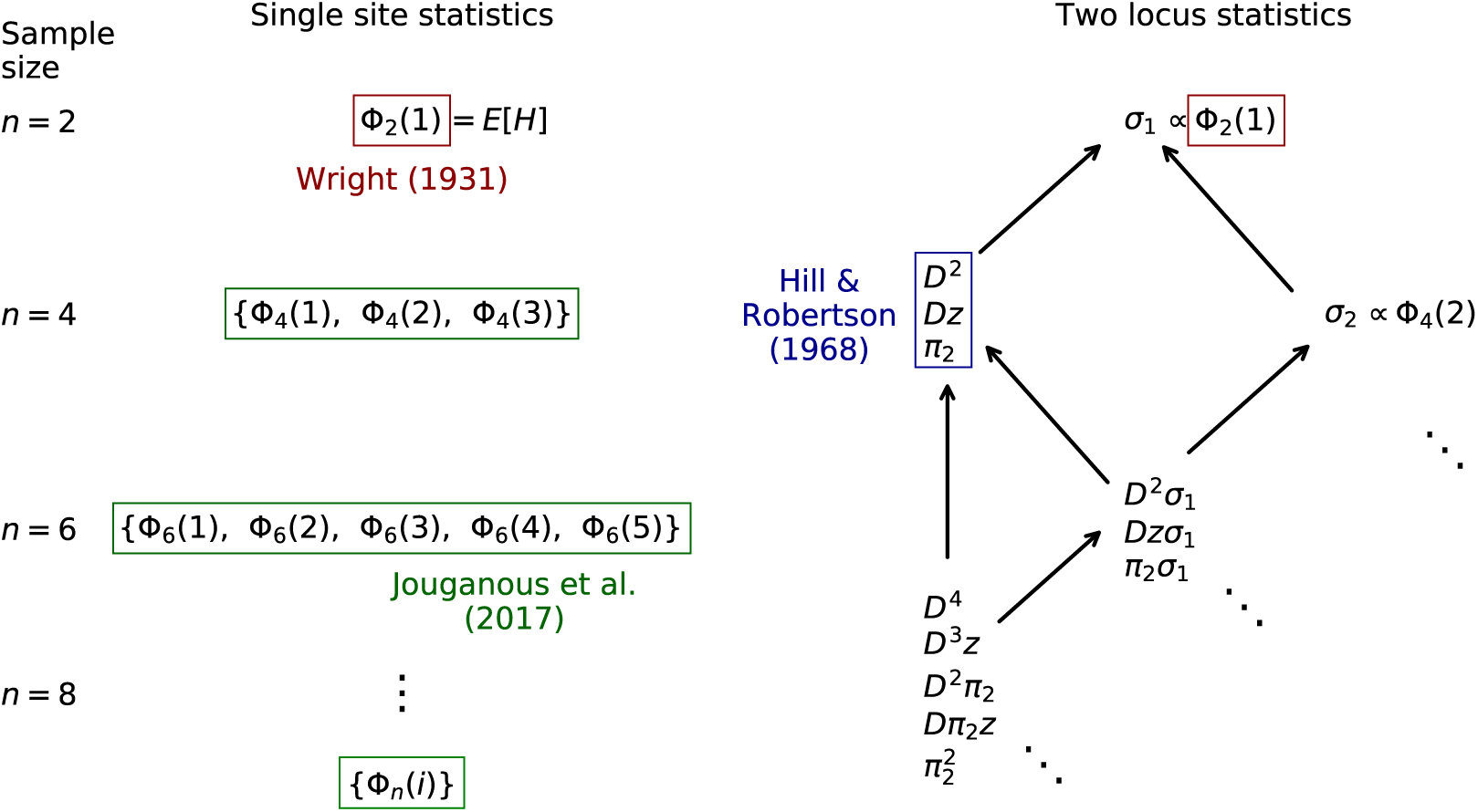
Hierarchy of even-order moments of Wright-Fisher evolution. Expected statistics under neutral Wright-Fisher evolution depend on equal or lower-order statistics in the previous generation, allowing for a hierarchy of closed recursion equations. Left: single-site statistics are represented as the entries in the size-*n* AFS, Φ_*n*_, and depend only on same-order statistics. Right: the corresponding two-locus statistics, including the Hill-Robertson system for 𝔼[*D*^2^], rely on statistics of the same or lower order. Closed recursions can be found for any given 𝔼[*D*^*m*^], leading to a sparse, linear system of ODEs. We denote *π*_2_ = *p*(1− *p*)*q*(1− *q*), *z* = (1 −2*p*)(1−2*q*), and *σ*_*i*_ = *p*^*i*^(1−*p*)^*i*^ + *q*^*i*^(1−*q*)^*i*^. Arrows indicate dependence of moments and highlighted moments indicate classical recursions. Odd-order moments are shown in Figure S1. Here we are particularly interested in such closed recursions in multi-population settings.

### Two-locus statistics

We will use this same intuition for the two-locus theory. First consider the model for two loci that each permit two alleles: alleles *A/a* at the left locus, and *B/b* at the right. There are four possible two-locus haplotypes, *AB, Ab, aB*, and *ab*, whose frequencies sum to 1 in the population. LD between two loci is measured as the covariance of their allele frequencies:

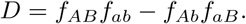

Therefore *D* can also be interpreted as the probability of drawing two lineages from the population and observing one lineage of type *AB* and the other of type *ab*, minus the probability of observing the two cross types *Ab* and *aB*. As such, 𝔼[*D*] is a two-haplotype statistic, meaning we require just two haploid copies of the genome (or a single phased diploid genome) to estimate genome-wide 𝔼[*D*], in the same way that the expected heterozygosity 𝔼[*H*] is a two-sample statistic of single-site variation.

#### Moment equations for *D* and *D*^*2*^

Enumerating possible copying, recombination, and mutation events for two lineages also leads to a well-known recursion for 𝔼[*D*] (Hill and Robertson, 1966). The possibility of sharing a common parent from the previous generation leads to the same 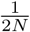 decay familiar from Equation 1. 𝔼[*D*] also decays due to recombination with rate proportional to the probability *r* of a recombination event between two loci in a *r*≪given generation. Throughout, we assume *r* 1, so that higher order terms may be ignored. For loosely linked or unlinked linked loci (*r* = 1/2), higher order terms must be considered (Hill and Robertson, 1968).

To leading order in *r, u*, and 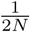 we have

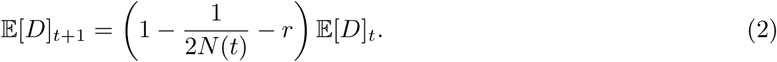

Mutation doesn’t contribute to 𝔼[*D*] because any mutation event is equally likely to contribute positively or negatively to the statistic. As a result, *D* is expected to be zero across the genome.

However, the second moment 𝔼[*D*^2^] is positive. Hill and Robertson (1968) found a recursion for a triplet of statistics including 𝔼[*D*^2^], which we write as

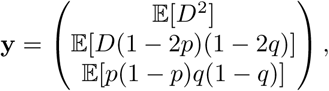

where *p* is the allele frequency of *A*, and *q* is the allele frequency of *B*. The recursion is

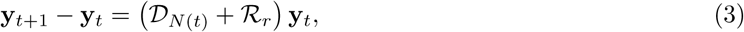

where *𝒟* and *ℛ* are matrix operators for drift and recombination, respectively. To leading order in 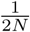 and *r*, these take the form

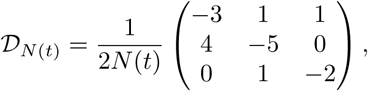

and

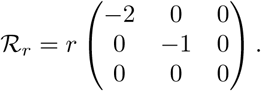

The three statistics in the Hill-Robertson system have a natural interpretation. 𝔼[*D*^2^] is the variance of *D* and has received plenty of attention over the years. The second statistic includes a term *z* = (1 *-* 2*p*)(1 *-* 2*q*) whose magnitude is largest when there are rare alleles at both loci, and which is positive when *p* and *q* both correspond to the minor allele (or both to the major allele). Thus 𝔼[*D*(1*-*2*p*)(1*-*2*q*)] = 𝔼[*Dz*] measures positive covariance among low frequency variants. Figure 2(A-C) shows that the decay of 𝔼[*D*^2^] and 𝔼[*Dz*] are sensitive to demographic history. Figure S2 shows how the bulk of the *Dz* statistic is contributed by pairs of variants where the rarest allele has frequency between 2 and 20%, while common variants comprises the bulk of *D*^2^.

**Figure 2:**
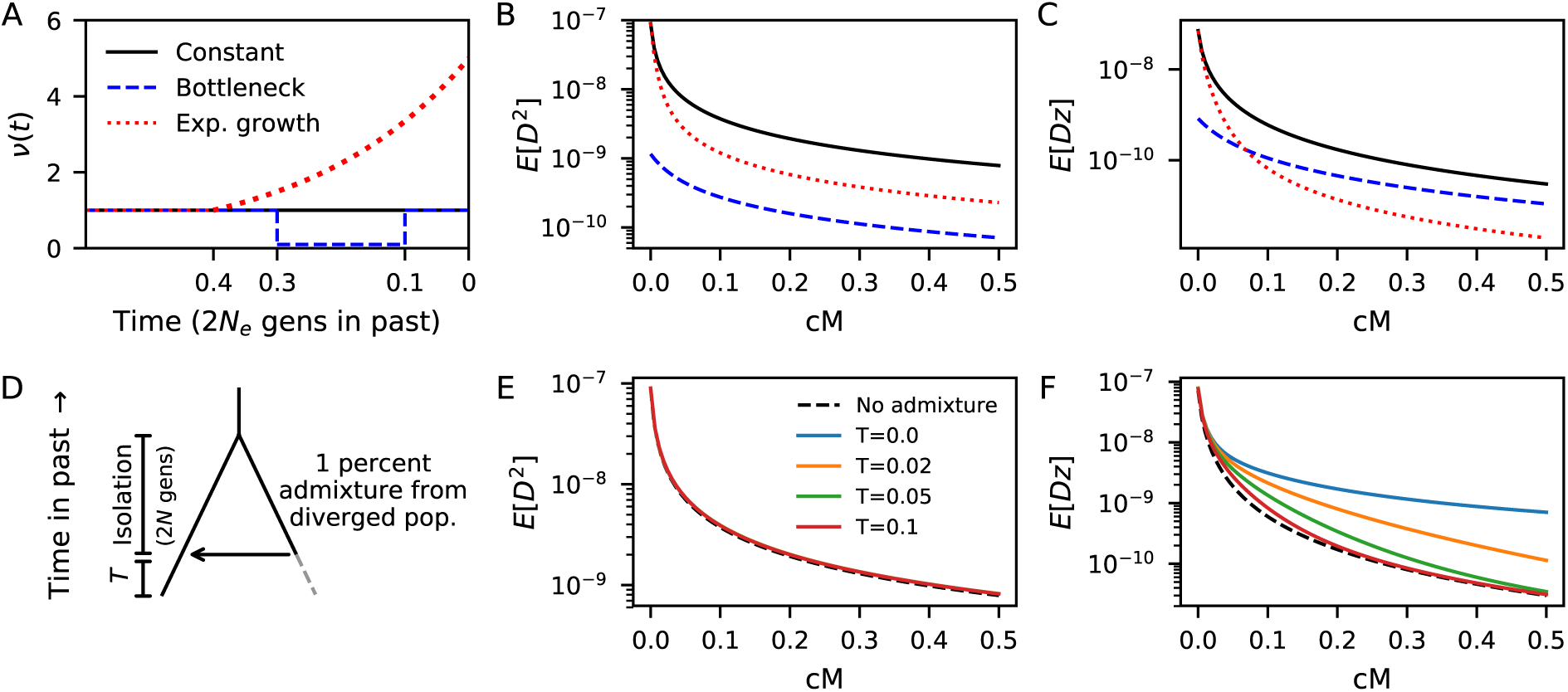
LD curves are sensitive to demography. Demographic histories shown in (A) affect statistics in the Hill-Robertson system and their dependance on recombination distance (B-C). Both the amplitude and shape of the LD curves differ between demographic models for (B) 𝔼[*D*^2^] and (C) 𝔼[*Dz*] = 𝔼[*D*(1−2*p*)(1 − 2*q*)]. (D) To illustrate the effect of admixture on LD curves, we consider two populations in isolation for 2*N* generations, followed by an admixture event where the focal population receives 1% of lineages from the diverged population. (E) 𝔼[*D*^2^] curves are largely unaffected by this low level of admixture. (F) However, 𝔼[*Dz*] is immediately and strongly elevated following admixture, and remains significantly elevated for prolonged time *T* (in units of 2*N* generations) since the admixture event.

𝔼[*π*_2_] = 𝔼[*p*(1*-p*)*q*(1*-q*)] is the joint heterozygosity across pairs of SNPs. If we sample four haplotypes from the population, this is proportional to the probability that the first pair differ at the left locus, and the second pair differ at the right locus.

The applications in this article focus on generalizing the Hill-Robertson equations to multi-population settings. However, we first outline generalizations to high-order moments and non-neutral evolution, leaving theoretical developments and simulations to the Appendix.

#### Generalizing to higher moments of *D*

The existence of tractable higher-order moment equations for one-locus statistics (Jouganous *et al.*, 2017) suggests the existence of a similar high-order system for two-locus statistics. Higher moments of *D* provide additional information about the distribution of two-locus haplotypes. Appendix S1.1 shows that the Hill-Robertson system can be extended to compute any moment of *D*, and presents recursions for those systems of arbitrary order *D*^*m*^ that closes under drift, recombination, and mutation.

This family of recursion equations takes a form similar to the *D*^2^ system: the evolution of 𝔼[*D*^*m*^] requires 𝔼[*D*^*m-*1^*z*] and 𝔼[*D*^*m-*2^*π*_2_], with each of those terms depending on additional terms of the same order and smaller orders (Figure 1). For any order *m*, Appendix S1.4.1 shows the system closes and forms a hierarchy of moment equations, in that the *D*^*m*^ recursion contains the *D*^*m-*2^ system, which itself contains the *D*^*m-*4^ system, and so on (Figure S1). Just as the Wright equation for heterozygosity generalizes naturally to equations for the more informative distribution of allele frequency (Jouganous *et al.*, 2017), the Hill and Robertson equations for 𝔼[*D*] and 𝔼[*D*^2^] generalize to informative higher-order LD statistics.

#### Generalizing to arbitrary two-locus haplotype distribution

Given the analogy between the frequency spectrum and the Hill-Robertson equations, it is natural to study the connection between the moment equations for 𝔼[*D*^*n*^] and the evolution of the two-locus haplotype frequency distribution Ψ_*n*_(*f*_*AB*_, *f*_*Ab*_, *f*_*aB*_, *f*_*ab*_).

While classical approaches for computing Ψ_*n*_ (Golding, 1984; Hudson, 2001) were limited to neutrality and steady-state demography, recent coalescent and diffusion developments allow for Ψ_*n*_ to be computed under non-equilibrium demography and selection (Kamm *et al.*, 2016; Ragsdale and Gutenkunst, 2017). These approaches are computationally expensive and limited to one population, as Ψ_*n*_ has size 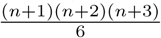 and the *P*-population distribution grows asymptotically as *n*^3*P*^.

Generalizing the approach of Jouganous *et al.* (2017), we can write a recursion equation on the entries of Ψ_*n*_ under drift, mutation, recombination, and selection at one or both loci (Appendix S1.3). As expected, this recursion does not close under selection: to find Ψ_*n*_ at time *t* + 1, we require Ψ_*n*+1_ and Ψ_*n*+2_ at time *t*. It also does not close under recombination, requiring a closure approximation. Using the same closure strategy for selection and recombination, however, we can approximate the entries of Ψ_*n*+1_ and Ψ_*n*+2_ as linear combinations of entries in Ψ_*n*_ and obtain a closed equation. This approach provides accurate approximation for moderate *n* under recombination and selection (Appendix S1.3.5) that represent a 10 to 100-fold speedup over the numerical PDE implementation in Ragsdale and Gutenkunst (2017) (Table S1). However, closure is inaccurate for small *n*.

By contrast to the full two-locus model, equations for moments of *D* close under recombination because the symmetric combination of haplotype frequencies that define *D* ensures the cancellation of higher-order terms (Appendix S1.1.2). This makes the moments of *D* particularly suitable for rapid computation of low-order statistics over a large number of populations.

The Hill-Robertson system does not close, however, if one or both loci are under selection. Appendix S1.1.4 considers a model where one of the two loci is under additive selection. We derive recursion equations for terms in the 𝔼[*D*^2^] system and describe the moment hierarchy and a closure approximation, though we leave its development to future work. In the following we focus on neutral evolution.

### Multiple populations

While a large body of work exists for computing expected LD in a single population, little progress has been made toward extending these models to multiple populations. Forward equations for the full two-locus sampling distribution become computationally intractable beyond just a single population, even with the moment-based approach described above. Here, we extend the Hill-Robertson system to any number of populations, allowing for population splits, admixture, and continuous migration.

#### Motivation: Heterozygosity across populations

To motivate our derivation of the multi-population Hill-Robertson system and provide intuition, we begin with a model for heterozygosity across populations with migration. With two populations we consider the cross-population heterozygosity, 𝔼[*H*_12_] =𝔼[*p*_1_(1*-p*_2_)]+𝔼[*p*_2_(1*-p*_1_)], where *p*_*i*_ and *q*_*i*_ are allele frequencies at the left and right loci, respectively, in population *i*. This is the probability that two lineages, one drawn from each population, differ by state. At the time of split between populations 1 and 2, 𝔼[*H*_1_] = 𝔼[*H*_2_] = 𝔼[*H*_12_]. Because coalescence between lineages in different populations is unlikely, 𝔼[*H*_12_] is not directly affected by drift. In the absence of migration and under the infinite-sites assumption used here, this statistic increases linearly with the mutation rate over time (Figure S3).

With migration, the evolution of 𝔼[*H*_12_] also depends on 𝔼[*H*_1_] and 𝔼[*H*_2_]. We define the migration rate *m*_12_ to be the probability that a lineage in population 2 has its parent in population 1. Assuming *m*_*ij*_≪1, the probability that both lineages in 𝔼[*H*_12_] come from population 1 is *m*_12_ (to leading order), in which case 𝔼[*H*_12_]_*t*+1_ is equal to 𝔼[*H*_1_]_*t*_, and the probability that both come from population 2 is *m*_21_. Then to leading order in *m*_*ij*_, we have

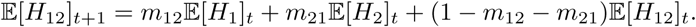

Similar intuition leads to recursions for 𝔼[*H*_1_] and 𝔼[*H*_2_] under migration, and this system easily extends to more than two populations.

#### The Hill-Robertson system with migration

We take the same approach to determine transition probabilities in the multi-population Hill-Robertson system. Suppose that at some time, a population splits into two populations. At the time of the split, expected two-locus statistics (*D*^2^, *Dz, π*_2_) in each population are each equal to those in the parental population at the time of split (Appendix S1.2.1). Additionally, the covariance of *D* between the two populations, 𝔼[*D*_1_*D*_2_], is initially equal to 𝔼[*D*^2^] in the parental population. In the absence of migration, Hill-Robertson statistics in each population evolve according to Equation 3, and

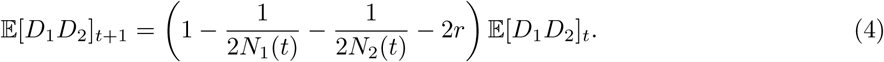

With migration, additional moments are needed to obtain a closed system. These additional terms take the same general form as the original terms in the Hill-Robertson system, but include cross-population statistics, analogous to *H*_12_ in the heterozygosity model with migration. Again using **y** to denote bases of Hill-Robertson moments, this basis is

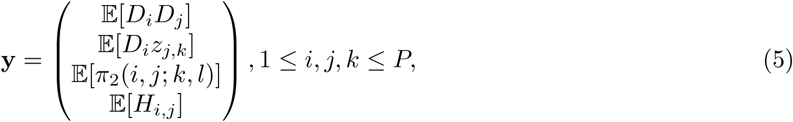

where *P* is the number of populations, and we slightly abuse notation so that *D*_*i*_*D*_*j*_ stands in for all index permutations (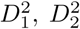, and *D*_1_*D*_2_ in the two-populations case). We derive transition probabilities under continuous migration in Appendix S1.2.2 leading to the closed recursion,

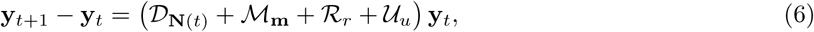

where 𝒟, ℳ, ℛ and 𝒰 are sparse matrices for drift, migration, recombination and mutation that depend on the number of populations, population sizes **N**(*t*), and migration rates **m**.

#### Admixture

Patterns of LD are sensitive to migration and admixture events, and low order LD statistics are commonly used to infer the parameters of admixture events (Moorjani *et al.*, 2011; Loh *et al.*, 2013). A well-known result (e.g., example 2.7 in Cavalli-Sforza and Bodmer (1971)) is that *D* in an admixed population can be nonzero even when *D* is zero in both parental populations if allele frequencies differ between the two parental populations. This is seen by enumerating all possible combinations of haplotype sampling when a fraction *f* of lineages were contributed by population 1, and 1 − *f* by population 2 (Appendix S1.2.3). More generally, immediately following the admixture event, the expectation 𝔼[*D*_adm_] in the admixed population is

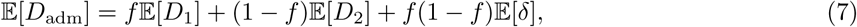

where *δ* = (*p*_1_− *p*_2_)(*q*_1_−*q*_2_) (Nei and Li, 1973).

To integrate the multi-population *D*^2^ system after an admixture event, we require 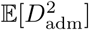 and other second order terms in the basis (5) involving the admixed population. Using the same enumeration approach as for Equation 7, the expectation immediately following the admixture event is

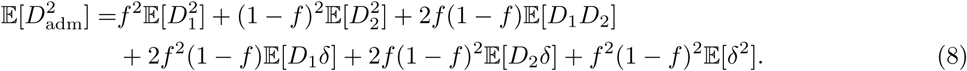

Each other required term can be found in a similar manner (Appendix S1.2.3). In this way, the set of moments may be expanded to include the admixed population and integrated forward in time using Equation 6.

### Numerical implementation

We rescale time by 2*N*_ref_ generations (*N*_ref_ is an arbitrary reference population size, often the ancestral population size), so that the recursion can be approximated as a differential equation

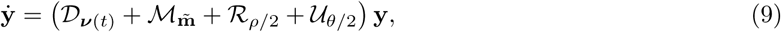

where ν are the relative population sizes at time *t* (*ν*_*i*_(*t*) = *N*_*i*_(*t*)*/N*_ref_), 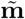 are the population size-scaled migration rates 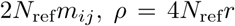, and 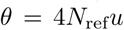. Each matrix is sparse, and this equation can be solved efficiently using a standard Crank-Nicolson integration scheme. Our implementation allows users to define general models with standard demographic events (migrations, splits and mergers, size changes, etc.) similar to the *∂*a*∂*i/moments interface (Gutenkunst *et al.*, 2009; Jouganous *et al.*, 2017). A single evaluation of the four-population model shown in Figure S4 can be computed in roughly 0.1 second. We packaged our method with `moments` (Jouganous *et al.*, 2017) as `moments.LD`, a python module that computes expected statistics and performs likelihood-based inference from observed data (described below), available at bitbucket.org/simongravel/moments.

#### Validation

We validated our numerical implementation and estimation of statistics from simulated genomes using `msprime` (Kelleher *et al.*, 2016). Expectations for low-order statistics match closely with coalescent simulations. For example, Figure S4 shows the agreement for a four population model with non-constant demography, continuous migration, and an admixture event, for which we computed expectations using `moments.LD` that matched estimates from `msprime`. While approximating expectations from `msprime` required the time-consuming running and parsing of many simulations, expectations from `moments.LD` were computed in seconds on a personal computer.

## Data and inference

### Genotype data

Computing *D* using the standard definition requires phased haplotype data (Appendix S1.5). However, most currently available whole genome sequence data is unphased, so that we must rely on two-locus statistics based on observed genotype counts instead of haplotype counts. One could estimate haplotype statistics using the Weir (1979) estimator

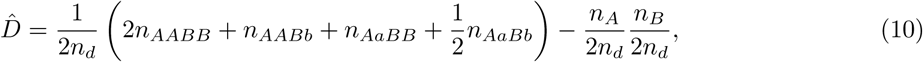

where *n*_*A*_ is the count of *A* at the left locus, *n*_*B*_ the count of *B* at the right locus, *n*_*d*_ the number of diploid individuals in the sample, and {*n*_*AABB*_, *n*_*AABb*_, …} the counts of each observed genotype. However, the Weir estimator for 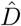 is biased. Fortunately, we can simply treat the Weir estimator *D* as a statistic and obtain an unbiased prediction for its expectation (Appendix S1.7.3). Even though 𝔼[*D*^*n*^] can be estimated from 2*n* phased haplotypes, more samples are required to accurately estimate LD for a given pair of SNPs. However, as we are interested in genome-wide averages of 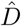 and other LD statistics, even when individual estimates are noisy, by averaging over a very large number of pairs of SNPs we can accurately estimate LD from relatively few diploid genomes.

### 1000 Genome Project data

We computed statistics from intergenic data in the Phase 3 1000 Genomes Project data (1000 Genomes Project Consortium *et al.*, 2015). The non-coding regions of the 1000 Genomes data is low coverage, which can lead to significant underestimation of low frequency variant counts, which distorts the frequency spectrum and can lead to biases in AFS-based demographic inference (Gravel *et al.*, 2011). However, low-order statistics in the Hill-Robertson system are robust to low coverage data in a large enough sample size (Figure S6), so that low coverage data are well suited for inference from LD statistics (see also Rogers (2014)).

To avoid possible confounding due to variable mutation rate across the genome, we calculated and compared statistics normalized by *π*_2_, the joint heterozygosity 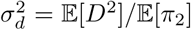, as in Rogers (2014). All figures showing 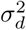-type statistics are normalized using *π*_2_(YRI), the joint heterozygosity in the Yoruba from Ibidan, Nigeria (YRI). This normalization removes all dependence of the statistics on the overall mutation rate, so that estimates of split times and population sizes are calibrated by the recombination rate per generation instead of the mutation rate (Ragsdale and Gutenkunst, 2017). This is convenient given that genome-wide estimates of the recombination rate tend to be more consistent across experimental approaches than estimates of the mutation rate.

We considered all pairs of intergenic SNPs with 10^*-*5^≤ *r* ≤ 2×10^*-*3^ using the African-American recombination map estimated by Hinch *et al.* (2011) using ancestry switch-points. The lower bound was chosen to further reduce the potential effect of short-range correlations of mutation rates, clustered mutations, experimental error, and low resolution of the recombination map at very short distances.

### Likelihood-based inference on LD-curves

To compare observed LD statistics in the data to model predictions, and thus to evaluate the fit of the model to data, we used a likelihood approach. We binned pairs of SNPs based on the recombination distance separating them (Appendix S1.7.2). Bins were defined by bin edges {*r*_0_, *r*_1_, *…, r*_*n*_}, roughly logarithmically spaced. The model is defined by the set of demographic parameters Θ. We included the ancestral *N*_ref_ as a parameter to be fit, which we also use to scale recombination bins as *ρ*_*i*_ = 4*N*_ref_*r*_*i*_.

For a given recombination bin (*ρ*_*i*_, *ρ*_*i*+1_], we computed statistics and normalize by *π*_2_ in one population (we used *π*_2_(YRI)), and denote this set of normalized statistics 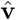. We computed expectations for normalized statistics from the model, **M**_*i*_, and then estimated the likelihood as

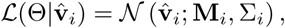

taking the probability of observing data 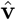 to be normally distributed with mean **M** and covariance matrix σ (the normal distribution assumption is validated in Figure S5).

We estimated σ directly from the data by constructing bootstrap replicates from sampled subregions of the genome with replacement. This has the advantage of accounting for the covariance of statistics in our basis, as well as non-independence between distinct neighboring or overlapping pairs of SNPs. To compute the composite likelihood across *ρ* bins, we simply took the product of likelihoods over values of recombination bins indexed by *i*, so that

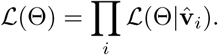

To compute confidence intervals on parameters, we used the approach proposed by Coffman *et al.* (2016), which adjusts uncertainty estimates to account for non-independence between recombination bins and neighboring pairs of SNPs.

## Results

### Human expansion models underestimate LD between low frequency variants

The demographic model for human out-of-Africa (OOA) expansion proposed and inferred by Gutenkunst *et al.* (2009) has been widely used for subsequent simulation studies, and parameter estimates have been refined as more data became available (Gravel *et al.*, 2011; Tennessen *et al.*, 2012; Jouganous *et al.*, 2017). These models have typically been fit to the single-locus joint AFS, with Yoruba of Ibidan, Nigeria (YRI), Utah residents of Western European ancestry (CEU), and Han Chinese from Beijing (CHB) as representative panels. Gutenkunst *et al.* (2009) verified that the observed decay of *r*^2^ was consistent with simulations under their inferred model.

We first asked if the OOA model (Figure 3(A)) is able to capture observed patterns of LD within and between these three populations. When fitting to all statistics in the multi-population basis, parameters diverged to infinite values, suggesting that the model is mis-specified. In particular, this model was unable to describe observed *Dz* statistics, with *Dz*-curves from the model drastically underestimating observations.

**Figure 3:**
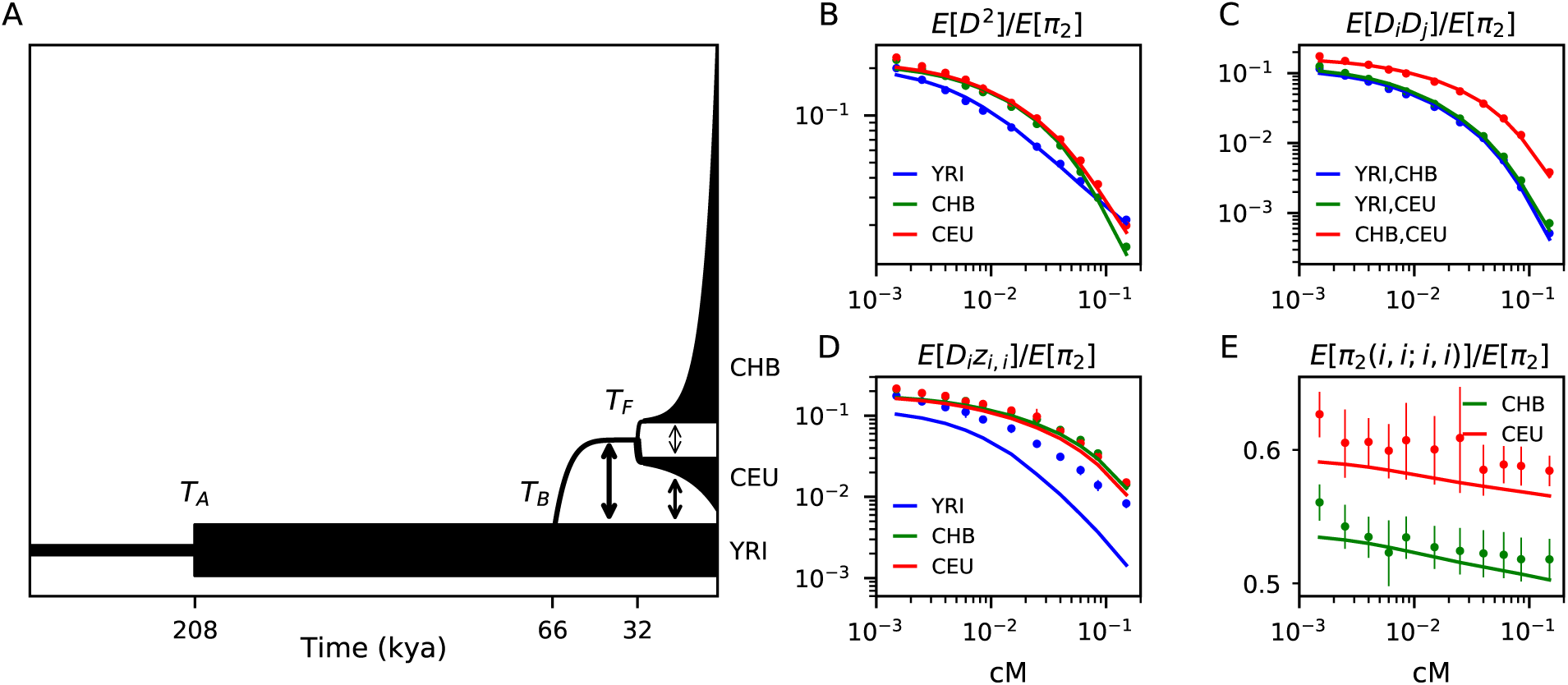
Standard out-of-Africa model underestimates LD among low frequency variants. (A) We fit the 13-parameter model proposed by Gutenkunst *et al.* (2009) to statistics in the two-locus, multi-population Hill-Robertson system. The remaining 35 statistics from the Hill-Robertson basis used in the fit are shown in Figure S7 and residuals are shown in Figure S8. Best fit values for labeled parameters are given in Table 1. Most statistics were accurately predicted by this model, including (B) the decays of 𝔼[*D*^2^] in each population, (C) the decay of the covariance of *D* between populations, and (E) the joint heterozygosity 𝔼[*π*_2_(*i*)]. (D) However, 𝔼[*D*_*i*_(1− 2*p*_*i*_)(1− 2*q*_*i*_)] was fit poorly by this model, and we were unable to find a three-population model that recovered these observed statistics, including with additional periods of growth, recent admixture between modern human populations, or substructure within modern populations. Error bars represent bootstrapped 95% confidence intervals on the statistic estimate.

We refit the OOA model without including *Dz* statistics, and we inferred best-fit parameters that generally align with estimates using the joint AFS (Table 1, left, and Figure 3). This model underestimated observed *Dz* in each population, especially in the YRI population (Figure 3(D)). Using AFS-inferred parameters from previous studies led to qualitatively similar results.

**Table 1:** Inferred parameters for OOA models. Two models for the out-of-Africa expansion. We fit the commonly used 13-parameter model to the multi-population Hill-Robertson statistics (left). The best fit parameters shown here were fit to the set of statistics without the 𝔼[*Dz*] terms, because the inclusion of those terms led to runaway parameter behavior in the optimization. This is often a sign of model mis-specification. On the right, the same 13-parameter model is augmented by the inclusion of two deeply diverged branches, putatively Neanderthal and an unknown lineage within Africa. We inferred that these branches split from the branch leading to modern humans roughly 460−650 kya, and contributed migrants until quite recently (∼ 19 kya). Times reported here assume a generation time of 29 years and are calibrated by the recombination (rather than mutation) rate. Confidence intervals were computed using the Godambe information matrix on bootstrap replicates of the data (Coffman *et al.*, 2016).

| Parameter | Model | OOA (fit w/o $Dz$) | | archaic admixture | |
| --- | --- | --- | --- | --- | --- |
|  |  | Estimates | 95% CI | Estimates | 95% CI |
| $N_0$ | | 2360 | 2190 – 2530 | 3600 | 2380 – 3920 |
| $N_{YRI}$ | | 13030 | 12200 – 13900 | 13900 | 12800 – 15000 |
| $N_B$ | | 1080 | 810 – 1350 | 880 | 670 – 1090 |
| $N_{CEU0}$ | | 1450 | 980 – 1920 | 2300 | 1810 – 2790 |
| $r_{CEU}(\%)$ | | 0.202 | 0.184 – 0.217 | 0.125 | 0.113 – 0.135 |
| $N_{CHB0}$ | | 410 | 340 – 480 | 650 | 540 – 750 |
| $r_{CHB}(\%)$ | | 0.498 | 0.445 – 0.531 | 0.372 | 0.333 – 0.398 |
| $m_{AF-B}(\times 10^{-5})$ | | 51.5 | 41.1 – 61.8 | 52.2 | 41.7 – 62.6 |
| $m_{YRI-CEU}(\times 10^{-5})$ | | 1.72 | 0.54 – 2.9 | 2.48 | 1.84 – 3.13 |
| $m_{YRI-CHB}(\times 10^{-5})$ | | 0 | — | 0 | — |
| $m_{CEU-CHB}(\times 10^{-5})$ | | 15.3 | 11.5 – 19.1 | 11.3 | 8.70 – 13.8 |
| $T_{AF}$ (kya) | | 208 | 196 – 220 | 300 | 277 – 323 |
| $T_{OOA}$ (kya) | | 65.7 | 52.4 – 79.0 | 60.7 | 50.3 – 64.2 |
| $T_{CEU-CHB}$ (kya) | | 31.9 | 28.8 – 35.0 | 36.0 | 32.3 – 39.6 |
| $T_{Arch. Af. split}$ (kya) | | | | 499 | 460 – 538 |
| $T_{Arch. Af. mig.}$ (kya) | | | | 125 | 89.2 – 160 |
| $m_{AF-Arch. Af.}(\times 10^{-5})$ | | | | 1.98 | 1.15 – 2.82 |
| $T_{Nean. split}$ (kya) | | | | 559 | 470 – 648 |
| $m_{OOA-Nean}(\times 10^{-5})$ | | | | 0.825 | 0.379 – 1.27 |
| $T_{Arch. adm. end}$ (kya) | | | | 18.7 | 15.1 – 22.4 |

The Gutenkunst model is a vast oversimplification of human evolutionary history, so its failure to account for *Dz* is not all that surprising. However, given the good agreement of the model to both allele frequencies and *r*^2^ decay (Gutenkunst *et al.*, 2009), we did not expect such a large discrepancy. Having ruled out low coverage and spatial correlations in the mutation rate as explaining factors, our next hypothesis was a more complex demographic history. We generalized the Gutenkunst model with a number of additional parameters accounting for recent events, including size changes in the YRI population, recent mixture between populations, and substructure within each continental population. None of these modifications provided satisfactory fit to the data and some did not converge to biologically realistic parameters.

### Inference of archaic admixture

𝔼[*Dz*] is a measure of positive covariance between low-frequency alleles (Figure S2). We therefore expect this statistic to be sensitive to the presence of rare, deep-coalescing lineages within the population, as those lineages will contribute haplotypes with a large number of tightly linked low frequency variants (see Discussion below).

Given prior genetic evidence for archaic admixture in Eurasia and Africa (reviewed in Wall and Brandt (2016)), we proposed a model that includes two deeply diverged human branches, with one branch mixing with Eurasian ancestors beginning at the OOA event, and the second one mixing with the ancestors of the Yoruba population over a time period that could include the OOA event. In this scenario, this second branch could also contribute to Eurasians through admixture prior to the OOA event (Figure 4(A)). Many human lineages coexisted on the African continent, possibly until quite recently (Rightmire, 2009; Harvati *et al.*, 2011; Berger *et al.*, 2017), and genetic evidence points to a history of archaic admixture or deep structure across many modern African populations (Hammer *et al.*, 2011; Lachance *et al.*, 2012; Hsieh *et al.*, 2016; Skoglund *et al.*, 2017; Durvasula and Sankararaman, 2018; Hey *et al.*, 2018). It is likely that modern humans have met and mixed with diverged lineages many times through history, rather than receiving just a single pulse of migrants (Browning *et al.*, 2018; Villanea and Schraiber, 2019). We chose to model the mixing of archaic and modern human branches as continuous and symmetric (Kuhlwilm *et al.*, 2016), parameterizing the migration rate between these branches and the times that migration began and ended.

**Figure 4:**
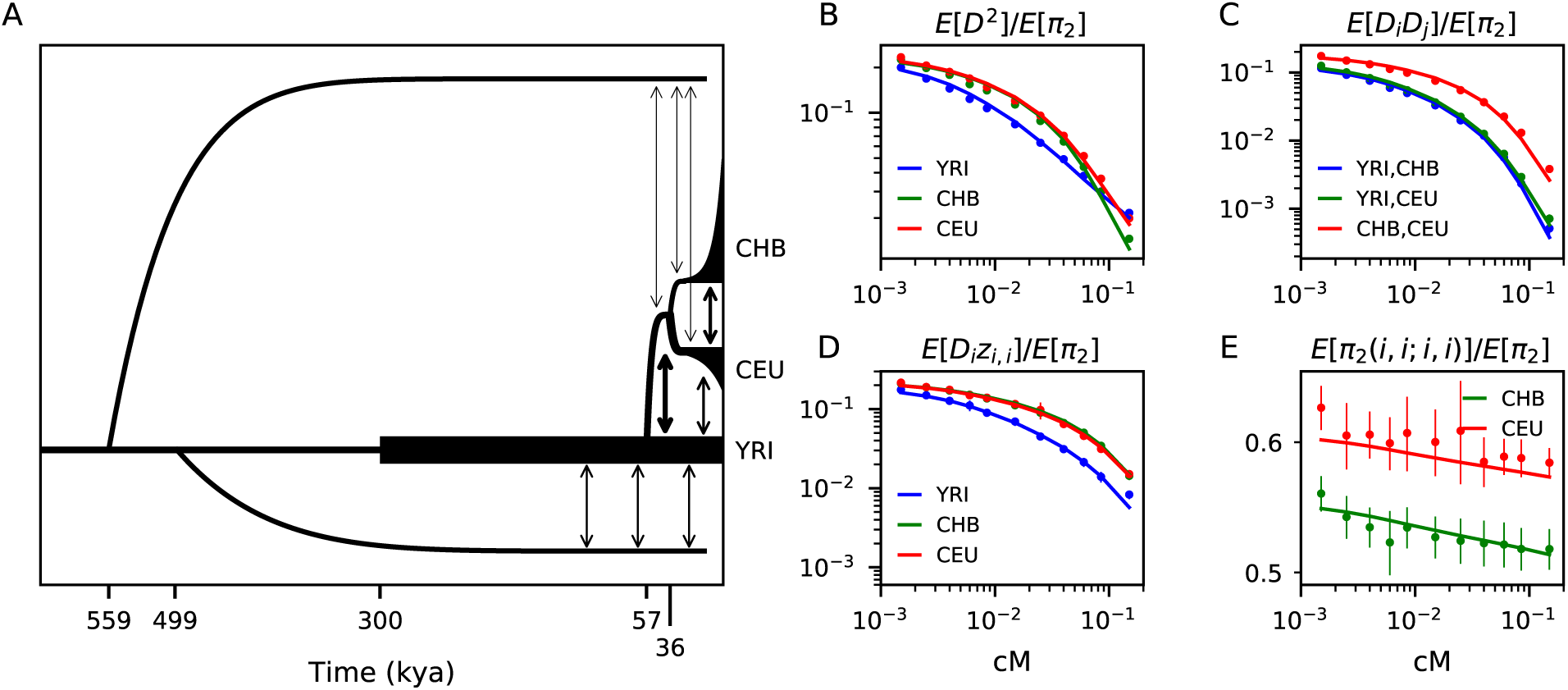
Inferred OOA model with archaic admixture. (A) We fit a model for out-of-Africa expansion related to the standard model in Figure 3(A). Demographic events for the three modern human populations are parameterized as above, but we also include two branches with deep split from the ancestral population to modern humans. A putatively Neanderthal branch that remains isolated until the Eurasian split from YRI, and a deep branch within Africa that is allowed to be isolated for some time before continuously exchanging migrants with the common ancestral branch and the YRI branch. (B-E) This model fits the data much better than the model without archaic admixture, and especially for the *Dz* statistics (D). Fits to 35 more curves and statistics are shown in Figure S7 and residuals are shown in Figure S8. The migration rates inferred between the diverged African branch and YRI provides an estimate of ∼ 7.5% contribution.

We considered two topologies for the archaic branches: 1) both branches split independently from that leading to modern humans (Figure 4(A)), and 2) one branch split from the modern human branch, which some time later split into the two populations (Figure S9(A)). Both models fit the data well with little statistical evidence to discriminate between these two models (Figures 4(B-E) and S8). The difference in log-likelihood between the two models was Δ*LL <* 1, as opposed to Δ*LL* = 1, 730 between models with and without archaic admixture. Δ*LL* between the best fit model with archaic admixture and the fully saturated model (using observations as expectations) was 767. Consistent among the inferred models was the age of the split between diverged and modern human branches within Africa at ∼ 500 kya, though uncertainty remains with regard to the relationship between archaic human lineages in Africa and Eurasia. The sequencing of archaic genomes within Africa would clearly be helpful in resolving these topologies.

We inferred an archaic population to have contributed measurably to Eurasian populations. This branch (putatively Eurasian Neanderthal) split from the branch leading to modern humans 470-650 thousand years ago (kya), which contributed 1.2±0.6% ancestry in modern CEU and CHB populations after the out-of-Africa split. This range of divergence dates from our maximum-likelihood model overlaps with previous estimates of the time of divergence between Neanderthals and human populations, estimated at 550 *–* 765 kya (Prüfer *et al.*, 2014). The diverged African branch split from the ancestors of modern humans 460-540 kya and contributed to both the pre-OOA human branch and the lineage leading to YRI. This admixture began between 90 − 160 kya, well before the estimated split between Eurasian and the YRI lineages, so that this archaic branch also contributed to the ancestors of Eurasian populations. We estimated 4.7 − 9.2% ancestry contribution from this unknown population to YRI, and 1.9−6.6% contribution to CEU and CHB.

We chose a separate population trio to validate our inference and compare levels of archaic admixture with different representative populations. This second trio consisted of the Luhya in Webuye, Kenya (LWK), Kinh in Ho Chi Minh City, Vietnam (KHV), and British in England and Scotland (GBR). We inferred the KHV and GBR populations to have experienced comparable levels of migration from the putatively Neanderthal branch. However, the LWK population exhibited lower levels of admixture (∼6%) in comparison to YRI, possibly suggesting population differences in archaic admixture events within the African continent (Table S3).

## Discussion

### Multi-population two-locus diversity statistics

The application presented here relied on the four-haplotype statistics (*D*^2^, *Dz, π*_2_). Studying these low-order multi-population statistics in a likelihood framework allowed us to infer a demographic model with archaic admixture, even without reference genomes from those diverged populations. We have also shown that higher order statistics may be computed through this same framework. Extending higher order two-locus moment systems to multiple populations would potentially provide further information about demography, particularly for past encounters with archaic lineages.

#### Relation to other statistics

There are many approaches for computing expected statistics for diversity under a wide range of scenarios. Single-site statistics, which include expected heterozygosity and the AFS, may be computed efficiently using forward- or reverse-time approaches. Beyond the classical recursions for 𝔼[*D*] and 𝔼[*D*^2^] (Hill and Robertson, 1968; Rogers, 2014), two-locus statistics are difficult to compute for non-equilibrium, multi-population demographic models. Sved (2009) proposed an IBD based recursion to compute 𝔼[*r*^2^] across subdivided populations, but its accuracy and interpretation remain debated (Rogers, 2014).

The moments-based approach presented here generalizes the recursion for the single-site AFS presented in (Jouganous *et al.*, 2017). The moments system includes all heterozygosity statistics, so we recover expected *F* -statistics under arbitrary demography, which are commonly used to test for admixture (Reich *et al.*, 2009; Patterson *et al.*, 2012; Peter, 2016). Long-range patterns of elevated LD in putatively admixed populations are used to infer the timing of admixture events and relative contributions of parental populations (Moorjani *et al.*, 2011; Loh *et al.*, 2013). These approaches rely on the recursion for 𝔼[*D*] after admixture events that is used here (Equations 2 and 7). Thus the generalized Hill-Robertson system is sensitive to ancient admixture, but also captures statistics used to identify recent admixture history, with fewer assumptions about early history.

Plagnol and Wall (2006) and Wall *et al.* (2009) introduced a statistic, *S*\*, specifically designed to scan for introgressed haplotypes without having sequence data from the diverged population. *S*\* uses an ad-hoc score to identify SNPs that likely arose on haplotypes contributed from a deeply diverged population, and is estimated through simulation. These SNPs will tend to be rare and in high LD, and therefore also contribute to *Dz* (Figure 2(D-F)). Thus even a small amount of archaic admixture will significantly elevate 𝔼[*Dz*] compared to that in an unadmixed population, and *Dz* itself could be used as an ad-hoc statistic similar to *S*\*. Given its conceptual relationship to *S*\*, it may not be so surprising that this previously overlooked statistic is particularly well suited for model-based inference of archaic admixture.

#### Caveats

Like many inference approaches in population genetics, we approximate human history using discrete, randomly mating populations with size and migration histories described by relatively few parameters. History is much more complex than this. Thus statistical uncertainties estimated using bootstrap analysis masks much larger, systematic errors due to model misspecification. In particular, some choices we made in modeling archaic admixture are certainly oversimplified, such as the assumption of symmetric and constant migration rates during the period of contact between archaic and modern humans.

Variability in fine-scale recombination rates between populations and over time contributes another source of systematic error. While large-scale recombination rates are generally better understood than the mutation rate in humans [for which current estimates vary over a factor of two (Scally, 2016)], recombination rates can vary at short distances. Spence and Song (2019) showed that recombination maps are highly concordant across populations represented in 1000 Genomes Project Consortium *et al.* (2015), although this correlation surely decreases at shorter distances. We filtered out pairs of mutations at very close distances (less than roughly 1kb) to reduce potential biases due to very fine scale variation. We therefore do not expect variation in recombination rate among human populations to explain the large differences in *Dz* compared to the Gutenkunst *et al.* model. However, the effect of population-specific recombination maps may play a role when considering finer-scale patterns and data from deeply diverged populations such as the Neanderthal.

Finally, our model and inferences assumed that mutations are evolving neutrally. We chose to analyze SNPs in intergenic regions and excluded genic and intronic regions in an effort to reduce biases due to selection acting on mutations included in the analysis or nearby selected regions, although some intergenic regions are expected to be affected by selection or biased gene conversion. While outside the scope of this study, a more detailed characterization of the effects of linked selection on Hill-Robertson statistics is warranted.

## Conclusion

We described an infinite hierarchy of multi-locus summaries of genomic diversity that are easy to compute under arbitrary, multi-population demographies. Some of these statistics are familiar, including expected heterozygosity, *F* -statistics, and LD decay, while others have been largely unexplored in multi-population models, such as the degree of LD between low frequency alleles (*Dz*) and the joint heterozygosity across sites and populations (*π*_2_). The one-population *Dz* statistic, in particular, has an interesting history, as it has come up in early work as a mathematical stepping-stone on the way to computing *D*^2^ (Hill and Robertson, 1968), but was, to our knowledge, never used in data analysis. As it happens, this ‘ghost’ statistic provides a unique window into human history.

Using this set of summary statistics, we explored a commonly used model of human demographic history derived from single-site AFS and validated using LD decay curves. While many statistics under this model fit the data well, the model dramatically underestimates levels of LD among rare alleles. Modeling archaic admixture worldwide resolved this discrepancy. We recovered the signal of Neanderthal admixture in Eurasian populations, and found evidence for substantial and long-lasting admixture from a deeply diverged lineage in two African populations that is consistent with evidence from previous studies (Plagnol and Wall, 2006; Skoglund *et al.*, 2017; Hey *et al.*, 2018; Durvasula and Sankararaman, 2018).

This model deserve a more thorough investigation, including data from ancient humans and additional contemporary African populations. We leave this to future work for three reasons. First, proposing a detailed multi-population model of evolution in Africa will require carefully incorporating anthropological and archaeological evidence, which is a substantial endeavor. Second, the inclusion of two-locus statistics from ancient genomes will require vetting possible biases associated with ancient DNA sequencing, although we see no problem with using two-locus statistics in modern populations jointly with one-locus statistics in ancient DNA.

Third, and more importantly, archaic admixture can hide in the blind spot of classical statistics, and widely used demographic models for simulating genomes underestimate LD between low frequency variants in populations around the globe, especially in Africa. This large bias affects neither the distribution of allele frequencies nor the amount of correlation measured by *D*^2^, but it may impact analyses aiming to identify disease variants based on overrepresentation of rare variants in specific genes or pathways. Thus both statistical and population geneticists would benefit from including archaic admixture into baseline models of human genomic diversity.

## Acknowledgements

The authors thank Brenna Henn, Mathias Steinrücken, Ryan Gutenkunst, and Chris Gignoux for useful discussions, and Ryan Gutenkunst for also making his source code open and accessible. We also thank Nick Patterson and an anonymous reviewer for useful comments that improved this manuscript.

