## Appendix for "Models of archaic admixture and recent history from two-locus statistics"

### Supporting material for: Models of archaic admixture and recent history from two-locus statistics

May 25, 2019

#### S1.1 Computing moment equations

In this section we describe a systematic approach for computing transition functions under drift, recombination, mutation, and selection for a system of statistics that include $\mathbb{E}[D^m]$. For any term $\mathbb{E}[f(D,p,q)]$, we change variables to the space of haplotype frequencies. $c_1$ is the frequency or count of type $AB$ haplotype, $c_2$ for $Ab$, $c_3$ for $aB$, and $c_4$ for $ab$, so $D = c_1c_4 - c_2c_3$, $p = c_1 + c_2$, $q = c_1 + c_3$. We then compute transition probabilities on the monomial expansion of this transformation in '$c$'-space, and then change variables back to $(p, q, D)$-space and simplify.

For example, for $\mathbb{E}[D]$, we transform the expectation to

$$\mathbb{E}[D] = \mathbb{E}[c_1c_4 - c_2c_3] = \mathbb{E}[c_1c_4] - \mathbb{E}[c_2c_3].$$

Then for each of the expectations, we calculate its change over one generation due to drift, recombination, mutation, or selection. For example, in the case of drift we find expectations after one generation by considering copying probabilities (described in Section S1.1.1). Then

$$\mathbb{E}[c_1c_4]_{t+1} = \left(1 - \frac{1}{2N(t)}\right)\mathbb{E}[c_1c_4]_t \tag{S1}$$

and

$$\mathbb{E}[c_2c_3]_{t+1} = \left(1 - \frac{1}{2N(t)}\right)\mathbb{E}[c_2c_3]_t. \tag{S2}$$

We then convert back to $(p, q, D)$-space and simplify, obtaining

$$\mathbb{E}[D]_{t+1} = \mathbb{E}[c_1c_4]_{t+1} - \mathbb{E}[c_2c_3]_{t+1} = \left(1 - \frac{1}{2N(t)}\right)\mathbb{E}[D]_t. \tag{S3}$$

Throughout, we assume populations are in Hardy-Weinberg equilibrium and randomly mating. In addition, we assume that the recombination rate $r$, migration rates $m_{ij}$, and mutation rate $u$ are small enough, and population sizes $N_i$ large enough, so that the product of $r$, $m_{ij}$, $u$, and $\frac{1}{N_i}$ may be ignored. In other words, we assume copying, recombination, mutation, and migration rates are small enough so that at most a single event occurs within the $n \ll N$ tracked lineages in a given generation.

##### S1.1.1 Drift

We compute how $\mathbb{E}[c_1^{k_1}c_2^{k_2}c_3^{k_3}c_4^{k_4}]$ is expected to change in a given generation under the action of drift. Let $k = \sum_{i=1}^4 k_i$. We imagine tracking $k$ lineages in the population, each of which is one of the four haplotypes. The expectation is proportional to the probability that $k_1$ of the lineages are in state $c_1$, $k_2$ are of type $c_2$, etc. We assume $k \ll N(t)$ so that the probability of a drift event (where one lineage copies itself over another within the $k$ tracked lineages) is small, and we may assume that at most one such event occurs in any given generation.

The probability of a copying event among $k$ lineages in generation $t$ is

$$P(\text{copying event}) = \frac{1}{2N(t)}\binom{k}{2}. \tag{S4}$$

Given that a drift event occurs, we can compute the probability of any possible transition. For example,

$$\begin{aligned} P((k_1,k_2,k_3,k_4) \to (k_1+1, k_2-1, k_3, k_4)) &= P(\text{choose one of each } c_1 \text{ and } c_2) \cdot P(c_1 \text{ copies over } c_2) \\ &= \frac{\binom{k_1}{1}\binom{k_2}{1}}{\binom{k}{2}} \times \frac{1}{2} \\ &= \frac{k_1k_2}{k(k-1)}. \end{aligned} \tag{S5}$$

We combine all possible copying events and find $\mathbb{E}[c_1^{k_1}c_2^{k_2}c_3^{k_3}c_4^{k_4}]_{t+1}$ under drift

$$\begin{aligned}\mathbb{E}\left[c_1^{k_1}c_2^{k_2}c_3^{k_3}c_4^{k_4}\right]_{t+1} =& (1-P(\text{copying event}))\mathbb{E}\left[c_1^{k_1}c_2^{k_2}c_3^{k_3}c_4^{k_4}\right] \\ &+ P(\text{copying event})\mathbb{E}\left[c_1^{k_1}c_2^{k_2}c_3^{k_3}c_4^{k_4}\cdot\sum_{1\le i,j\le 4}\left(\frac{k_ik_j}{k(k-1)}\frac{c_i}{c_j}\delta_{i\neq j}+\frac{k_i(k_i-1)}{k(k-1)}\delta_{i=j}\right)\right].\end{aligned} \tag{S6}$$

We wrote the $D^2$ drift matrix in the main text. The system of statistics for $D^3$ is

$$\mathbf{y} = \begin{pmatrix}\mathbb{E}[D^3]\\ \mathbb{E}[D^2z]\\ \mathbb{E}[D\pi_2]\\ \mathbb{E}[\pi z]\\ \mathbb{E}[D\sigma_1]\\ \mathbb{E}[z\sigma_1]\\ \mathbb{E}[D]\\ \mathbb{E}[z]\end{pmatrix},$$

where $\sigma_1 = p(1-p)+q(1-q)$, and has transition matrix

$$\mathcal{D} = \frac{1}{2N}\begin{pmatrix}-6 & 3 & 3 & 0 & 0 & 0 & 0 & 0\\ 4 & -11 & 16 & 1 & -4 & 0 & 1 & 0\\ 0 & 1 & -11 & 0 & 1 & 0 & 0 & 0\\ 0 & 0 & 36 & -6 & -6 & 0 & 1 & 0\\ 0 & 0 & 0 & 0 & -6 & 0 & 2 & 0\\ 0 & 0 & 0 & 0 & 12 & -3 & -4 & 0\\ 0 & 0 & 0 & 0 & 0 & 0 & -1 & 0\\ 0 & 0 & 0 & 0 & 0 & 0 & 4 & 0\end{pmatrix}.$$

For the $D^4$ system, with terms

$$\mathbf{y} = \begin{pmatrix}\mathbb{E}[D^4]\\ \mathbb{E}[D^3z]\\ \mathbb{E}[D^2\pi_2]\\ \mathbb{E}[D\pi z]\\ \mathbb{E}[\pi_2^2]\\ \mathbb{E}[D^2\sigma_1]\\ \mathbb{E}[Dz\sigma_1]\\ \mathbb{E}[\pi_2\sigma_1]\\ \mathbb{E}[\sigma_2]\\ \mathbb{E}[D^2]\\ \mathbb{E}[Dz]\\ \mathbb{E}[\pi_2]\\ \mathbb{E}[\sigma_1]\end{pmatrix},$$

the transition matrix is

$$\mathcal{D} = \frac{1}{2N} \begin{pmatrix} -10 & 6 & 6 & 0 & 0 & 0 & 0 & 0 & 0 & 0 & 0 & 0 & 0 \\ 4 & -18 & 48 & 3 & 0 & -12 & 0 & 0 & 0 & 3 & 0 & 0 & 0 \\ 0 & 1 & -21 & 1 & 1 & 2 & 0 & 0 & 0 & 0 & 0 & 0 & 0 \\ 0 & 0 & 36 & -19 & 0 & -6 & 1 & 0 & 0 & 1 & 0 & 0 & 0 \\ 0 & 0 & 0 & 4 & -12 & 0 & 0 & 1 & 0 & 0 & 0 & 0 & 0 \\ 0 & 0 & 0 & 0 & 0 & -12 & 1 & 1 & 0 & 4 & 0 & 0 & 0 \\ 0 & 0 & 0 & 0 & 0 & 12 & -12 & 0 & 0 & -4 & 2 & 0 & 0 \\ 0 & 0 & 0 & 0 & 0 & 0 & 2 & -7 & 0 & 0 & 0 & 2 & 0 \\ 0 & 0 & 0 & 0 & 0 & 0 & 0 & 0 & -6 & 0 & 0 & 0 & 1 \\ 0 & 0 & 0 & 0 & 0 & 0 & 0 & 0 & 0 & -3 & 1 & 1 & 0 \\ 0 & 0 & 0 & 0 & 0 & 0 & 0 & 0 & 0 & 4 & -5 & 0 & 0 \\ 0 & 0 & 0 & 0 & 0 & 0 & 0 & 0 & 0 & 0 & 1 & -2 & 0 \\ 0 & 0 & 0 & 0 & 0 & 0 & 0 & 0 & 0 & 0 & 0 & 0 & -1 \end{pmatrix}.$$

In general, the block upper triangular structure reflects the hierarchy of the moments system (Figure S1), and the sparseness allows for rapid integration.

##### S1.1.2 Recombination

Recombination changes $D$ in one generation at rate directly proportional to the recombination distance between the two loci. This can be seen by considering the effect of recombination on the terms $\mathbb{E}[c_1 c_4]$ and $\mathbb{E}[c_2 c_3]$. Recombination occurs within one haplotype with probability $r$, assuming $r$ is small. If recombination occurs within the $AB$ ($c_1$) haplotype, our lineage changes state to $Ab$ with probability $\frac{1}{2}(1-q)$ or to $aB$ with probability $\frac{1}{2}(1-p)$, and is otherwise unchanged. Similarly, recombination in the $ab$ ($c_4$) haplotype gives rise to $Ab$ with probability $\frac{1}{2}p$ or $aB$ with probability $\frac{1}{2}q$. Summing together these probabilities and simplifying, the probability that recombination removes this haplotype pair's contribution to $D$ is $r$. Thus, under recombination, the expected value of $D$ decays as

$$E[D]_{t+1} = (1-r)E[D]_t.$$

Assuming that $r \ll 1$, the expected decay of LD is, to leading order,

$$E[D]_{t+1} = \left(1 - \frac{1}{2N} - r\right) E[D]_t.$$

In general, haplotype frequencies are expected to change over one generations according to

$$\begin{aligned} c_1' &= c_1 - rD \\ c_2' &= c_2 + rD \\ c_3' &= c_3 + rD \\ c_4' &= c_4 - rD. \end{aligned}$$

Then $\mathbb{E}[\prod c_i^{k_i}]_{t+1} = \mathbb{E}[\prod c_i'^{k_i}]$. Recombination does not change the expected allele frequencies at each locus, so for any moments $\mathbb{E}[D^\alpha f(p,q)]$, this simplifies to the recursion

$$\mathbb{E}[D^\alpha f(p,q)]_{t+1} = (1-\alpha r)\mathbb{E}[D^\alpha f(p,q)]_t. \tag{S7}$$

##### S1.1.3 Mutation

In the infinite-sites model, mutations are assumed to occur once at any given locus (i.e., no recurrent or back mutation). A two-locus pair that is observed to be polymorphic at both loci must have experienced a mutation first at one locus, and then a second to occur at the paired locus. Thus, while $\mathbb{E}[H] = \mathbb{E}[2p(1-p)] \propto \theta$, the joint heterozygosity $\mathbb{E}[p(1-p)q(1-q)] \propto \theta^2$.

In the Hill-Robertson system, a mutation can "create" one-locus diversity from an invariant sites, and create two-locus diversity from one-locus diversity. Other terms in the Hill-Robertson system are unchanged by mutation. For higher order systems, we also have terms of the form $\mathbb{E}[\pi_2\sigma_i] = \mathbb{E}[p(1-p)q(1-q)(p^i(1-p)^i + q^i(1-q)^i)]$ that change due to mutation:

$$\begin{aligned}
\Delta_{\mathcal{U}}\mathbb{E}[\sigma_1] &= \frac{\theta}{2}, \\
\Delta_{\mathcal{U}}\mathbb{E}[\pi_2] &= \frac{\theta}{2}\mathbb{E}[p(1-p) + q(1-q)], \\
\Delta_{\mathcal{U}}\mathbb{E}[\pi_2\sigma_i] &= \frac{\theta}{2}\mathbb{E}[\sigma_i],
\end{aligned}$$

where $\Delta_{\mathcal{U}}$ is used to denote $\mathbb{E}[\cdot]_{t+1} - \mathbb{E}[\cdot]_t$ due to mutation. Here, we have assumed that the mutation rates are equal at the left and right locus, although this approach allows for differing mutation rates between the two loci, as in Ohta and Kimura (1969a).

Reversible mutations are also handled in a similarly straightforward manner. Under a reversible mutation model, we find the same recursions due to mutation in the infinite sites model, with the additional decay of each term $f(p, q, D)$ in $\mathbf{y}$ as

$$\Delta_{\mathcal{U}}\mathbb{E}[f(p, q, D)] = -k\frac{\theta}{2}\mathbb{E}[f(p, q, D)],$$

where $k$ is the sample-size order of the term (that is, the number of sampled haplotypes required to estimate that term, as described in the Section S1.1.1) (Ohta and Kimura, 1969a).

###### S1.1.4 Selection in the Hill-Robertson system

While the Hill-Robertson system closes under drift and recombination, the $D^2$ system (and all other orders) does not close under selection. Here, we consider a simple selection model, with additive selection acting on the $A$ allele at the left locus (with selection strength $s$, $|s| \ll 1$), while the right locus remains neutral. Thus, selection acts for or against $AB$ and $Ab$ haplotypes, with strength $1 + s$ relative to $aB$ and $ab$ haplotypes. This selection model is relevant to computing the expect LD between a selected site and a neutral marker separated by recombination distance $r$.

In this setting, we compute how selection is expected to change $D^2$, $D(1-2p)(1-2q)$, and $p(1-p)q(1-q)$, in expectation. Again we change variables to $c$-space and compute how selection is expected to change terms in the monomial expansion. We denote $c_i'$ to be the expected frequency of type $i$ after one generation. For example,

$$\begin{aligned}
c_1' &= \frac{c_1(1+s)}{(c_1+c_2)(1+s) + c_3 + c_4} \\
&= \frac{c_1(1+s)}{1 + (c_1+c_2)s} \\
&\approx c_1(1+s)(1-(c_1+c_2)s) \approx c_1(1+(1-p)s),
\end{aligned} \tag{S8}$$

to first order in $s$. Similarly,

$$\begin{aligned}
c_2' &\approx c_2(1+(1-p)s), \\
c_3' &\approx c_3(1-ps), \\
c_4' &\approx c_4(1-ps).
\end{aligned}$$

Then in one generation,

$$\begin{aligned}
\mathbb{E}\left[{c_1'}^{k_1}{c_2'}^{k_2}{c_3'}^{k_3}{c_4'}^{k_4}\right] &\approx {c_1}^{k_1}{c_2}^{k_2}{c_3}^{k_3}{c_4}^{k_4}\left(1 + k_1(1-p)s + k_2(1-p)s - k_3ps - k_4ps\right) \\
&\approx {c_1}^{k_1}{c_2}^{k_2}{c_3}^{k_3}{c_4}^{k_4}\left(1 + (k_1+k_2-kp)s\right),
\end{aligned}$$

where $k = \sum_i k_i$.

In one generation, the change in moments $\Delta_{\mathcal{S}}\mathbb{E}[\cdot] = \mathbb{E}[\cdot]_{t+1} - \mathbb{E}[\cdot]_t$ due to selection is

$$\Delta_{\mathcal{S}}\mathbb{E}[D^2] = 2s\mathbb{E}[D^2(1-2p)]$$

$$\Delta_{\mathcal{S}}\mathbb{E}[D(1-2p)(1-2q)] = -2s\mathbb{E}[D^2(1-2p)] + \frac{3}{2}s\mathbb{E}[D(1-2p)^2(1-2q)] - \frac{s}{2}\mathbb{E}[D(1-2p)]$$
$$\Delta_{\mathcal{S}}\mathbb{E}[p(1-p)q(1-q)] = s\mathbb{E}[p(1-p)q(1-q)] + s\mathbb{E}[Dp(1-p)(1-2q)],$$

Thus we need to include additional terms: $\mathbb{E}[D^2(1-2p)]$, $\mathbb{E}[D(1-2p)^2(1-2q)]$, etc. These additional terms require their own additional terms, so that this system grows and includes all terms (in expectation)

$$\begin{pmatrix} D^2p^i(1-p)^i(1-2p)^j \\ Dp^i(1-p)^i(1-2p)^j(1-2q) \\ p^i(1-p)^i(1-2p)^jq(1-q) \end{pmatrix}, \forall i, j \geq 0.$$

To solve this system, we will require a moments closure approximation. This could be achieved, for example, by approximating $D^2p^i(1-p)^i(1-2p)^j$ as a linear combination of terms of lower order, $\{D^2p^{i-k}(1-p)^{i-k}(1-2p)^{j-l}\}$, through a jackknife approximation (Jouganous *et al.*, 2017). Alternatively, we could choose some values $i$ and $j$ to truncate the system and approximate the necessary higher order terms as $\mathbb{E}[f(p,q,D)]$ as $\mathbb{E}[g(p,q,D)]\mathbb{E}[h(p,q,D)]$ where $gh = f$, which leads to a nonlinear system of ODEs. While beyond the scope of this paper, work is ongoing to assess accuracy of closure approximations and incorporate models of selection into multi-population LD models.

#### S1.2 Multiple populations

In this section, we describe the multi-population basis analogous to the Hill-Robertson system for a single populations. We derive recursion equations for the multi-population basis under migration and admixture events.

##### S1.2.1 Population splits

Consider a single populations (denoted 0) that splits into two populations (denoted 1 and 2). In the time of the population split ($t_0$), expected two-locus statistics in populations 1 and 2 are equal to those in population 0. This can be seen by considering the probability of sampling haplotypes in the two split populations. We compute terms of the form $\prod_{j=1}^2\prod_{i=1}^4 {c_{j,i}}^{k_{j,i}}$, where $c_{j,i}$ denotes the probability of sampling haplotype $i$ in population $j$. We observe that at the time of the split, $c_{j,i} = c_{0,i}$, since expected haplotype frequencies in the split populations are equal to expected haplotype frequencies in the parental population. Then,

$$\prod_{j=1}^{2}\prod_{i=1}^{4} {c_{j,i}}^{k_{j,i}} = \prod_{j=1}^{2}\prod_{i=1}^{4} {c_{0,i}}^{k_{j,i}} = \prod_{i=1}^{4} {c_{0,i}}^{k_{1,i}+k_{2,i}}. \tag{S9}$$

Thus
$$\mathbb{E}[D_1^2]_{t_0} = \mathbb{E}[D_2^2]_{t_0} = \mathbb{E}[D_0^2]_{t_0},$$
$$\mathbb{E}[D_1(1-2p_1)(1-2q_1)]_{t_0} = \mathbb{E}[D_2(1-2p_2)(1-2q_2)]_{t_0} = \mathbb{E}[D_0(1-2p_0)(1-2q_0)]_{t_0},$$

and so on. Additionally, we consider $\mathbb{E}[D_1D_2]$, the covariance of $D$ across populations 1 and 2, which initially is

$$\begin{aligned}\mathbb{E}[D_1D_2]_{t_0} &= \mathbb{E}[(c_{1,1}c_{1,4} - c_{1,2}c_{1,3})(c_{2,1}c_{2,4} - c_{2,2}c_{2,3})]_{t_0} \\ &= \mathbb{E}[c_{1,1}c_{1,4}c_{2,1}c_{2,4}]_{t_0} - \mathbb{E}[c_{1,1}c_{1,4}c_{2,2}c_{2,3}]_{t_0} - \mathbb{E}[c_{1,2}c_{1,3}c_{2,1}c_{2,4}]_{t_0} + \mathbb{E}[c_{1,2}c_{1,3}c_{2,2}c_{2,3}]_{t_0} \\ &= \mathbb{E}[c_{0,1}^2c_{0,4}^2]_{t_0} - 2\mathbb{E}[c_{0,1}c_{0,2}c_{0,3}c_{0,4}]_{t_0} + \mathbb{E}[c_{0,2}^2c_{0,3}^2]_{t_0} \\ &= \mathbb{E}[D_0^2]_{t_0}.\end{aligned}$$

In the absence of migration, the $D^2$-systems in each of the populations evolves according to the Hill-Robertson equations, while

$$\mathbb{E}[D_1D_2]_{t+1} = \left(1 - \frac{1}{2N_1} - \frac{1}{2N_2} - 2r\right)\mathbb{E}[D_1D_2]_t. \tag{S10}$$

##### S1.2.2 Migration

With the inclusion of migration, additional moments are needed to obtain a closed system. We write the full basis with migration as

$$\mathbf{y} = \begin{pmatrix} \mathbb{E}[D_i D_j] \\ \mathbb{E}[D_i z_{j,k}] \\ \mathbb{E}[\pi_2(i,j;k,l)] \\ \mathbb{E}[H_{i,j}] \end{pmatrix},$$

where $i, j, k, l$ index populations, and

$$\begin{aligned} D_i z_{j,k} &= D_i(1-2p_j)(1-2q_k), \\ \pi_2(i,j;k,l) &= \frac{1}{4}p_i(1-p_j)q_k(1-q_l) + \frac{1}{4}p_i(1-p_j)q_l(1-q_k) \\ &\quad + \frac{1}{4}p_j(1-p_i)q_k(1-q_l) + \frac{1}{4}p_j(1-p_i)q_l(1-q_k), \\ H_{i,j} &= \frac{1}{2}p_i(1-p_j) + \frac{1}{2}p_j(1-p_i), \end{aligned}$$

Initial values for each cross term are found using Equation S9 in Section S1.2.1.

The number of terms in the system grows in the number $P$ of populations, as $P^3 + \left(\frac{P(P+1)}{2}\right)^2 + 3\frac{P(P+1)}{2}$. While the size of the full joint AFS grows exponentially in $P$, the multi-population Hill-Robertson system remains manageable for even large $P$: for example, the 10-population system has 4,190 terms, which is not at all computationally burdensome for a sparse, linear system. When the left and right loci have equal mutation rate, as we assume in most of our models, redundant terms exist in this basis so the system may be further reduced in size.

We want to know how migration changes expected statistics in our system. We again work in $c$-space, where $c_{i1}$ represents the haplotype $AB$ in population $i$, $c_{i2}$ represents $Ab$ in population $i$, $c_{i3}$ represents $aB$ in $i$, and $c_{i4}$ represents $ab$ in $i$. We consider terms of the general form

$$\mathbb{E}\left[\prod_{i=1}^{P}\prod_{l=1}^{4} c_{il}^{k_{il}}\right],$$

and set $k = \sum_{i,l} k_{il}$ as the total number of tracked lineages in our subsample across all populations, corresponding to the sample size (order) of the moment.

We denote migration rates $m_{ij}$ as the probability that a lineage in population $j$ is replaced by a migrant lineage from population $i$ (in other words, it is the probability that a lineage in $i$ has a parent in $j$). We assume $m_{ij} \ll 1$ so that at most a single migrant replaces a lineage among the sample of size $k \ll N_j$ in population $j$ in any generation. Then in one generation, the change due to migration is

$$\Delta_{\mathcal{M}}\mathbb{E}\left[\prod_{i=1}^{P}\prod_{l=1}^{4} c_{il}^{k_{il}}\right]_t = \sum_{i=1}^{P}\sum_{j=1}^{P}\delta_{i\neq j} m_{ij}\sum_{l=1}^{4}\mathbb{E}\left[\frac{k_{il}(c_{il}-c_{jl})}{c_{jl}}\right]. \tag{S11}$$

In words, we sum over each pair of populations $i, j$ and consider the probability that lineages of each type migrate from population $i$ and copies over a lineage within our sample in population $j$.

As a simple example, consider the expected frequency at a single locus $p_1$ in population 1 (among $P$ total populations). Using the above formula, after one generation we find

$$\Delta_{\mathcal{M}}\mathbb{E}[p_1] = \sum_{i=2}^{P} m_{i1}(\mathbb{E}[p_i] - \mathbb{E}[p_1]).$$

Expected changes due to migration may be found for each term in the system.

For two populations, with migration rates $m_{12}$ from population 1 to 2, and $m_{21}$ from population 2 to 1, considering the terms (in expectation) in the system

$$\mathbf{y} = \begin{pmatrix} D_1^2 \\ D_1 D_2 \\ D_2^2 \\ D_1 z_{1,1} \\ D_1 z_{1,2} \\ D_1 z_{2,1} \\ D_1 z_{2,2} \\ D_2 z_{1,1} \\ D_2 z_{1,2} \\ D_2 z_{2,1} \\ D_2 z_{2,2} \\ \pi_2(1,1;1,1) \\ \pi_2(1,1;1,2) \\ \pi_2(1,1;2,2) \\ \pi_2(1,2;1,1) \\ \pi_2(1,2;1,2) \\ \pi_2(1,2;2,2) \\ \pi_2(2,2;1,1) \\ \pi_2(2,2;1,2) \\ \pi_2(2,2;2,2) \\ H_{1,1} \\ H_{1,2} \\ H_{2,2} \end{pmatrix},$$

we obtain the migration operator

$$\mathcal{M} = m_{12} \begin{pmatrix}
0 & 0 & 0 & 0 & 0 & 0 & 0 & 0 & 0 & 0 & 0 & 0 & 0 & 0 & 0 & 0 & 0 & 0 & 0 & 0 & 0 & 0 & 0 \\
1 & -1 & 0 & 0 & 0 & 0 & 0 & 0 & 0 & 0 & 0 & 0 & 0 & 0 & 0 & 0 & 0 & 0 & 0 & 0 & 0 & 0 & 0 \\
0 & 2 & -2 & 0 & 0 & 0 & 0 & 0 & 0 & 0 & 0 & 0 & 0 & 0 & 0 & 0 & 0 & 0 & 0 & 0 & 0 & 0 & 0 \\
0 & 0 & 0 & 0 & 0 & 0 & 0 & 0 & 0 & 0 & 0 & 0 & 0 & 0 & 0 & 0 & 0 & 0 & 0 & 0 & 0 & 0 & 0 \\
0 & 0 & 0 & 1 & -1 & 0 & 0 & 0 & 0 & 0 & 0 & 0 & 0 & 0 & 0 & 0 & 0 & 0 & 0 & 0 & 0 & 0 & 0 \\
0 & 0 & 0 & 1 & 0 & -1 & 0 & 0 & 0 & 0 & 0 & 0 & 0 & 0 & 0 & 0 & 0 & 0 & 0 & 0 & 0 & 0 & 0 \\
0 & 0 & 0 & 0 & 1 & 1 & -2 & 0 & 0 & 0 & 0 & 0 & 0 & 0 & 0 & 0 & 0 & 0 & 0 & 0 & 0 & 0 & 0 \\
0 & 0 & 0 & 1 & 0 & 0 & 0 & -1 & 0 & 0 & 0 & 4 & -4 & 0 & -4 & 4 & 0 & 0 & 0 & 0 & 0 & 0 & 0 \\
0 & 0 & 0 & 0 & 1 & 0 & 0 & 1 & -2 & 0 & 0 & 0 & 4 & -4 & 0 & -4 & 4 & 0 & 0 & 0 & 0 & 0 & 0 \\
0 & 0 & 0 & 0 & 0 & 1 & 0 & 1 & 0 & -2 & 0 & 0 & 0 & 0 & 4 & -4 & 0 & -4 & 4 & 0 & 0 & 0 & 0 \\
0 & 0 & 0 & 0 & 0 & 0 & 1 & 0 & 1 & 1 & -3 & 0 & 0 & 0 & 0 & 4 & -4 & 0 & -4 & 4 & 0 & 0 & 0 \\
0 & 0 & 0 & 0 & 0 & 0 & 0 & 0 & 0 & 0 & 0 & 0 & 0 & 0 & 0 & 0 & 0 & 0 & 0 & 0 & 0 & 0 & 0 \\
0 & 0 & 0 & 0 & 0 & 0 & 0 & 0 & 0 & 0 & 0 & 1 & -1 & 0 & 0 & 0 & 0 & 0 & 0 & 0 & 0 & 0 & 0 \\
0 & 0 & 0 & 0 & 0 & 0 & 0 & 0 & 0 & 0 & 0 & 0 & 2 & -2 & 0 & 0 & 0 & 0 & 0 & 0 & 0 & 0 & 0 \\
0 & 0 & 0 & 0 & 0 & 0 & 0 & 0 & 0 & 0 & 0 & 1 & 0 & 0 & -1 & 0 & 0 & 0 & 0 & 0 & 0 & 0 & 0 \\
0 & 0 & 0 & 0 & 0 & 0 & 0 & 0 & 0 & 0 & 0 & 0 & 1 & 0 & 1 & -2 & 0 & 0 & 0 & 0 & 0 & 0 & 0 \\
0 & 0 & 0 & 0 & 0 & 0 & 0 & 0 & 0 & 0 & 0 & 0 & 0 & 1 & 0 & 2 & -3 & 0 & 0 & 0 & 0 & 0 & 0 \\
0 & 0 & 0 & 0 & 0 & 0 & 0 & 0 & 0 & 0 & 0 & 0 & 0 & 0 & 2 & 0 & 0 & -2 & 0 & 0 & 0 & 0 & 0 \\
0 & 0 & 0 & 0 & 0 & 0 & 0 & 0 & 0 & 0 & 0 & 0 & 0 & 0 & 0 & 2 & 0 & 1 & -3 & 0 & 0 & 0 & 0 \\
0 & 0 & 0 & 0 & 0 & 0 & 0 & 0 & 0 & 0 & 0 & 0 & 0 & 0 & 0 & 0 & 2 & 0 & 2 & -4 & 0 & 0 & 0 \\
0 & 0 & 0 & 0 & 0 & 0 & 0 & 0 & 0 & 0 & 0 & 0 & 0 & 0 & 0 & 0 & 0 & 0 & 0 & 0 & 0 & 0 & 0 \\
0 & 0 & 0 & 0 & 0 & 0 & 0 & 0 & 0 & 0 & 0 & 0 & 0 & 0 & 0 & 0 & 0 & 0 & 0 & 0 & 1 & -1 & 0 \\
0 & 0 & 0 & 0 & 0 & 0 & 0 & 0 & 0 & 0 & 0 & 0 & 0 & 0 & 0 & 0 & 0 & 0 & 0 & 0 & 0 & 2 & -2
\end{pmatrix}$$

$$+ m_{21} \begin{pmatrix}
-2 & 2 & 0 & 0 & 0 & 0 & 0 & 0 & 0 & 0 & 0 & 0 & 0 & 0 & 0 & 0 & 0 & 0 & 0 & 0 & 0 & 0 & 0 \\
0 & -1 & 1 & 0 & 0 & 0 & 0 & 0 & 0 & 0 & 0 & 0 & 0 & 0 & 0 & 0 & 0 & 0 & 0 & 0 & 0 & 0 & 0 \\
0 & 0 & 0 & 0 & 0 & 0 & 0 & 0 & 0 & 0 & 0 & 0 & 0 & 0 & 0 & 0 & 0 & 0 & 0 & 0 & 0 & 0 & 0 \\
0 & 0 & 0 & -3 & 1 & 1 & 0 & 1 & 0 & 0 & 0 & 4 & -4 & 0 & -4 & 4 & 0 & 0 & 0 & 0 & 0 & 0 & 0 \\
0 & 0 & 0 & 0 & -2 & 0 & 1 & 0 & 1 & 0 & 0 & 0 & 4 & -4 & 0 & -4 & 4 & 0 & 0 & 0 & 0 & 0 & 0 \\
0 & 0 & 0 & 0 & 0 & -2 & 1 & 0 & 0 & 1 & 0 & 0 & 0 & 0 & 4 & -4 & 0 & -4 & 4 & 0 & 0 & 0 & 0 \\
0 & 0 & 0 & 0 & 0 & 0 & -1 & 0 & 0 & 0 & 1 & 0 & 0 & 0 & 0 & 4 & -4 & 0 & -4 & 4 & 0 & 0 & 0 \\
0 & 0 & 0 & 0 & 0 & 0 & 0 & -2 & 1 & 1 & 0 & 0 & 0 & 0 & 0 & 0 & 0 & 0 & 0 & 0 & 0 & 0 & 0 \\
0 & 0 & 0 & 0 & 0 & 0 & 0 & 0 & -1 & 0 & 1 & 0 & 0 & 0 & 0 & 0 & 0 & 0 & 0 & 0 & 0 & 0 & 0 \\
0 & 0 & 0 & 0 & 0 & 0 & 0 & 0 & 0 & -1 & 1 & 0 & 0 & 0 & 0 & 0 & 0 & 0 & 0 & 0 & 0 & 0 & 0 \\
0 & 0 & 0 & 0 & 0 & 0 & 0 & 0 & 0 & 0 & 0 & 0 & 0 & 0 & 0 & 0 & 0 & 0 & 0 & 0 & 0 & 0 & 0 \\
0 & 0 & 0 & 0 & 0 & 0 & 0 & 0 & 0 & 0 & 0 & -4 & 2 & 0 & 2 & 0 & 0 & 0 & 0 & 0 & 0 & 0 & 0 \\
0 & 0 & 0 & 0 & 0 & 0 & 0 & 0 & 0 & 0 & 0 & 0 & -3 & 1 & 0 & 2 & 0 & 0 & 0 & 0 & 0 & 0 & 0 \\
0 & 0 & 0 & 0 & 0 & 0 & 0 & 0 & 0 & 0 & 0 & 0 & 0 & -2 & 0 & 0 & 2 & 0 & 0 & 0 & 0 & 0 & 0 \\
0 & 0 & 0 & 0 & 0 & 0 & 0 & 0 & 0 & 0 & 0 & 0 & 0 & 0 & -3 & 2 & 0 & 1 & 0 & 0 & 0 & 0 & 0 \\
0 & 0 & 0 & 0 & 0 & 0 & 0 & 0 & 0 & 0 & 0 & 0 & 0 & 0 & 0 & -2 & 1 & 0 & 1 & 0 & 0 & 0 & 0 \\
0 & 0 & 0 & 0 & 0 & 0 & 0 & 0 & 0 & 0 & 0 & 0 & 0 & 0 & 0 & 0 & -1 & 0 & 0 & 1 & 0 & 0 & 0 \\
0 & 0 & 0 & 0 & 0 & 0 & 0 & 0 & 0 & 0 & 0 & 0 & 0 & 0 & 0 & 0 & 0 & -2 & 2 & 0 & 0 & 0 & 0 \\
0 & 0 & 0 & 0 & 0 & 0 & 0 & 0 & 0 & 0 & 0 & 0 & 0 & 0 & 0 & 0 & 0 & 0 & -1 & 1 & 0 & 0 & 0 \\
0 & 0 & 0 & 0 & 0 & 0 & 0 & 0 & 0 & 0 & 0 & 0 & 0 & 0 & 0 & 0 & 0 & 0 & 0 & 0 & 0 & 0 & 0 \\
0 & 0 & 0 & 0 & 0 & 0 & 0 & 0 & 0 & 0 & 0 & 0 & 0 & 0 & 0 & 0 & 0 & 0 & 0 & 0 & -2 & 2 & 0 \\
0 & 0 & 0 & 0 & 0 & 0 & 0 & 0 & 0 & 0 & 0 & 0 & 0 & 0 & 0 & 0 & 0 & 0 & 0 & 0 & 0 & -1 & 1 \\
0 & 0 & 0 & 0 & 0 & 0 & 0 & 0 & 0 & 0 & 0 & 0 & 0 & 0 & 0 & 0 & 0 & 0 & 0 & 0 & 0 & 0 & 0
\end{pmatrix}.$$

The two-population drift operator in the migration basis is

$$
\mathcal{D} = \frac{1}{2N_1}\begin{pmatrix}
-3 & 0 & 0 & 1 & 0 & 0 & 0 & 0 & 0 & 0 & 0 & 1 & 0 & 0 & 0 & 0 & 0 & 0 & 0 & 0 & 0 & 0 & 0 \\
0 & -1 & 0 & 0 & 0 & 0 & 0 & 0 & 0 & 0 & 0 & 0 & 0 & 0 & 0 & 0 & 0 & 0 & 0 & 0 & 0 & 0 & 0 \\
0 & 0 & 0 & 0 & 0 & 0 & 0 & 0 & 0 & 0 & 0 & 0 & 0 & 0 & 0 & 0 & 0 & 0 & 0 & 0 & 0 & 0 & 0 \\
4 & 0 & 0 & -5 & 0 & 0 & 0 & 0 & 0 & 0 & 0 & 0 & 0 & 0 & 0 & 0 & 0 & 0 & 0 & 0 & 0 & 0 & 0 \\
0 & 0 & 0 & 0 & -3 & 0 & 0 & 0 & 0 & 0 & 0 & 0 & 0 & 0 & 0 & 0 & 0 & 0 & 0 & 0 & 0 & 0 & 0 \\
0 & 0 & 0 & 0 & 0 & -3 & 0 & 0 & 0 & 0 & 0 & 0 & 0 & 0 & 0 & 0 & 0 & 0 & 0 & 0 & 0 & 0 & 0 \\
0 & 0 & 0 & 0 & 0 & 0 & -1 & 0 & 0 & 0 & 0 & 0 & 0 & 0 & 0 & 0 & 0 & 0 & 0 & 0 & 0 & 0 & 0 \\
0 & 4 & 0 & 0 & 0 & 0 & 0 & 0 & 0 & 0 & 0 & 0 & 0 & 0 & 0 & 0 & 0 & 0 & 0 & 0 & 0 & 0 & 0 \\
0 & 0 & 0 & 0 & 0 & 0 & 0 & 0 & 0 & 0 & 0 & 0 & 0 & 0 & 0 & 0 & 0 & 0 & 0 & 0 & 0 & 0 & 0 \\
0 & 0 & 0 & 0 & 0 & 0 & 0 & 0 & 0 & 0 & 0 & 0 & 0 & 0 & 0 & 0 & 0 & 0 & 0 & 0 & 0 & 0 & 0 \\
0 & 0 & 0 & 0 & 0 & 0 & 0 & 0 & 0 & 0 & 0 & 0 & 0 & 0 & 0 & 0 & 0 & 0 & 0 & 0 & 0 & 0 & 0 \\
0 & 0 & 0 & 1 & 0 & 0 & 0 & 0 & 0 & 0 & 0 & -2 & 0 & 0 & 0 & 0 & 0 & 0 & 0 & 0 & 0 & 0 & 0 \\
0 & 0 & 0 & 0 & 0 & 0 & 0 & 0 & 0 & 0 & 0 & 0 & -1 & 0 & 0 & 0 & 0 & 0 & 0 & 0 & 0 & 0 & 0 \\
0 & 0 & 0 & 0 & 0 & 0 & 0 & 0 & 0 & 0 & 0 & 0 & 0 & -1 & 0 & 0 & 0 & 0 & 0 & 0 & 0 & 0 & 0 \\
0 & 0 & 0 & 0 & 0 & 0 & 0 & 0 & 0 & 0 & 0 & 0 & 0 & 0 & -1 & 0 & 0 & 0 & 0 & 0 & 0 & 0 & 0 \\
0 & 0 & 0 & 0 & 0 & 0 & 0 & 0 & 0 & 0 & 0 & 0 & 0 & 0 & 0 & 0 & 0 & 0 & 0 & 0 & 0 & 0 & 0 \\
0 & 0 & 0 & 0 & 0 & 0 & 0 & 0 & 0 & 0 & 0 & 0 & 0 & 0 & 0 & 0 & 0 & 0 & 0 & 0 & 0 & 0 & 0 \\
0 & 0 & 0 & 0 & 0 & 0 & 0 & 0 & 0 & 0 & 0 & 0 & 0 & 0 & 0 & 0 & 0 & -1 & 0 & 0 & 0 & 0 & 0 \\
0 & 0 & 0 & 0 & 0 & 0 & 0 & 0 & 0 & 0 & 0 & 0 & 0 & 0 & 0 & 0 & 0 & 0 & 0 & 0 & 0 & 0 & 0 \\
0 & 0 & 0 & 0 & 0 & 0 & 0 & 0 & 0 & 0 & 0 & 0 & 0 & 0 & 0 & 0 & 0 & 0 & 0 & 0 & 0 & 0 & 0 \\
0 & 0 & 0 & 0 & 0 & 0 & 0 & 0 & 0 & 0 & 0 & 0 & 0 & 0 & 0 & 0 & 0 & 0 & 0 & 0 & -1 & 0 & 0 \\
0 & 0 & 0 & 0 & 0 & 0 & 0 & 0 & 0 & 0 & 0 & 0 & 0 & 0 & 0 & 0 & 0 & 0 & 0 & 0 & 0 & 0 & 0 \\
0 & 0 & 0 & 0 & 0 & 0 & 0 & 0 & 0 & 0 & 0 & 0 & 0 & 0 & 0 & 0 & 0 & 0 & 0 & 0 & 0 & 0 & 0
\end{pmatrix}
+ \frac{1}{2N_2}\begin{pmatrix}
0 & 0 & 0 & 0 & 0 & 0 & 0 & 0 & 0 & 0 & 0 & 0 & 0 & 0 & 0 & 0 & 0 & 0 & 0 & 0 & 0 & 0 & 0 \\
0 & -1 & 0 & 0 & 0 & 0 & 0 & 0 & 0 & 0 & 0 & 0 & 0 & 0 & 0 & 0 & 0 & 0 & 0 & 0 & 0 & 0 & 0 \\
0 & 0 & -3 & 0 & 0 & 0 & 0 & 0 & 0 & 0 & 1 & 0 & 0 & 0 & 0 & 0 & 0 & 0 & 0 & 1 & 0 & 0 & 0 \\
0 & 0 & 0 & 0 & 0 & 0 & 0 & 0 & 0 & 0 & 0 & 0 & 0 & 0 & 0 & 0 & 0 & 0 & 0 & 0 & 0 & 0 & 0 \\
0 & 0 & 0 & 0 & 0 & 0 & 0 & 0 & 0 & 0 & 0 & 0 & 0 & 0 & 0 & 0 & 0 & 0 & 0 & 0 & 0 & 0 & 0 \\
0 & 0 & 0 & 0 & 0 & 0 & 0 & 0 & 0 & 0 & 0 & 0 & 0 & 0 & 0 & 0 & 0 & 0 & 0 & 0 & 0 & 0 & 0 \\
0 & 4 & 0 & 0 & 0 & 0 & 0 & 0 & 0 & 0 & 0 & 0 & 0 & 0 & 0 & 0 & 0 & 0 & 0 & 0 & 0 & 0 & 0 \\
0 & 0 & 0 & 0 & 0 & 0 & 0 & -1 & 0 & 0 & 0 & 0 & 0 & 0 & 0 & 0 & 0 & 0 & 0 & 0 & 0 & 0 & 0 \\
0 & 0 & 0 & 0 & 0 & 0 & 0 & 0 & -3 & 0 & 0 & 0 & 0 & 0 & 0 & 0 & 0 & 0 & 0 & 0 & 0 & 0 & 0 \\
0 & 0 & 0 & 0 & 0 & 0 & 0 & 0 & 0 & -3 & 0 & 0 & 0 & 0 & 0 & 0 & 0 & 0 & 0 & 0 & 0 & 0 & 0 \\
0 & 0 & 4 & 0 & 0 & 0 & 0 & 0 & 0 & 0 & -5 & 0 & 0 & 0 & 0 & 0 & 0 & 0 & 0 & 0 & 0 & 0 & 0 \\
0 & 0 & 0 & 0 & 0 & 0 & 0 & 0 & 0 & 0 & 0 & 0 & 0 & 0 & 0 & 0 & 0 & 0 & 0 & 0 & 0 & 0 & 0 \\
0 & 0 & 0 & 0 & 0 & 0 & 0 & 0 & 0 & 0 & 0 & 0 & 0 & 0 & 0 & 0 & 0 & 0 & 0 & 0 & 0 & 0 & 0 \\
0 & 0 & 0 & 0 & 0 & 0 & 0 & 0 & 0 & 0 & 0 & 0 & 0 & -1 & 0 & 0 & 0 & 0 & 0 & 0 & 0 & 0 & 0 \\
0 & 0 & 0 & 0 & 0 & 0 & 0 & 0 & 0 & 0 & 0 & 0 & 0 & 0 & 0 & 0 & 0 & 0 & 0 & 0 & 0 & 0 & 0 \\
0 & 0 & 0 & 0 & 0 & 0 & 0 & 0 & 0 & 0 & 0 & 0 & 0 & 0 & 0 & 0 & 0 & 0 & 0 & 0 & 0 & 0 & 0 \\
0 & 0 & 0 & 0 & 0 & 0 & 0 & 0 & 0 & 0 & 0 & 0 & 0 & 0 & 0 & 0 & -1 & 0 & 0 & 0 & 0 & 0 & 0 \\
0 & 0 & 0 & 0 & 0 & 0 & 0 & 0 & 0 & 0 & 0 & 0 & 0 & 0 & 0 & 0 & 0 & -1 & 0 & 0 & 0 & 0 & 0 \\
0 & 0 & 0 & 0 & 0 & 0 & 0 & 0 & 0 & 0 & 0 & 0 & 0 & 0 & 0 & 0 & 0 & 0 & -1 & 0 & 0 & 0 & 0 \\
0 & 0 & 0 & 0 & 0 & 0 & 0 & 0 & 0 & 0 & 1 & 0 & 0 & 0 & 0 & 0 & 0 & 0 & 0 & -2 & 0 & 0 & 0 \\
0 & 0 & 0 & 0 & 0 & 0 & 0 & 0 & 0 & 0 & 0 & 0 & 0 & 0 & 0 & 0 & 0 & 0 & 0 & 0 & 0 & 0 & 0 \\
0 & 0 & 0 & 0 & 0 & 0 & 0 & 0 & 0 & 0 & 0 & 0 & 0 & 0 & 0 & 0 & 0 & 0 & 0 & 0 & 0 & 0 & 0 \\
0 & 0 & 0 & 0 & 0 & 0 & 0 & 0 & 0 & 0 & 0 & 0 & 0 & 0 & 0 & 0 & 0 & 0 & 0 & 0 & 0 & 0 & -1
\end{pmatrix}.
$$

For any number of populations, we have the closed recursion

$$
\mathbf{y}_{t+1} - \mathbf{y}_t = \left(\mathcal{D}_{\mathbf{N}} + \mathcal{M}_{\mathbf{m}} + \mathcal{R}_r + \mathcal{U}_u\right)\mathbf{y}_t, \tag{S12}
$$

where $\mathcal{D}, \mathcal{R}, \mathcal{U}$, and $\mathcal{M}$ are sparse matrices for drift, recombination, mutation, and migration, with population sizes $\mathbf{N}$ and migration rates $\mathbf{m}$.

##### S1.2.3 Admixture

We find the expected value for terms in basis (5) that include a new admixed population. We assume the admixed population arises from two parental populations, with lineages arriving from the two populations with probability $f$ and $1-f$. Computing terms involving the admixed population simply requires enumerating over the haplotype probabilities of lineages arising from each parental population, which have been computed up to the time of admixture, and then simplifying.

Suppose an admixture event between populations 1 (with proportion $f$) and 2 (with proportion $1-f$), with $P$ total populations, forms a new population indexed $P+1$. Expected haplotype frequencies in the admixed population depend on $f$ and the expected haplotype frequencies in the source populations. For example, $c_{P+1,1}$ ($c_1$ in the admixed population) is equal to $fc_{1,1} + (1-f)c_{2,1}$, $c_{P+1,2} = fc_{1,2} + (1-f)c_{2,2}$, and so on. Cavalli-Sforza and Bodmer (1971) (page 69) used this approach to compute $\mathbb{E}[D]$ in an admixed population (Equation 7).

For any statistic $f(\mathbf{p}, \mathbf{q}, \mathbf{D})$, we first convert to $c$-space variables. Then for each term in the $c$-space expansion $\prod_{i=1}^{P+1}\prod_{l=1}^{4} {c_{i,l}}^{k_{i,l}}$, where $c_{i,l}$ represents haplotype $l$ in population $i$, we compute

$$\prod_{i=1}^{P+1}\prod_{l=1}^{4} {c_{i,l}}^{k_{i,l}} = \left(\prod_{i=1}^{P}\prod_{l=1}^{4} {c_{i,l}}^{k_{i,l}}\right)(fc_{1,1}+(1-f)c_{2,1})^{k_{P+1,1}}(fc_{1,2}+(1-f)c_{2,2})^{k_{P+1,2}}$$
$$\times (fc_{1,3}+(1-f)c_{2,3})^{k_{P+1,3}}(fc_{1,4}+(1-f)c_{2,4})^{k_{P+1,4}}.$$

We then convert back to the $(\mathbf{p}, \mathbf{q}, \mathbf{D})$ variables and simplify. If a term does not include any term with $p_{P+1}$, $q_{P+1}$ or $D_{P+1}$, it remains remains unchanged. Otherwise, the term can be written as a linear combination of terms in the $P$-population basis.

For example, with two populations, with contribution $f$ from population 1 and $1-f$ from population 2,

$$\begin{aligned}\mathbb{E}[D_{\text{adm}}^2] =& f^2\mathbb{E}[D_1^2] + (1-f)^2\mathbb{E}[D_2^2] + 2f(1-f)\mathbb{E}[D_1D_2] \\ &+ 2f^2(1-f)\mathbb{E}[D_1\delta] + 2f(1-f)^2\mathbb{E}[D_2\delta] + f^2(1-f)^2\mathbb{E}[\delta^2].\end{aligned} \tag{S13}$$

$\mathbb{E}[D_i\delta]$ and $\mathbb{E}[\delta^2]$ can be written as linear combinations of terms in the multi-population basis (5). Similar equations exist for each new term in the basis with the additional admixed population, and this system can then be integrated forward in time using Equation 6. The full set of equations for arbitrary number of population is implemented in our software `moments.LD`, available at `http://www.bitbucket.org/simongravel/moments`.

#### S1.3 Haplotype frequency spectrum

The allele frequency spectrum (AFS) is the distribution of allele counts in a sample of size $n$, denoted $\Phi_n$. Because $\Phi_n$ is sensitive to demographic and evolutionary processes, it is widely used to infer demographic history, patterns of selection, and mutation rates. Forward-in-time approaches solve the underlying diffusion equation, which describes the time-evolution of the distribution of allele frequencies in one or more populations. In one population, this takes the form

$$\frac{\partial\phi}{\partial t} = \frac{1}{2N}\frac{\partial^2}{\partial x^2}[x(1-x)\phi] - s\frac{\partial}{\partial x}[(h+(1-2h)x)x(1-x)\phi], \tag{S14}$$

where $N$ is the effective population size and can change over time. $\Phi_n$ can then be found by integrating $\phi$ against the binomial sampling distribution.

Analytic solutions to Equation S14 have only been found for simple scenarios, such as steady-state solutions. To compute $\Phi_n$ for non-equilibrium demography, we turn to numerical solutions, e.g. (Evans *et al.*, 2007; Gutenkunst *et al.*, 2009; Lukic and Hey, 2012). Recently, Jouganous *et al.* (2017) recognized that the entries of $\Phi_n$ themselves comprise a moments system that allows for direct integration of $\Phi_n$ without having to numerically solve Equation S14. This system closes under drift, while selection requires a moment-closure approximation.

The two-locus frequency spectrum ($\Psi_n$) is defined similarly to the single locus AFS, but instead tracks the haplotype frequencies of two-locus pairs. We consider a model that permits two alleles at each of the two loci: $A/a$ at the left locus, and $B/b$ at the right. Four haplotypes are possible in the two-locus model $(AB, Ab, aB, ab)$ whose frequencies in the population sum to one. Then $\Psi_n(i,j,k)$ is the expected number of two-locus pairs in a sample of size $n$ in which we observe $i$ copies of type $AB$, $j$ of type $Ab$, $k$ of type $aB$, and $n-i-j-k$ of type $ab$.

The two-locus frequency spectrum can be found at steady-state using a recursion due to Golding (1984). For non-equilibrium demography, Kamm *et al.* (2016) presented a coalescent approach for $\Psi_n$ via a Moran model closely related to the look-down approach of Donnelly and Kurtz (1999). Alternatively, $\Psi_n$ can be found by first solving the associated two-locus diffusion equation $\psi$ (Kimura, 1955; Hill and Robertson, 1966), the density of two-locus haplotype frequencies in the population,

$$\frac{\partial\psi}{\partial t} = \frac{1}{2}\sum\sum_{1\leq i,j\leq 3}\frac{\partial^2}{\partial x_i\partial x_j}\left[\frac{x_i(\delta_{ij}-x_j)\psi}{N(t)}\right] + \frac{\rho}{2}\left(\frac{\partial}{\partial x_1}[D\psi] - \frac{\partial}{\partial x_2}[D\psi] - \frac{\partial}{\partial x_3}[D\psi]\right), \tag{S15}$$

shown here without terms for selection and $D = x_1(1 - x_1 - x_2 - x_3) - x_2x_3$, and then integrating $\psi$ against the multinomial sampling distribution. Ragsdale and Gutenkunst (2017) solved Equation S15 using finite differences, which they used in single-population demographic inference. The advantage to the diffusion approach is that selection is easily incorporated at one or both loci, which allows us to directly model the effect of linked selection with any recombination recombination rate and non-equilibrium demography.

In Sections S1.3.1, S1.3.2, S1.3.3, and S1.3.4 instead of finding a numerical solution to Equation S15, we show that we can directly solve for $\Psi_n$ through a recursion on its entries just as Jouganous *et al.* (2017) proposed for the single-locus AFS. By considering how haplotype frequencies within $n$ tracked lineages are expected to change due to drift, recombination, selection, and mutation, we obtain the recursion

$$\Psi_n^{t+1}(i,j,k) - \Psi_n^t(i,j,k) = \mathcal{D}_{2N(t),n;i,j,k}\Psi_n^t + \mathcal{U}_{\mathbf{u}}\Psi_n^t + \mathcal{R}_r\Psi_{n+1}^t + \mathcal{S}_{\mathbf{s},h}\Psi_{n+2}^t. \tag{S16}$$

Here, $\mathcal{D}$ is a sparse matrix to account for drift, $\mathcal{R}$ accounts for recombination with rate $r$, $\mathcal{S}$ for selection with arbitrary selection and dominance coefficients for each haplotype, and $\mathcal{U}$ is a mutation operator for either an infinite sites or reversible mutation model.

Under drift and mutation, in the neutral case and with no recombination, Equation S16 is closed and can be solved exactly. With selection and recombination, $\Psi_n$ relies on the slightly larger frequency spectra $\Psi_{n+1}$ and $\Psi_{n+2}$, and so the system does not close. Intuitively, drift closes because we are just concerned with lineages copying over each other within the subsample of $n$ tracked lineages. However, we require additional lineages in the case of non-zero recombination and selection. If a recombination event occurs within our $n$ tracked lineages, we require an additional lineage to be drawn from the full population for the recombining lineage to be paired with. In the case of selection, a replacement lineage must be drawn from the entire population, so we need to know the expected distribution of a larger size sample. We close the system using a jackknife extrapolation, which estimates $\Psi_{n+l}$ using the known $\Psi_n$, as was done for the single locus frequency spectrum in Jouganous *et al.* (2017). In practice, we find the jackknife to be reasonably accurate for moderate sample sizes ($n \gtrsim 20$), with accuracy increasing in $n$ (Table S1).

Below, we derive each operator in turn. We consider tracking a subsample of $n$ lineages in the population and find how drift, mutation, recombination, and selection are each expected to change probabilities of two-locus haplotype frequencies in a given generation.

##### S1.3.1 Drift

Just as in Section S1.1.1, transition probabilities for $\Psi_n$ are derived by considering haplotype frequencies within a sample of size $n$. Allele frequencies within the subsample of $n$ lineages change due to drift in one generation if one lineage within the $n$ subsampled lineages copies itself onto another lineage within our subsample. For simplicity, we assume $n \ll N$, so that at most a single copying event occurs between two of the $n$ lineages in any given generation. Generalization to multiple coalescences per generation could be performed as in Jouganous *et al.* (2017).

The probability of a single copying event within the sample in a single generation at time $t$ (to leading order in $1/N$) is

$$P_{n,N}(1 \to 2) = \frac{1}{2N(t)}\binom{n}{2}, \tag{S17}$$

the classical large-population limit from coalescence theory. In the case of a copying event, one lineage is drawn twice and another is not drawn. Haplotype frequencies will only change if the two drawn lineages differ in state.

We compute the probability of all possible haplotype frequency changes ($(i,j,k) \to (i',j',k')$) in one generation, given a coalescent event occurs. For example, frequencies change from $(i,j,k)$ to $(i+1,j-1,k)$ if an $AB$ lineage is chosen to copy over an $Ab$ lineage. (Note that in this section we switch to the more familiar notation $i,j,k$ to denote counts of $AB$, $Ab$, and $ab$ haplotypes, respectively. This is analogous to the $c$-space notation in our derivation of the Hill-Robertson recursion, and the drift operator presented here is closely related to Equation S6.) For a given frequency bin $(i,j,k)$, this occurs if we first choose an $AB$ lineage that copies itself twice (with probability $i/n$), then choose an $Ab$ lineage to be copied over (with probability $j/(n-1)$), scaled by the density of two-locus haploypes with those frequencies ($\Psi_n(i,j,k)$). Together we get

$$P_{1\to 2}((i,j,k) \to (i+1,j-1,k)) = \frac{i}{n}\frac{j}{n-1}\Psi_n(i,j,k). \tag{S18}$$

We account for all possible changes in haplotype frequencies due to copying events in this way:

$$
\begin{aligned}
P_{1\to 2}((i,j,k) \to (i+1,j,k)) &= \frac{i}{n}\frac{n-i-j-k}{n-1}\Psi_n(i,j,k) \\
P_{1\to 2}((i,j,k) \to (i-1,j,k)) &= \frac{i}{n}\frac{n-i-j-k}{n-1}\Psi_n(i,j,k) \\
P_{1\to 2}((i,j,k) \to (i,j+1,k)) &= \frac{j}{n}\frac{n-i-j-k}{n-1}\Psi_n(i,j,k) \\
P_{1\to 2}((i,j,k) \to (i,j-1,k)) &= \frac{j}{n}\frac{n-i-j-k}{n-1}\Psi_n(i,j,k) \\
P_{1\to 2}((i,j,k) \to (i,j,k+1)) &= \frac{k}{n}\frac{n-i-j-k}{n-1}\Psi_n(i,j,k) \\
P_{1\to 2}((i,j,k) \to (i,j,k-1)) &= \frac{k}{n}\frac{n-i-j-k}{n-1}\Psi_n(i,j,k) \\
P_{1\to 2}((i,j,k) \to (i+1,j-1,k)) &= \frac{i}{n}\frac{j}{n-1}\Psi_n(i,j,k) \\
P_{1\to 2}((i,j,k) \to (i+1,j,k-1)) &= \frac{i}{n}\frac{k}{n-1}\Psi_n(i,j,k) \\
P_{1\to 2}((i,j,k) \to (i-1,j+1,k)) &= \frac{i}{n}\frac{j}{n-1}\Psi_n(i,j,k) \\
P_{1\to 2}((i,j,k) \to (i-1,j,k+1)) &= \frac{i}{n}\frac{k}{n-1}\Psi_n(i,j,k) \\
P_{1\to 2}((i,j,k) \to (i,j+1,k-1)) &= \frac{j}{n}\frac{k}{n-1}\Psi_n(i,j,k) \\
P_{1\to 2}((i,j,k) \to (i,j-1,k+1)) &= \frac{j}{n}\frac{k}{n-1}\Psi_n(i,j,k)
\end{aligned}
$$

$$
\begin{aligned}
P_{1\to 2}((i-1,j,k) \to (i,j,k)) &= \frac{i-1}{n}\frac{n-i-j-k+1}{n-1}\Psi_n(i-1,j,k)\delta_{i>0} \\
P_{1\to 2}((i+1,j,k) \to (i,j,k)) &= \frac{i+1}{n}\frac{n-i-j-k-1}{n-1}\Psi_n(i+1,j,k)\delta_{i<n} \\
P_{1\to 2}((i,j-1,k) \to (i,j,k)) &= \frac{j-1}{n}\frac{n-i-j-k+1}{n-1}\Psi_n(i,j-1,k)\delta_{j>0} \\
P_{1\to 2}((i,j+1,k) \to (i,j,k)) &= \frac{j+1}{n}\frac{n-i-j-k-1}{n-1}\Psi_n(i,j+1,k)\delta_{j<n} \\
P_{1\to 2}((i,j,k-1) \to (i,j,k)) &= \frac{k-1}{n}\frac{n-i-j-k+1}{n-1}\Psi_n(i,j,k-1)\delta_{k>0} \\
P_{1\to 2}((i,j,k+1) \to (i,j,k)) &= \frac{k+1}{n}\frac{n-i-j-k-1}{n-1}\Psi_n(i,j,k+1)\delta_{k<n} \\
P_{1\to 2}((i-1,j+1,k) \to (i,j,k)) &= \frac{i-1}{n}\frac{j+1}{n-1}\Psi_n(i-1,j+1,k)\delta_{i>0} \\
P_{1\to 2}((i-1,j,k+1) \to (i,j,k)) &= \frac{i-1}{n}\frac{k+1}{n-1}\Psi_n(i-1,j,k+1)\delta_{i>0} \\
P_{1\to 2}((i+1,j-1,k) \to (i,j,k)) &= \frac{i+1}{n}\frac{j-1}{n-1}\Psi_n(i+1,j-1,k)\delta_{j>0} \\
P_{1\to 2}((i+1,j,k-1) \to (i,j,k)) &= \frac{i+1}{n}\frac{k-1}{n-1}\Psi_n(i+1,j,k-1)\delta_{k>0} \\
P_{1\to 2}((i,j-1,k+1) \to (i,j,k)) &= \frac{j-1}{n}\frac{k+1}{n-1}\Psi_n(i,j-1,k+1)\delta_{j>0} \\
P_{1\to 2}((i,j+1,k-1) \to (i,j,k)) &= \frac{j+1}{n}\frac{k-1}{n-1}\Psi_n(i,j+1,k-1)\delta_{k>0}
\end{aligned}
$$

Taken together, we obtain $\mathcal{D}_n(i,j,k)$.

##### S1.3.2 Mutation

The moment system for $\Psi_n$ allows for flexible mutation operators. Both the infinite sites model (ISM) and a reversible mutation model are straightforward to derive and implement.

For the ISM model, mutations at each locus occur only once from ancestral to derived state. We suppose the mutation rate at the left locus is $u_1$ (for $a \to A$) and at the right locus is $u_2$ ($b \to B$). For a two-locus pair to segregate at both loci, a mutation must first occur at one of the loci, and then a mutation must occur at the second locus while the first locus is still segregating. For the first mutation, we introduce density in the singleton bins (one copy of either $Ab$ or $aB$) as

$$\mathcal{U}\Psi_n(0,1,0) = n\,u_1, \tag{S19}$$
$$\mathcal{U}\Psi_n(0,0,1) = n\,u_2. \tag{S20}$$

The second mutation, which occurs while the first locus is already segregating, introduces pairs of segregating loci proportional to the marginal single locus AFS at each locus. We account for whether, for example, a mutation $a \to A$ occurs on a $B$ or $b$ background:

$$\mathcal{U}\Psi_n(1,j,0) = u_2(j+1)\Psi_n(0,j+1,0) \qquad \forall j \in \{0,1,\dots,n-1\} \tag{S21}$$
$$\mathcal{U}\Psi_n(0,j,1) = u_2(n-j)\Psi_n(0,j,0) \qquad \forall j \in \{1,2,\dots,n\} \tag{S22}$$
$$\mathcal{U}\Psi_n(1,0,k) = u_1(k+1)\Psi_n(0,0,k+1) \qquad \forall k \in \{0,1,\dots,n-1\} \tag{S23}$$
$$\mathcal{U}\Psi_n(0,1,k) = u_1(n-k)\Psi_n(0,0,k) \qquad \forall k \in \{1,2,\dots,n\}. \tag{S24}$$

We can allow recurrent, reversible mutations, with rates

$$a \underset{v_1}{\overset{u_1}{\rightleftharpoons}} A \text{ and } b \underset{v_2}{\overset{u_2}{\rightleftharpoons}} B. \tag{S25}$$

In this case, there are no absorbing states. For example, the probability that a mutation event $a \to A$ occurs ($u_1$) that changes an $aB$ haplotype to an $AB$ haplotype depends on the number of $aB$ haplotypes present in the sample ($k$) and the probability of observing the required haplotype frequencies $\Psi_n(i,j,k)$. Then $P_{\text{mut}}((i,j,k) \to (i+1,j,k-1)) = u_1(k)\Psi_n(i,j,k)$. All together, the mutation operator is

$$\begin{aligned}
\mathcal{U}_{\text{rev}}\Psi_n(i,j,k) = {} & u_1(k+1)\Psi_n(i-1,j,k+1) - u_1(k)\Psi_n(i,j,k) \\
& + u_1(n-i-j-k+1)\Psi_n(i,j-1,k) - u_1(n-i-j-k)\Psi_n(i,j,k) \\
& + v_1(i+1)\Psi_n(i+1,j,k-1) - v_1(i)\Psi_n(i,j,k) \\
& + v_1(j+1)\Psi_n(i,j+1,k) - v_1(j)\Psi_n(i,j,k) \\
& + u_2(j+1)\Psi_n(i-1,j+1,k) - u_2(j)\Psi_n(i,j,k) \\
& + u_2(n-i-j-k+1)\Psi_n(i,j,k-1) - u_2(n-i-j-k)\Psi_n(i,j,k) \\
& + v_2(i+1)\Psi_n(i+1,j-1,k) - v_2(i)\Psi_n(i,j,k) \\
& + v_2(k+1)\Psi_n(i,j,k+1) - v_2(k)\Psi_n(i,j,k).
\end{aligned}$$

##### S1.3.3 Recombination

Here we derive the probabilities for transitions in frequencies due to recombination events, where a lineage in the sample is chosen to recombine with a lineage drawn from the full population, and one of the two recombinant types replaces the chosen lineage. Because we need to choose an extra lineage from the full population, $\mathcal{R}\Psi_n$ depends on $\Psi_{n+1}$, leading to the system being unclosed. For example, we write the probability of recombination between an $ab$ type lineage from within our $n$ subsampled lineages and an $AB$ haplotype from the full population, and subsequently changing frequencies from $(i,j,k)$ to $(i',j',k')$, as $\mathcal{R}_{ab\times AB}((i,j,k) \to (i',j',k'))$. These probabilities are

$$\begin{aligned}
\mathcal{R}_{ab\times AB}((i,j,k) \to (i,j+1,k)) &= \frac{1}{2}\Psi_{n+1}(i+1,j,k)\frac{n-i-j-k}{n+1}\frac{i+1}{n} \\
\mathcal{R}_{ab\times AB}((i,j,k) \to (i,j,k+1)) &= \frac{1}{2}\Psi_{n+1}(i+1,j,k)\frac{n-i-j-k}{n+1}\frac{i+1}{n} \\
\mathcal{R}_{ab\times Ab}((i,j,k) \to (i,j+1,k)) &= \frac{1}{2}\Psi_{n+1}(i,j+1,k)\frac{n-i-j-k}{n+1}\frac{j+1}{n} \\
\mathcal{R}_{ab\times aB}((i,j,k) \to (i,j,k+1)) &= \frac{1}{2}\Psi_{n+1}(i,j,k+1)\frac{n-i-j-k}{n+1}\frac{k+1}{n}
\end{aligned}$$

$$\mathcal{R}_{aB\times AB}((i,j,k)\to(i+1,j,k-1)) = \frac{1}{2}\Psi_{n+1}(i+1,j,k)\frac{k}{n+1}\frac{i+1}{n}$$
$$\mathcal{R}_{aB\times Ab}((i,j,k)\to(i+1,j,k-1)) = \frac{1}{2}\Psi_{n+1}(i,j+1,k)\frac{k}{n+1}\frac{j+1}{n}$$
$$\mathcal{R}_{aB\times Ab}((i,j,k)\to(i,j,k-1)) = \frac{1}{2}\Psi_{n+1}(i,j+1,k)\frac{k}{n+1}\frac{j+1}{n}$$
$$\mathcal{R}_{aB\times ab}((i,j,k)\to(i,j,k-1)) = \frac{1}{2}\Psi_{n+1}(i,j,k)\frac{k}{n+1}\frac{n-i-j-k+1}{n}$$
$$\mathcal{R}_{Ab\times AB}((i,j,k)\to(i+1,j-1,k)) = \frac{1}{2}\Psi_{n+1}(i+1,j,k)\frac{j}{n+1}\frac{i+1}{n}$$
$$\mathcal{R}_{Ab\times aB}((i,j,k)\to(i+1,j-1,k)) = \frac{1}{2}\Psi_{n+1}(i,j,k+1)\frac{j}{n+1}\frac{k+1}{n}$$
$$\mathcal{R}_{Ab\times aB}((i,j,k)\to(i,j-1,k)) = \frac{1}{2}\Psi_{n+1}(i,j,k+1)\frac{j}{n+1}\frac{k+1}{n}$$
$$\mathcal{R}_{Ab\times ab}((i,j,k)\to(i,j-1,k)) = \frac{1}{2}\Psi_{n+1}(i,j,k)\frac{j}{n+1}\frac{n-i-j-k+1}{n}$$
$$\mathcal{R}_{AB\times Ab}((i,j,k)\to(i-1,j,k)) = \frac{1}{2}\Psi_{n+1}(i,j,k)\frac{i}{n+1}\frac{j+1}{n}$$
$$\mathcal{R}_{AB\times aB}((i,j,k)\to(i-1,j,k)) = \frac{1}{2}\Psi_{n+1}(i,j,k)\frac{i}{n+1}\frac{k+1}{n}$$
$$\mathcal{R}_{AB\times ab}((i,j,k)\to(i-1,j,k)) = \frac{1}{2}\Psi_{n+1}(i,j+1,k)\frac{i}{n+1}\frac{n-i-j-k+1}{n}$$
$$\mathcal{R}_{AB\times ab}((i,j,k)\to(i-1,j,k)) = \frac{1}{2}\Psi_{n+1}(i,j,k+1)\frac{i}{n+1}\frac{n-i-j-k+1}{n}$$

and

$$\mathcal{R}_{ab\times AB}((i,j-1,k)\to(i,j,k)) = \frac{1}{2}\Psi_{n+1}(i+1,j-1,k)\frac{n-i-j-k+1}{n+1}\frac{i+1}{n}$$
$$\mathcal{R}_{ab\times AB}((i,j,k-1)\to(i,j,k)) = \frac{1}{2}\Psi_{n+1}(i+1,j,k-1)\frac{n-i-j-k+1}{n+1}\frac{i+1}{n}$$
$$\mathcal{R}_{ab\times Ab}((i,j-1,k)\to(i,j,k)) = \frac{1}{2}\Psi_{n+1}(i,j,k)\frac{n-i-j-k+1}{n+1}\frac{j}{n}$$
$$\mathcal{R}_{ab\times aB}((i,j,k-1)\to(i,j,k)) = \frac{1}{2}\Psi_{n+1}(i,j,k)\frac{n-i-j-k+1}{n+1}\frac{k}{n}$$
$$\mathcal{R}_{aB\times AB}((i-1,j,k+1)\to(i,j,k)) = \frac{1}{2}\Psi_{n+1}(i,j,k+1)\frac{k+1}{n+1}\frac{i}{n}$$
$$\mathcal{R}_{aB\times Ab}((i-1,j,k+1)\to(i,j,k)) = \frac{1}{2}\Psi_{n+1}(i-1,j+1,k+1)\frac{k+1}{n+1}\frac{j+1}{n}$$
$$\mathcal{R}_{aB\times Ab}((i-1,j,k+1)\to(i,j,k)) = \frac{1}{2}\Psi_{n+1}(i,j+1,k+1)\frac{k+1}{n+1}\frac{j+1}{n}$$
$$\mathcal{R}_{aB\times ab}((i,j,k+1)\to(i,j,k)) = \frac{1}{2}\Psi_{n+1}(i,j,k+1)\frac{k+1}{n+1}\frac{n-i-j-k}{n}$$
$$\mathcal{R}_{Ab\times AB}((i-1,j+1,k)\to(i,j,k)) = \frac{1}{2}\Psi_{n+1}(i,j+1,k)\frac{j+1}{n+1}\frac{i}{n}$$
$$\mathcal{R}_{Ab\times aB}((i-1,j+1,k)\to(i,j,k)) = \frac{1}{2}\Psi_{n+1}(i-1,j+1,k+1)\frac{j+1}{n+1}\frac{k+1}{n}$$
$$\mathcal{R}_{Ab\times aB}((i,j+1,k)\to(i,j,k)) = \frac{1}{2}\Psi_{n+1}(i,j+1,k+1)\frac{j+1}{n+1}\frac{k+1}{n}$$
$$\mathcal{R}_{Ab\times ab}((i,j+1,k)\to(i,j,k)) = \frac{1}{2}\Psi_{n+1}(i,j+1,k)\frac{j+1}{n+1}\frac{n-i-j-k}{n}$$
$$\mathcal{R}_{AB\times Ab}((i+1,j-1,k)\to(i,j,k)) = \frac{1}{2}\Psi_{n+1}(i+1,j,k)\frac{i+1}{n+1}\frac{j}{n}$$
$$\mathcal{R}_{AB\times aB}((i+1,j,k-1)\to(i,j,k)) = \frac{1}{2}\Psi_{n+1}(i+1,j,k)\frac{i+1}{n+1}\frac{k}{n}$$
$$\mathcal{R}_{AB\times ab}((i+1,j-1,k)\to(i,j,k)) = \frac{1}{2}\Psi_{n+1}(i+1,j-1,k)\frac{i+1}{n+1}\frac{n-i-j-k+1}{n}$$

$$\mathcal{R}_{AB\times ab}((i+1,j,k-1)\to(i,j,k)) = \frac{1}{2}\Psi_{n+1}(i+1,j,k-1)\frac{i+1}{n+1}\frac{n-i-j-k+1}{n}$$

Multiplying by the probability that a recombination event occurs on a lineage in our sample ($nr$, assuming $r \ll 1$), and then cancelling terms and simplifying, we find

$$\begin{aligned}
\mathcal{R}_{n,r}\Psi_n(i,j,k) = nr\times \Bigg[&\Psi_{n+1}(i+1,j-1,k)\frac{i+1}{n+1}\frac{n-i-j-k+1}{n} \\
&+ \Psi_{n+1}(i+1,j,k-1)\frac{i+1}{n+1}\frac{n-i-j-k+1}{n} \\
&+ \Psi_{n+1}(i-1,j+1,k+1)\frac{j+1}{n+1}\frac{k+1}{n} \\
&+ \Psi_{n+1}(i-1,j+1,k+1)\frac{j+1}{n+1}\frac{k+1}{n} \\
&- \Psi_{n+1}(i+1,j,k)\frac{i+1}{n+1}\frac{n-i-j-k}{n} \\
&- \Psi_{n+1}(i,j+1,k)\frac{j+1}{n+1}\frac{k}{n} \\
&- \Psi_{n+1}(i,j,k+1)\frac{j}{n+1}\frac{k+1}{n} \\
&-\Psi_{n+1}(i,j,k)\frac{i}{n+1}\frac{n-i-j-k+1}{n}\Bigg].
\end{aligned}$$

##### S1.3.4 Selection

Here, we consider a model that allows selection at the left locus ($A/a$). In this setting we model a neutral locus linked to a selected locus separated by arbitrary recombination rate. We suppose $A$ has selection and dominance coefficients $s$ (with $|s| \ll 1$) and $h$, so that a diploid carrying $AA$ has relative fitness $(1+s)$ compared to a $aa$ diploid, while the heterozygote $Aa$ has relative fitness $(1+hs)$. If $h = 1/2$, this reduces to the simple haploid selection model with $A$ having fitness $(1+s)$ relative to $a$.

We consider how selection is expected to change haplotype frequencies over a given generation. As was the case for recombination and drift events above, there are many possible haplotype frequency changes that may occur due to a selection event.

For example, suppose that selection acts against the $A$ allele ($s < 0$). We want to estimate the probability of events where, for example, a selection event occurs in which an $AB$ lineage fails to replicate to the subsequent generation and is replaced by an $Ab$ haplotype, drawn from the full population. The $AB$ lineage is eliminated with probability $-s$ if the parent is a homozygote at the $A/a$ locus (i.e. its diploid pair is type $AB$ or $Ab$), and it is eliminated with probability $-sh$ if a heterozygote (i.e. paired with $aB$ or $ab$). To compute the probability of any such event, we must draw two additional lineages from the population (one for the diploid pairing and one for the replacement lineage) in addition to the $n$ tracked lineages in $\Psi_n$. Thus, we require $\Psi_{n+2}$ to compute the evolution of $\Psi_n$.

Here, we write the probability that an $AB$ lineage paired with $aB$ is replaced by an $Ab$ lineage as $\mathcal{S}_{(aB,\mathbf{AB})\leftarrow Ab}$. This event has probability

$$\mathcal{S}_{(aB,\mathbf{AB})\leftarrow Ab} = -sh\cdot n\cdot\Psi_{n+2}(i,j+1,k+1)\frac{\binom{i}{1}\binom{j+1}{1}\binom{k+1}{1}}{\binom{n}{3}}\frac{1}{6}, \tag{S26}$$

where we account for choosing the three necessary lineages in the correct order. Probabilities for the other three possible diploid pairings take a similar form:

$$\mathcal{S}_{(AB,\mathbf{AB})\leftarrow Ab} = -s\cdot n\cdot\Psi_{n+2}(i+1,j+1,k+1)\frac{\binom{i+1}{2}\binom{j+1}{1}}{\binom{n}{3}}\frac{1}{3}$$

$$\mathcal{S}_{(Ab,\mathbf{AB})\leftarrow Ab} = -s\cdot n\cdot\Psi_{n+2}(i,j+2,k)\frac{\binom{i}{1}\binom{j+2}{2}}{\binom{n}{3}}\frac{1}{3}$$

$$\mathcal{S}_{(ab,\mathbf{AB})\leftarrow Ab} = -sh \cdot n \cdot \Psi_{n+2}(i,j+1,k)\frac{\binom{j+1}{1}\binom{k+1}{1}\binom{n-i-j-k+1}{1}}{\binom{n}{3}}\frac{1}{6}.$$

Accounting for all possible selection events (including the replaced lineage, its diploid pair, and the replacement lineage), we find the selection operator

$$\begin{aligned}\mathcal{S}_{n,s,h} =& s\Bigg\{\frac{h}{n+1}\Bigg[(i+j)(k+1)\Psi_{n+1}(i,j,k+1)+ \\ &+ (i+j)(n-i-j-k+1)\Psi_{n+1}(i,j,k) \\ &- (i+1)(n-i-j)\Psi_{n+1}(i+1,j,k) \\ &- (j+1)(n-i-j)\Psi_{n+1}(i,j+1,k)\Bigg] \\ &+ \frac{1-2h}{(n+2)(n+1)}\Bigg[(i+1)(k+1)(i+j)\Psi_{n+2}(i+1,j,k+1) \\ &+ (i+1)(n-i-j-k+1)(i+j)\Psi_{n+2}(i+1,j,k) \\ &+ (j+1)(k+1)(i+j)\Psi_{n+2}(i,j+1,k+1) \\ &+ (j+1)(n-i-j-k+1)(i+j)\Psi_{n+2}(i,j+1,k) \\ &- (i+2)(i+1)(n-i-j)\Psi_{n+2}(i+2,j,k) \\ &- 2(i+1)(j+1)(n-i-j)\Psi_{n+2}(i+1,j+1,k) \\ &- (j+2)(j+1)(n-i-j)\Psi_{n+2}(i,j+2,k)\Bigg]\Bigg\}.\end{aligned} \tag{S27}$$

Here we've used the downsampling formula to simplify the additive terms: if $h = 1/2$, $\mathcal{S}$ only requires $\Psi_{n+1}$ since selection is independent of the state of the paired lineage.

##### S1.3.5 Moment closure approximation for $\Psi_n$

Because neither recombination nor selection close in the full two-locus haplotype frequency system, we require a moment closure approximation to integrate Equation S16 forward in time. Here we will use a jackknife extrapolation, similar to Jouganous *et al.* (2017) for the single locus AFS, to express $\Psi_{n+2}$ and $\Psi_{n+1}$ as linear combinations of $\Psi_n$, so that

$$\Psi_{n+l}(i,j,k) \approx \sum_{(i',j',k')\in I_{(i,j,k)}} w_{(i',j',k')}\Psi_n(i',j',k'), \tag{S28}$$

where $l \in 1,2$ and the entries $(i',j',k') \in I_{(i,j,k)}$ are chosen so that $(i'/n, j'/n, k'/n)$ are close to $(i/(n+l), j/(n+l), k/(n+l))$. Each choice $l$ will have its own set $I$ and weights $w$.

We then find the appropriate set of entries $I$ and weights $w$, for a given $l$ and entry in that frequency spectrum. First, we note that for any continuous function $\psi$ that solves Equation S15, we can find the entries of $\Psi_n(i,j,k)$ for any $n,i,j,k$, using the multinomial sampling formula

$$\Psi_n(i,j,k) = \iiint_{\substack{x,y,z\geq 0,\\ x+y+z\leq 1}} \psi(x,y,z)\binom{n}{i,j,k}x^i y^j z^k(1-x-y-z)^{n-i-j-k}\,dx\,dy\,dz. \tag{S29}$$

We make the assumption here that $\psi$ can be approximated *locally* as a quadratic, so that

$$\psi(x,y,z) \approx a_1 + a_2x + a_3y + a_4z + a_5x^2 + a_6xy + a_7xz + a_8y^2 + a_9yz + a_{10}z^2. \tag{S30}$$

Using this approximation of $\psi$, the multinomial sampling integral can be computed analytically, to get

$$\tilde{\Psi}_n(i,j,k) = \frac{1}{(n+5)(n+4)(n+3)(n+2)(n+1)}\Big[a_1(n+5)(n+4) + a_2(n+5)(i+1) \tag{S31}$$

$$
\begin{aligned}
&+ a_3(n+5)(j+1) + a_4(n+5)(k+1) \\
&+ a_5(i+2)(i+1) + a_6(i+1)(j+1) \\
&+ a_7(i+1)(k+1) + a_8(j+2)(j+1) \\
&+ a_9(j+1)(k+1) + a_{10}(k+2)(k+1)\Big]
\end{aligned}
$$

For a given $\Psi_n(i,j,k)$ we take the ten closest values $I = \{(i'_1, j'_1, k'_1), (i'_2, j'_2, k'_2), \ldots, (i'_{10}, j'_{10}, k'_{10})\}$ as described above, with the added condition that the sets of $\{i'\}, \{j'\}$, and $\{k'\}$ each have size at least three. We then set

$$
\tilde{\Psi}_{n+l}(i,j,k) = \sum w_{(i',j',k')} \tilde{\Psi}_n(i',j',k'). \tag{S32}
$$

This holds for any $\psi$, so we set $(a_1, a_2, \ldots, a_{10}) = (1, 0, \ldots, 0)$, $(a_1, a_2, \ldots, a_{10}) = (0, 1, \ldots, 0)$, etc, in turn to get a system of equations for the weights $w_m = w_{(i'_m, j'_m, k'_m)}$, $m = 1, \ldots, 10$:

$$
\begin{cases}
\frac{(n+3)!(n+l)!}{n!(n+3+l)!} = \sum_{m=1}^{10} w_m \\
\frac{(n+4)!(n+l)!}{n!(n+4+l)!} = \sum_{m=1}^{10} (i'_m + 1) w_m \\
\frac{(n+4)!(n+l)!}{n!(n+4+l)!} = \sum_{m=1}^{10} (j'_m + 1) w_m \\
\frac{(n+4)!(n+l)!}{n!(n+4+l)!} = \sum_{m=1}^{10} (k'_m + 1) w_m \\
\frac{(n+5)!(n+l)!}{n!(n+5+l)!} = \sum_{m=1}^{10} (i'_m + 2)(i'_m + 1) w_m \\
\frac{(n+5)!(n+l)!}{n!(n+5+l)!} = \sum_{m=1}^{10} (i'_m + 1)(j'_m + 1) w_m \\
\frac{(n+5)!(n+l)!}{n!(n+5+l)!} = \sum_{m=1}^{10} (i'_m + 1)(k'_m + 1) w_m \\
\frac{(n+5)!(n+l)!}{n!(n+5+l)!} = \sum_{m=1}^{10} (j'_m + 2)(j'_m + 1) w_m \\
\frac{(n+5)!(n+l)!}{n!(n+5+l)!} = \sum_{m=1}^{10} (j'_m + 1)(k'_m + 1) w_m \\
\frac{(n+5)!(n+l)!}{n!(n+5+l)!} = \sum_{m=1}^{10} (k'_m + 2)(k'_m + 1) w_m,
\end{cases}
$$

which can be solved either analytically or numerically.

###### S1.3.6 Comparing methods to compute $\Psi_n$

We compared the accuracy and computational time needed for computing $\Psi_n$ using the numerical PDE approach presented in Ragsdale and Gutenkunst (2017) and the moment approach presented here. We performed this comparison for $n \in \{30, 50\}$ and $\rho \in \{0, 10\}$. We considered two demographic models for each sample size and recombination rate: equilibrium demography with no size changes, and a bottleneck demography. In the bottleneck demographic model the population is initially at steady state, then instantaneously changes in size to 1/10 the original size for 0.05 time units (measured in $2N_e$), and then it recovers to the original size for 0.2 time units.

For the equilibrium $\Psi_n$, we compared to Hudson's Monte Carlo implementation (described in Hudson (2001) and available from the author's website) for $n = 100$ projected down to sample sizes $n = 30$ and $n = 50$. We projected from a larger sample size for improved accuracy in Hudson's estimate and considered this distribution the "true" distribution to compare against. For the bottleneck distribution, we computed a numerical approximation with a larger sample size ($n = 80$) and 100x smaller time step than the default using `moments.TwoLocus` and then projected to $n = 30$ and $n = 50$. To compute $\Psi_n$ in ∂a∂i, we numerically solve for $\Psi_n$ for three grid spacings and three time steps, and then perform Richardson extrapolation (detailed in Ragsdale and Gutenkunst (2017)). We set integration time steps to $[0.005, 0.0025, 0.001]$ and grid points $[40, 50, 60]$ for $n = 30$ and $[60, 70, 80]$ for $n = 50$. We measured accuracy as $\sum (\Psi_n(\text{model}) - \Psi_n(\text{True}))^2 / \Psi_n(\text{True})$. In general, `moments` performs favorably compared to ∂a∂i, with orders of magnitude improved accuracy and faster integration and evaluation (Table S1).

##### S1.4 Deriving moment equations from the PDE

So far we have derived recursion equations in both the Hill-Robertson basis and for the full haplotype frequency spectrum by tracking an appropriately sized subset of lineages within the full population and considering the effects such as drift and recombination within these lineages. The statistics in these recursion equations are all non-canonical moments of the full two-locus distribution. Thus an alternative route to deriving all of the recursion equations presented in this paper is directly through the partial differential equation (PDE) describing the evolution of this full distribution.

Classically, two equivalent PDEs describing this distribution were studied, one in the variables of haplotype frequencies $(x_1, x_2, x_3, x_4)$ and the other in the variables $(p, q, D)$ (Hill and Robertson, 1966; Ohta and Kimura, 1969b). In this section, we outline the approach to obtain the Hill-Robertson $D^2$ system from the latter of these bases, do the same for the haplotype frequency spectrum $\Psi_n$, and then intuitively discuss why recombination closes in the Hill-Robertson basis but not the haplotype frequency basis from a PDE perspective.

Without selection, the neutral two-locus distribution $\psi(p, q, D)$ follows the forward Kolmogorov equation,

$$\begin{aligned}\frac{\partial \psi}{\partial \tau} =& \frac{1}{2}\frac{\partial^2}{\partial p^2}\left(p(1-p)\psi\right) + \frac{1}{2}\frac{\partial^2}{\partial q^2}\left(q(1-q)\psi\right) + \frac{\partial^2}{\partial p \partial q}\left(D\psi\right) + \frac{\partial^2}{\partial p \partial D}\left(D(1-2p)\psi\right) + \frac{\partial^2}{\partial q \partial D}\left(D(1-2q)\psi\right) \\ &+ \frac{1}{2}\frac{\partial^2}{\partial D^2}\left(\left(p(1-p)q(1-q) + D(1-2p)(1-2q) - D^2\right)\psi\right) + \frac{\partial}{\partial D}\left(D\left(1+\frac{\rho}{2}\right)\psi\right), \end{aligned} \tag{S33}$$

where time $\tau$ is measured in $2N$ generations. To obtain the time evolution of any moment of this distribution, we take

$$\begin{aligned}\partial_\tau \mathbb{E}[f(p, q, D)] &= \frac{\partial}{\partial \tau}\int \psi f(p, q, D) \\ &= \int \frac{\partial \psi}{\partial \tau} f(p, q, D) \\ &= \int (\text{RHS}) f(p, q, D),\end{aligned}$$

where RHS are the terms in Equation S33. Here, we have abused the integral notation, and it should be understood to be the triple integral over the domain of the function $\psi$.

For the $D^2$ system, we first find $\partial_\tau \mathbb{E}[D^2]$:

$$\begin{aligned}\partial_\tau \mathbb{E}[D^2] =& \frac{1}{2}\int D^2 \frac{\partial^2}{\partial p^2} p(1-p)\psi + \frac{1}{2}\int D^2 \frac{\partial^2}{\partial q^2} q(1-q)\psi + \int D^2 \frac{\partial}{\partial p \partial q} pq\psi \\ &+ \int D^2 \frac{\partial^2}{\partial p \partial D} D(1-2p)\psi + \int D^2 \frac{\partial^2}{\partial p \partial D} D(1-2p)\psi \\ &+ \frac{1}{2}\int D^2 \frac{\partial^2}{\partial D^2}\left(p(1-p)q(1-q) + D(1-2p)(1-2q) - D^2\right)\psi \\ &+ \int D^2 \frac{\partial}{\partial D} D\left(1+\frac{\rho}{2}\right)\psi \\ \overset{\text{IBP}}{=}& \int \left(p(1-p)q(1-q) + D(1-2p)(1-2q) - D^2\right)\psi - 2\int D^2\left(1+\frac{\rho}{2}\right)\psi \\ =& -3\mathbb{E}[D^2] + \mathbb{E}[D(1-2p)(1-2q)] + \mathbb{E}[p(1-p)q(1-q)] - \rho\mathbb{E}[D^2],\end{aligned}$$

which recovers, to first order, the Hill-Robertson equation for $\mathbb{E}[D^2]$. Surface terms vanish since the functions decay to zero at the boundary. The other two terms in the system can by found by similarly integrating by parts:

$$\begin{aligned}\partial_\tau \mathbb{E}[D(1-2p)(1-2q)] &= 4\mathbb{E}[D^2] - 5\mathbb{E}[D(1-2p)(1-2q)] - \frac{\rho}{2}\mathbb{E}[D(1-2p)(1-2q)] \\ \partial_\tau \mathbb{E}[p(1-p)q(1-q)] &= \mathbb{E}[D(1-2p)(1-2q)] - 2\mathbb{E}[p(1-p)q(1-q)]\end{aligned}$$

An equivalent PDE for $\psi$ is expressed in haplotype frequencies $(x_1, x_2, x_3)$:

$$\frac{\partial \psi}{\partial \tau} = \frac{1}{2}\sum_{i=1}^{3}\sum_{j=1}^{3}\frac{\partial^2}{\partial x_i \partial x_j} x_i(\delta_{ij} - x_j)\psi + \frac{\rho}{2}\left(\frac{\partial}{\partial x_1}D\psi - \frac{\partial}{\partial x_2}D\psi - \frac{\partial}{\partial x_3}D\psi\right), \tag{S34}$$

where $\delta_{ij}$ is the Kronecker delta function, and formally $D = x_1(1 - x_1 - x_2 - x_3) - x_2 x_3$. The first set of terms in the double sum account for drift, and the second set of terms account of recombination with scaled rate $\rho$.

We want to find evolution equations for $\Psi_n(i,j,k)$, and we take a similar approach as above. Given the continuous distribution $\psi$, we compute $\Psi_n(i,j,k)$ by integrating $\psi$ against the multinomial distribution:

$$\Psi_n(i,j,k) = \iiint \binom{n}{i,j,k,n-i-j-k} x_1^i x_2^j x_3^k (1 - x_1 - x_2 - x_3)^{n-i-j-k} \psi(x_1, x_2, x_3),$$

where $\binom{n}{i,j,k,n-i-j-k} = \frac{n!}{i!j!k!(n-i-j-k)!}$ is the multinomial coefficient. Then integrating both sizes of Equation S34 against this sampling function, we can find

$$\partial_\tau \Psi_n(i,j,k) = \iiint \binom{n}{i,j,k,n-i-j-k} x_1^i x_2^j x_3^k (1 - x_1 - x_2 - x_3)^{n-i-j-k}$$
$$\left( \frac{1}{2} \sum_{i=1}^{3} \sum_{j=1}^{3} \frac{\partial^2}{\partial x_i \partial x_j} x_i(\delta_{ij} - x_j)\psi + \frac{\rho}{2} \left( \frac{\partial}{\partial x_1} D\psi - \frac{\partial}{\partial x_2} D\psi - \frac{\partial}{\partial x_3} D\psi \right) \right).$$

In brief, for the drift terms we integrate by parts twice, obtain a series of multinomial sampling functions against $\psi$, and then simplify. This results in the same set of equations as derived above by computing copying probabilities. This derivation follows closely the approach described in Jouganous *et al.* (2017) for the single site AFS.

To understand closure properties, we can consider whether the order of terms on the RHS is always equal or smaller to $n$. If so, the set of all moments of order $n$ will only ever require moments of order less than or equal to $n$, ensuring closure of the moment equation. When counting the order of terms on the RHS, we must subtract one for each derivative: in the integration by part, each derivative reduces the degree of the polynomial coefficient by at least one. Thus the drift term on the RHS has order $n$ and closes, whereas the recombination term has order $n+1$, and as a result does not close.

##### S1.4.1 Closure of Hill-Robertson moments

Any order moment system for $\mathbb{E}[D^m]$ closes under drift and recombination. First, we observe that the recombination transition matrix will be diagonal by considering the PDE without the drift terms: $\psi_\tau = \frac{\rho}{2}(D\psi)_D$. For any moment $\mathbb{E}[D^\alpha f(p,q)]$, we get

$$\partial_\tau \mathbb{E}[D^\alpha f(p,q)] = \frac{\rho}{2} \int D^\alpha f(p,q) \frac{\partial}{\partial D} D\psi$$
$$= -\alpha \frac{\rho}{2} \int D^\alpha f(p,q)\psi = -\alpha \frac{\rho}{2} \mathbb{E}[D^\alpha f(p,q)].$$

Thus any basis of moments expressed as a functions of $(p,q,D)$ will close under recombination.

Intuitively, for any moment $f(p,q,D)$ we can expect to find a closed system for its evolution under drift. This can be seen from the PDE for $\psi$ (Equation S33): the coefficient of each spatial derivative has order equal to the derivative. Thus integrating by parts does not result in any moments of $\psi$ of higher order than the original moment $f$. Since there are a finite number of moments of any given order, this system must also be finite in size, and thus necessarily close.

Here we compute the recursion for the $D^m$ system (for $m$ even) and explicitly demonstrate that it closes by considering the new terms required to compute $\mathbb{E}[D^m]$. Again, we define $\pi_2 = p(1-p)q(1-q)$, $z = (1-2p)(1-2q)$, and $\sigma_j = p^j(1-p)^j + q^j(1-q)^j$.

First, suppose that the $D^{m-2}$ system closes. The $D^{m-2}$ system contains all terms

$$\left\{ \mathbb{E}[D^{m-2i-2j-k-2} \pi_2^i z^k \sigma_j] \right\}, \text{ for } k \in \{0,1\}, j \in \left\{0, 1, \ldots, \frac{m-2k-2}{2}\right\}, i \in \left\{0, 1, \ldots, \frac{m-2k-2j-2}{2}\right\}.$$

It also contains all terms in the $D^{m-4}$ system, which take the same form as above, and so on. By computing $\partial_\tau \mathbb{E}[D^m]$ and its dependent terms, we find that all terms are present in the $D^{m-2}$ system or are in

$$\left\{ \mathbb{E}[D^{m-2i-2j-k} \pi_2^i z^k \sigma_j] \right\}, \text{ for } k \in \{0,1\}, j \in \left\{0, 1, \ldots, \frac{m-2k}{2}\right\}, i \in \left\{0, 1, \ldots, \frac{m-2k-2j}{2}\right\}. \quad \text{(S35)}$$

For the $m^{\text{th}}$ moment of $D$, we can use this same approach to obtain moment dependencies and compute evolution equations, just as we did in the previous section for the Hill-Robertson system. For $\mathbb{E}[D^m]$ (here showing just the terms for drift), by integrating by parts, we have

$$\begin{aligned}\partial_\tau \mathbb{E}[D^m] &= \frac{1}{2}\int D^m \frac{\partial^2}{\partial D^2}\left(\pi_2 + Dz - D^2\right)\psi + \int D^m \frac{\partial}{\partial D} D\psi \\ &= \frac{m(m-1)}{2}\int D^{m-2}\left(\pi_2 + Dz - D^2\right)\psi - m\int D^m \psi \\ &= -\frac{m(m+1)}{2}\mathbb{E}[D^m] + \frac{m(m-1)}{2}\mathbb{E}[D^{m-1}z] + \frac{m(m-1)}{2}\mathbb{E}[D^{m-2}\pi_2].\end{aligned}$$

In the same way, we compute the time derivative for each dependent moment:

$$\begin{aligned}\partial_\tau \mathbb{E}[D^{m-1}z] &= 4\mathbb{E}[D^m] - \frac{(m+8)(m-1)}{2}\mathbb{E}[D^{m-1}z] + 8(m-1)(m-2)\mathbb{E}[D^{m-2}\pi_2] \\ &\quad - 2(m-1)(m-2)\mathbb{E}[D^{m-2}\sigma_1] + \frac{(m-1)(m-2)}{2}\mathbb{E}[D^{m-2}] + \frac{(m-1)(m-2)}{2}\mathbb{E}[D^{m-3}\pi_2 z] \\ \partial_\tau \mathbb{E}[D^{m-2}\pi_2] &= \mathbb{E}[D^{m-1}z] - \frac{1}{2}(m^2+13m-26)\mathbb{E}[D^{m-2}\pi_2] + (m-2)\mathbb{E}[D^{m-2}\sigma_1] \\ &\quad + \frac{(m-2)(m-3)}{2}\mathbb{E}[D^{m-3}\pi_2 z] + \frac{(m-2)(m-3)}{2}\mathbb{E}[D^{m-4}\pi_2^2], \\ &\ \ \vdots \\ \partial_\tau \mathbb{E}[D^{m-2i}\pi_2^i] &= i^2\mathbb{E}[D^{m-2i+1}\pi_2^{i-1}z] - \frac{m^2+(1+12i)m-2i(3+10i)}{2}\mathbb{E}[D^{m-2i}\pi_2^i] \\ &\quad (im - i(1+3i)/2)\mathbb{E}[D^{m-2i}\pi_2^{i-1}\sigma_1] + \frac{(m-i+1)(m-i)}{2}\mathbb{E}[D^{m-2i-1}\pi_2^i z] \\ &\quad + \frac{(m-i+1)(m-i)}{2}\mathbb{E}[D^{m-2i-2}\pi_2^{i+1}] \\ \partial_\tau \mathbb{E}[D^{m-2i-1}\pi_2^i z] &= 4(1+2i)^2\mathbb{E}[D^{m-2i}\pi_2^i] - 2i(1+2i)\mathbb{E}[D^{m-2i}\pi_2^{i-1}z] + i^2\mathbb{E}[D^{m-2i}\pi_2^{i-1}] \\ &\quad - \frac{m^2+(7+12i)m-2(10i^2+13i+4)}{2}\mathbb{E}[D^{m-2i-1}\pi_2^i z] \\ &\quad + (im - 3i(i+1)/2)\mathbb{E}[D^{m-2i-1}\pi_2^{i-1}z\sigma_1] + 8(m-2i-2)(m-2i-1)\mathbb{E}[D^{m-2i-2}\pi_2^{i+1}] \\ &\quad - 2(m-2i-2)(m-2i-1)\mathbb{E}[D^{m-2i-2}\pi_2^i\sigma_1] + \frac{(m-2i-2)(m-2i-1)}{2}\mathbb{E}[D^{m-2i-2}\pi_2^i] \\ &\quad + \frac{(m-2i-2)(m-2i-1)}{2}\mathbb{E}[D^{m-2i-3}\pi_2^{i+1}z] \\ &\ \ \vdots \\ \partial_\tau \mathbb{E}[D\pi_2^{\frac{m-2}{2}}z] &= \frac{(m-1)^2}{4}\mathbb{E}[D^2\pi_2^{\frac{m-2}{2}}] - (m-2)(m-1)\mathbb{E}[D^2\pi_2^{\frac{m-4}{2}}\sigma_1] + \frac{(m-2)^2}{4}\mathbb{E}[D^2\pi_2^{\frac{m-4}{2}}] \\ &\quad - (m^2+m-1)\mathbb{E}[D\pi_2^{\frac{m-2}{2}}z] + \frac{m(m-2)}{2}\mathbb{E}[D\pi_2^{\frac{m-4}{2}}z\sigma_1] \\ \partial_\tau \mathbb{E}[\pi_2^{\frac{m}{2}}] &= \frac{m^2}{4}\mathbb{E}[D\pi_2^{\frac{m-2}{2}}z] - m(m-1)\mathbb{E}[\pi_2^{\frac{m}{2}}] + \frac{m(m-2)}{8}\mathbb{E}[\pi_2^{\frac{m-2}{2}}\sigma_1]\end{aligned}$$

The evolution for the terms computed here require those already considered, terms in the $D^{m-2}$ system, or additional terms of the form

$$\{D^{m-2}\sigma_1, D^{m-3}z\sigma_1, D^{m-4}\pi_2\sigma_1, \ldots, \pi_2^{\frac{m-2}{2}}\sigma_1\}.$$

In general, for $j > 0$, $k \in \{0,1\}$, $i \in \{0,\ldots,(m-2j-2k)/2\}$, we have terms of the form $\mathbb{E}[D^{m-2i-2j-k}\pi_2^i z^k \sigma_j]$. We compute transition probabilities for each of these terms:

$$\begin{aligned}\partial_\tau \mathbb{E}[D^{m-2j}\sigma_j] = &-\frac{m^2+(4j+1)m-4i(1+2i)}{2}\mathbb{E}[D^{m-2j}\sigma_j] + (im + i(1+3i)/2)\mathbb{E}[D^{m-2j}\sigma_{j-1}] \\ &+ \frac{(m-2j-1)(m-2j)}{2}\mathbb{E}[D^{m-2j-1}z\sigma_j] + \frac{(m-2j-1)(m-2j)}{2}\mathbb{E}[D^{m-2j-2}\pi_2\sigma_j]\end{aligned}$$

$$
\begin{aligned}
\partial_\tau \mathbb{E}[D^{m-2j-1} z\sigma_j] = {} & 4(1+2i)\mathbb{E}[D^{m-2j}\sigma_j] - 2i\mathbb{E}[D^{m-2j}\sigma_{j-1}] \\
& - \frac{m^2 + (7+4i)m - 4(i + 2(1+i)^2)}{2}\mathbb{E}[D^{m-2j-1}z\sigma_j] \\
& + (4m - 3i(i+1)/2)\mathbb{E}[D^{m-2j-1}z\sigma_{j-1}] + 8(m-2j-2)(m-2j-1)\mathbb{E}[D^{m-2j-2}\pi_2\sigma_j] \\
& - 2(m-2j-2)(m-2j-1)\mathbb{E}[D^{m-2j-2}\pi_2\sigma_{j-1}] - 2(m-2j-2)(m-2j-1)\mathbb{E}[D^{m-2j-2}\sigma_{j+1}] \\
& + \frac{(m-2j-2)(m-2j-1)}{2}\mathbb{E}[D^{m-2j-2}\sigma_j] + \frac{(m-2j-2)(m-2j-1)}{2}\mathbb{E}[D^{m-2j-3}\pi_2 z\sigma_j] \\
& \vdots \\
\partial_\tau \mathbb{E}[D\pi_2^{\frac{m-2-2j}{2}} z\sigma_j] = {} & 4(m-2j-1)(m-1)\mathbb{E}[D^2\pi_2^{\frac{m-2j-2}{2}}\sigma_j] - (m-2j-1)(m-2)\mathbb{E}[D^2\pi_2^{\frac{m-2j-2}{2}}\sigma_{j-1}] \\
& - (m-2j-2)(m-1)\mathbb{E}[D^2\pi_2^{\frac{m-2j-4}{2}}\sigma_{j+1}] + \frac{(m-2j-2)(m-2)}{4}\mathbb{E}[D^2\pi_2^{\frac{m-2j-4}{2}}\sigma_{j-1}] \\
& - (m^2 - (2j-1)m + (2j+1)(j-1))\mathbb{E}[D\pi_2^{\frac{m-2j-2}{2}}z\sigma_j] + \frac{m(m-2)}{8}\mathbb{E}[D\pi_2^{\frac{m-2j-2}{2}}z\sigma_{j-1}] \\
& + \frac{(m-2j-2)(m-2j)}{8}\mathbb{E}[D\pi_2^{\frac{m-2j-4}{2}}z\sigma_{j+1}] \\
\partial_\tau \mathbb{E}[\pi_2^{\frac{m-2j}{2}}\sigma_j] = {} & \frac{m(m-2j)}{4}\mathbb{E}[D\pi_2^{\frac{m-2j-2}{2}}z\sigma_j] - (m^2 - (2j+1)m + j(2j+1))\mathbb{E}[\pi_2^{\frac{m-2j-2}{2}}\sigma_j] \\
& + \frac{m(m-2)}{8}\mathbb{E}[\pi_2^{\frac{m-2j-2}{2}}\sigma_j] + \frac{(m-2j-2)(m-2j)}{8}\mathbb{E}[\pi_2^{\frac{m-2j-4}{2}}\sigma_{j+1}].
\end{aligned}
$$

Each term appearing here belongs to the $D^{m-2}$ system or is found in the set of new moments enumerated in Equation S35.

#### S1.5 Sampling bias and the relationship between $\Psi_n$ and Hill-Robertson statistics

$\mathbb{E}[D]$ is a two-haplotype statistic, meaning we require one phased diploid genome to estimate $D$ genome-wide. In practice, to estimate $D$ we count the number of times we observe a $(AB|ab)$ pairing, subtract the counts of observed $(Ab|aB)$ pairings, and normalize by the total number of two-locus pairs considered. From the two-locus haplotype frequency spectrum, computed under the appropriate per-base mutation rate,

$$
\mathbb{E}[D] = \frac{1}{2}\left(\Psi_2(1,0,0) - \Psi_2(0,1,1)\right).
$$

A more accurate estimate may be obtained by considering haplotype frequencies in a larger sample size than $n = 2$, although the estimate would then need to be corrected due to sampling bias. Alternatively, we can use hypergeometric projection to directly calculate an unbiased estimate of $\mathbb{E}[D]$ for $n > 2$ samples. A two-locus pair with sample size $n$ and observed haplotype counts $(n_{AB}, n_{Ab}, n_{aB}, n_{ab})$ contributes to $\mathbb{E}[D]$ by

$$
\frac{1}{2}\frac{\binom{n_{AB}}{1}\binom{n_{ab}}{1}}{\binom{n}{2}} - \frac{1}{2}\frac{\binom{n_{Ab}}{1}\binom{n_{aB}}{1}}{\binom{n}{2}} = \frac{n_{AB}n_{ab}}{n(n-1)} - \frac{n_{Ab}n_{aB}}{n(n-1)}. \tag{S36}
$$

This approach not only provides an unbiased estimate of $\mathbb{E}[D]$, but also allows us to compute $\mathbb{E}[D]$ over pairs of sites with different sample sizes (e.g. to account for missing data).

We can express $\mathbb{E}[D^2]$ or any other term in the Hill-Robertson system as a linear combination of entries in $\Psi_4$. For example,

$$
\begin{aligned}
\mathbb{E}[D^2] &= \mathbb{E}[(f_{AB}f_{ab} - f_{Ab}f_{aB})^2] \\
&= 2\left(\frac{1}{\binom{4}{2,0,0,2}}\Psi_4(2,0,0) + \frac{1}{\binom{4}{0,2,2,0}}\Psi_4(0,2,2) - \frac{2}{\binom{4}{1,1,1,1}}\Psi_4(1,1,1)\right) \\
&= \frac{1}{3}\left(\Psi_4(2,0,0) + \Psi_4(0,2,2) - \frac{1}{2}\Psi_4(1,1,1)\right).
\end{aligned} \tag{S37}
$$

The multinomial factors arise because the entries of $\Psi_n$ are unsorted configuration probabilities, while $\mathbb{E}[D^2]$ implies a particular order of drawn haplotypes. This implies that we may obtain an unbiased estimate for any quantity (such as $\mathbb{E}[D^2]$) from an arbitrary sample size $n$ through hypergeometric projection to the appropriate sample size. For example, in a sample of size $n$ a two-locus pair (with observed haplotype counts $(n_{AB}, n_{Ab}, n_{aB}, n_{ab})$) contributes to $\mathbb{E}[D^2]$

$$\frac{1}{3}\frac{\binom{n_{AB}}{2}\binom{n_{ab}}{2}}{\binom{n}{4}} + \frac{1}{3}\frac{\binom{n_{Ab}}{2}\binom{n_{aB}}{2}}{\binom{n}{4}} - \frac{1}{6}\frac{\binom{n_{AB}}{1}\binom{n_{Ab}}{1}\binom{n_{aB}}{1}\binom{n_{ab}}{1}}{\binom{n}{4}}. \tag{S38}$$

This allows for direct comparison between observed haplotype data and expectations from the model without having to correct for sample size bias. We discuss our approach for unphased data below in Data processing.

#### S1.6 Low coverage data

Low coverage sequencing data is known to miss a sizable proportion of low frequency variation, so that singleton and doubleton bins of the AFS may be significantly underestimated (Gravel *et al.*, 2011). This may bias demographic inference based on the AFS from low coverage data, particularly for recent population size or growth parameters. Low order LD statistics studied in this paper are less sensitive to low coverage data, because low frequency variants contribute relatively little to aggregate statistics in the Hill-Robertson system across the genome (Rogers, 2014). Rogers (2014) argued that $\sigma_d^2$ type statistics are insensitive to variants with low heterozygosity, and thus insensitive to low coverage data or sequencing error.

To confirm this claim with real data, we examined the effect of low coverage on 40 individuals from the CHB population in the 1000 Genomes data that were also sequenced at high (∼80x) coverage in (Lan *et al.*, 2017). By comparing statistics computed from intergenic regions in the same individuals between the two datasets, we can see if low coverage biases our estimates. In Figure S6, we compare $\sigma_d^2 = \mathbb{E}[D^2]/\mathbb{E}[\pi_2]$, $\mathbb{E}[Dz]/\mathbb{E}[\pi_2]$, and the folded single-site AFS. We find that the low order two-locus statistics are unaffected by low coverage data, but low frequency bins of the AFS are underestimated in the low coverage data (17.5% fewer singletons, 7% fewer doubletons).

#### S1.7 Data processing

##### S1.7.1 Intergenic data

We used data from all intergenic regions on autosomal chromosomes, as identified using the GRCh37 build from the Genome Reference Consortium. We chose intergenic regions to reduce possible biases in statistic estimation due to selection (Figure S10). We considered only keeping SNPs at least a given distance from the nearest gene, to further reduce selective effects, but that would come at the cost of more noise in the statistic estimates, as we would be left with fewer loci from which to estimate statistics. We chose to use all intergenic SNPs because there was little difference between statistics estimated using all intergenic loci and using all loci at least a given distance from genes (Figure S11).

##### S1.7.2 Recombination map and binning pairs by recombination distance

We considered all pairs of biallelic SNPs in intergenic regions separated by recombination distances $0.00001 \leq r < 0.002$, and binned pairs of SNPs with bin edges (0.00001, 0.00002, 0.00003, 0.00005, 0.00007, 0.0001, 0.0002, 0.0003, 0.0005, 0.0007, 0.001, 0.002). Recombination distances for each pair of SNPs were computed using the African American recombination map from Hinch *et al.* (2011).

##### S1.7.3 Computing LD statistics from unphased data and accounting for sampling bias

For a given pair of biallelic SNPs, we computed their contribution to the set of statistics in the multi-population Hill-Robertson system (for three populations: YRI, CHB, and CEU). We determined two-locus genotype counts within each population $(n_{AABB}, n_{AaBB}, n_{aaBB}, n_{AABb}, \ldots)$, as well as allele frequencies at the left and right loci within each population. Because we worked with genotype data instead of phased haplotypes, we used the estimator $\hat{D}$ of Weir (1979),

$$\hat{D} = \frac{1}{2n_d}\left(2n_{AABB} + n_{AABb} + n_{AaBB} + \frac{1}{2}n_{AaBb}\right) - \frac{n_A}{2n_d}\frac{n_B}{2n_d},$$

and define

$$\hat{z}_{j,k} = (1-2\hat{p}_j)(1-2\hat{q}_k).$$

We then computed the values $\hat{D}_i\hat{D}_j$, $\hat{D}_i\hat{z}_{jk}$, and so on from genotype data.

$\hat{D}$ is known to be a biased estimator due to finite sample size. To compare model predictions to data, we treated $\hat{D}$ as a statistic and computed unbiased predictions for it. In the large sample size limit, $\mathbb{E}[f(p,q,D)] = \mathbb{E}[f(\hat{p},\hat{q},\hat{D})]$.

To account for sample size bias for each statistic in each population $(n_1, n_2, \ldots)$, we adjusted expectations to match observed biased statistics by computing expected values for each statistic under the multinomial sampling process. For a given statistic, we converted the $\hat{D}$ statistic to genotype frequency space (e.g., using $\hat{D} = (g_1 + g_2/2 + g_3/2 + g_5/4) - (g_1 + g_2 + g_3 + g_4/2 + g_5/2 + g_6/2)(g_1 + g_2/2 + g_4 + g_5/2 + g_7 + g_8/2)$, where $(g_1, g_2, \ldots, g_9) = (n_{AABB}/n_d,\ n_{AABb}/n_d,\ n_{AAbb}/n_d,\ n_{AaBB}/n_d,\ n_{AaBb}/n_d,\ n_{Aabb}/n_d,\ n_{aaBB}/n_d,\ n_{aaBb}/n_d,\ n_{aabb}/n_d))$.

Then for each term in the expansion in $g$-space, we computed the expected sampling probabilities through the multinomial moment generating function (Weir, 1979). We then converted those adjusted sampling probabilities to expected haplotype probabilities, and we found that we could express these sampling probabilities in the Hill-Robertson basis. In the Hill-Robertson system, for a sample size of $n$ diploid genomes, we have

$$\begin{pmatrix}\mathbb{E}_n[\hat{D}^2]\\ \mathbb{E}_n[\hat{D}\hat{z}]\\ \mathbb{E}_n[\hat{\pi}_2]\\ \mathbb{E}_n[\hat{H}]\end{pmatrix} = \begin{pmatrix}\frac{(n^2-n+1)(n-1)}{n^3} & \frac{(n-1)^2}{2n^3} & \frac{n-1}{n^2} & 0\\ \frac{2(n-1)^2}{n^3} & \frac{(n-1)^3}{n^3} & 0 & 0\\ \frac{(2n-1)}{4n^3} & \frac{(2n-1)^2}{8n^3} & \frac{(2n-1)^2}{4n^2} & 0\\ 0 & 0 & 0 & \frac{2n-1}{2n}\end{pmatrix}\begin{pmatrix}\mathbb{E}[D^2]\\ \mathbb{E}[Dz]\\ \mathbb{E}[\pi_2]\\ \mathbb{E}[H]\end{pmatrix},$$

where $\mathbb{E}_n$ denotes expected $\hat{D}$ statistics corrected for sample size bias for a given $n$. Here, we have taken the approach of computing predictions from the model and then adjusting those predictions through multinomial sampling theory, giving us predictions for biased estimates from the data. We could instead invert this matrix to compute unbiased estimates from data to compare directly to model predictions. While either approach should be acceptable, we chose to build the bias into the model to avoid any potential instability in solving the inverse problem.

In practice, we worked with $\sigma_d^2$-type statistics of the form $\mathbb{E}[\cdot]/\mathbb{E}[\pi_2(\text{YRI})]$ and compared to the same statistics computed from `moments.LD`. To compute $\mathbb{E}[D^2]$ or any other statistic for a given recombination bin, we sum all contributions of pairs of SNPs with recombination distance falling within that bin, and then divide by the total number of pairs of sites along the genome which could have contributed to that bin, whether they are variable or not. For $\sigma_d^2$-type statistics, we don't need to compute the total possible number of pairs per bin, and need only sum all contributions and divide by the total sum of contributions to $\mathbb{E}[\pi_2(\text{YRI})]$.

###### S1.7.4 Bootstraps for likelihood computation

We used a multivariate normal function to estimate likelihood from expected values. For a given recombination bin, we compared the observed vector of LD statistics to expectations computed under the model with adjustments for phasing and sample size. To compute likelihoods, we needed to estimate the covariances of statistics in this vector. To compute the covariance matrix, we divided the genome into 500 regions, each with approximately the same length of intergenic regions. We computed statistics over each of the 500 regions, and then constructed 500 bootstrap replicate sets of statistics by sampling with replacement 500 times. We used these bootstrap replicates to estimate the covariance matrix $\Sigma$ for each recombination bin. This same set of bootstrap replicates and covariance matrix was used to estimate confidence intervals, as proposed by Coffman *et al.* (2016), to account for non-independence of pairs of SNPs in the same region or pairs with overlapping SNPs.

#### Supplementary figures and tables

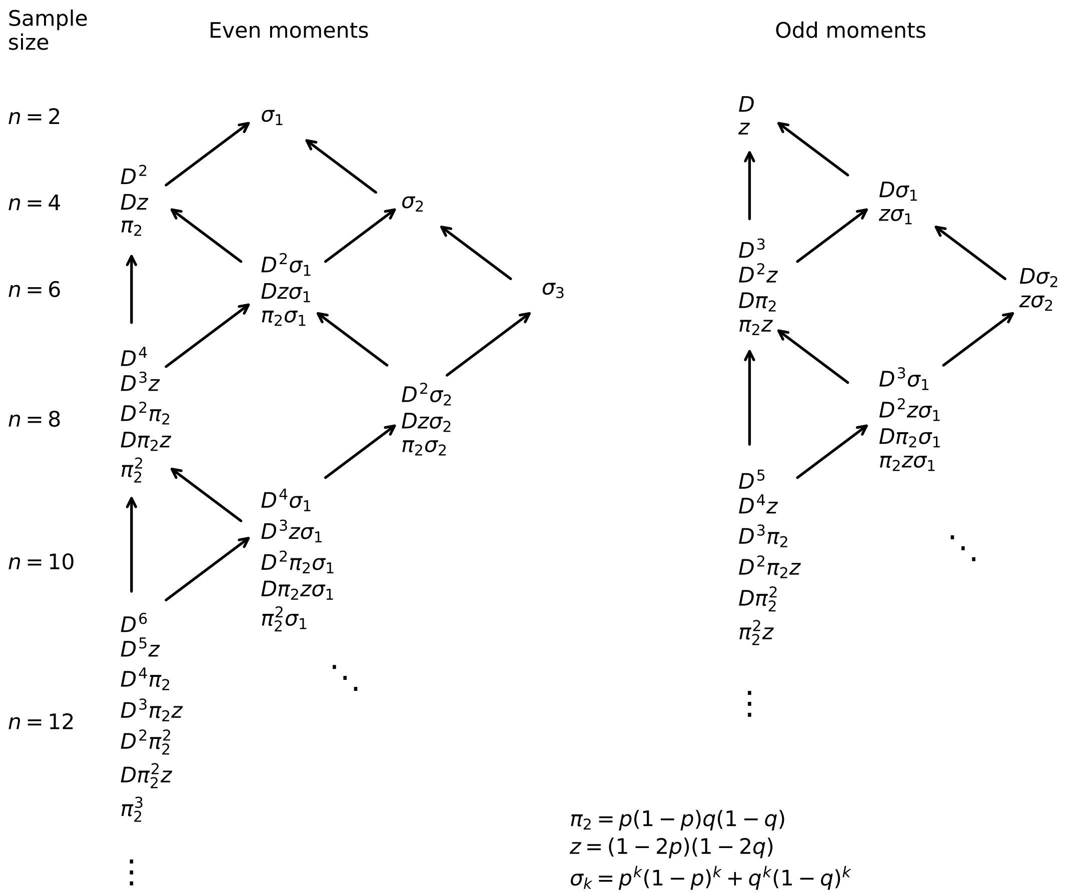

Figure S1: **Hierarchy of moments.** Even and odd moments separate into distinct hierarchical systems. Arrows indicate dependence of sets of moments.

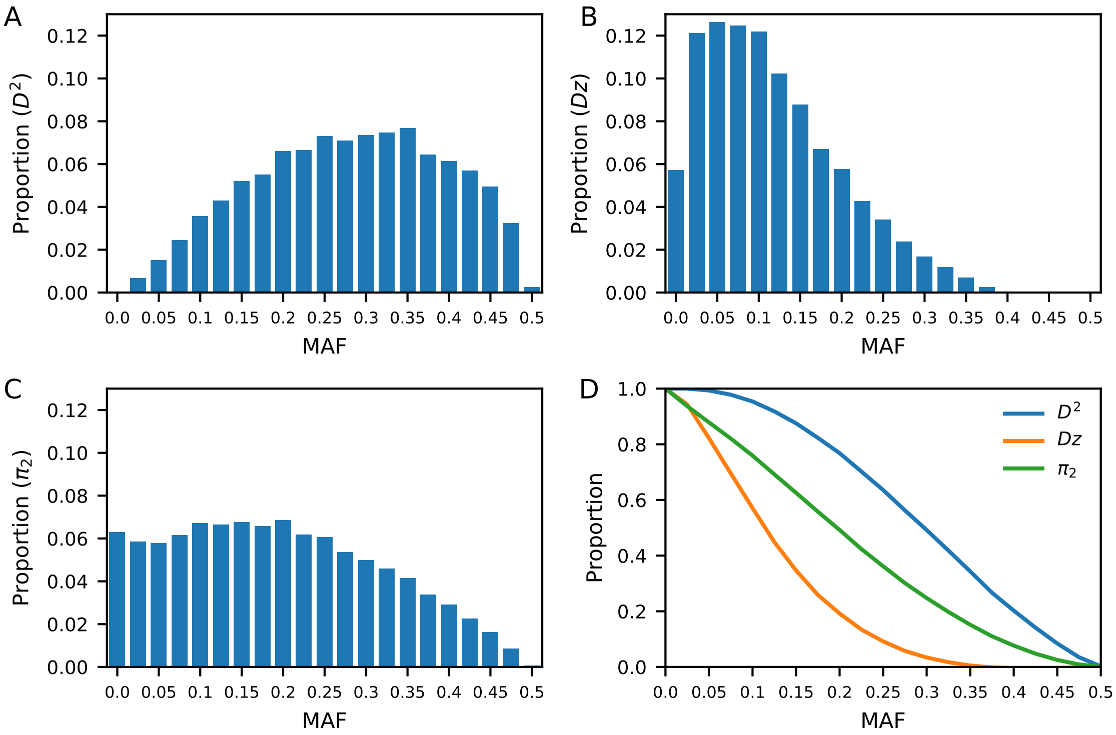

Figure S2: **Dependence of Hill-Robertson statistics on lowest minor allele frequency.** (A-C) The proportion of each statistic contributed by pairs of loci binned by the lowest MAF of the two loci. $D^2$ has contribution mainly from common variants (95% interval 0.1-0.5), while $Dz$ has contribution mainly from rare to low frequency variants (95% interval 0-0.35). (D) The proportion of each statistic contributed by pairs of alleles where both alleles have frequency above a given MAF. We computed these distributions from 40 high coverage CHB samples from Lan *et al.* (2017) that overlap with the 1000 Genomes Project Consortium *et al.* (2015).

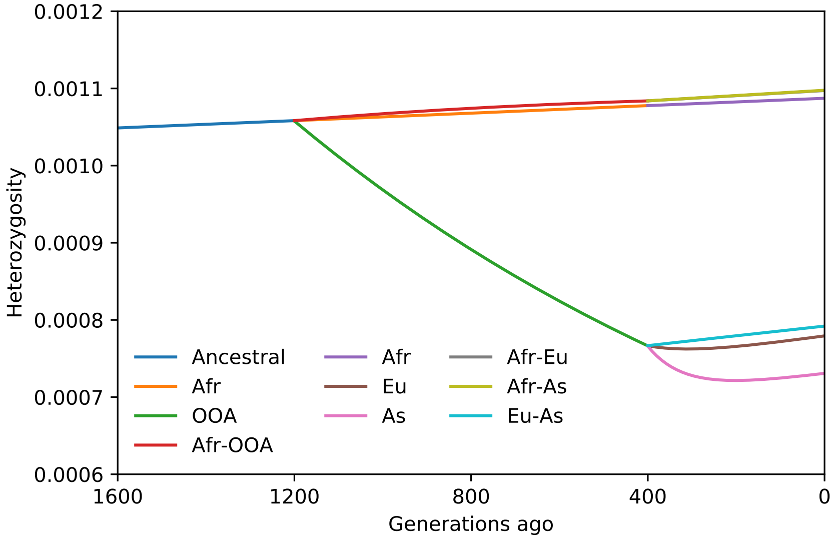

Figure S3: **Within and between population heterozygosity.** Toy model for out-of-Africa expansion, with subsequent migration between split populations. The OOA population experiences a steady decay of heterozygosity due to the prolonged bottleneck, and different bottleneck strengths and exponential growth rates between more recent Eu and As populations account for differences in observed heterozygosity in those populations. Drift does not directly affect cross-population heterozygosity, which increases linearly in the absence of migration, and more slowly with low levels of migration. Strong migration would lead to cross-population $H$ intermediate between the two populations.

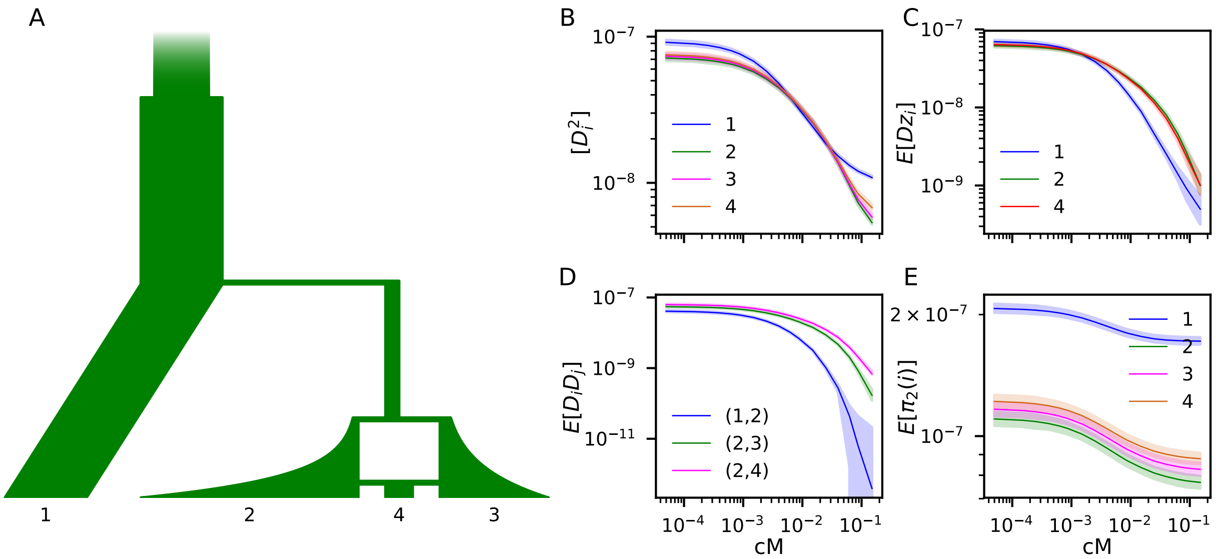

Figure S4: **Validation: computed LD curves match simulation.** (A) We simulated 200 replicates of 100Mb genome using `msprime` (Kelleher *et al.*, 2016) under the illustrated four-population demography. Demographic parameters for this simulation were $\nu_A = 1.5$, $T_A = 0.3$, $\nu_B = 0.25$, $T_B = 0.16$, $\nu_{2,0} = 0.1$, $\nu_{2_f} = 4.0$, $\nu_{3,0} = 0.2$, $\nu_{3,f} = 2.0$, $T_3 = 0.06$, $\nu_4 = 0.5$, $T_4 = 0.01$, and $f = 0.5$, where $\nu_i$ is the relative size of population $i$ compared to the reference ancestral size, $N_e = 10^4$. $u = 2 \times 10^{-8}$ and $r = 2 \times 10^{-8}$ per base pair. (B-E) Shaded regions indicate 95% confidence intervals of statistics from the 200 simulations, while solid curves are expectations from `moments.LD`.

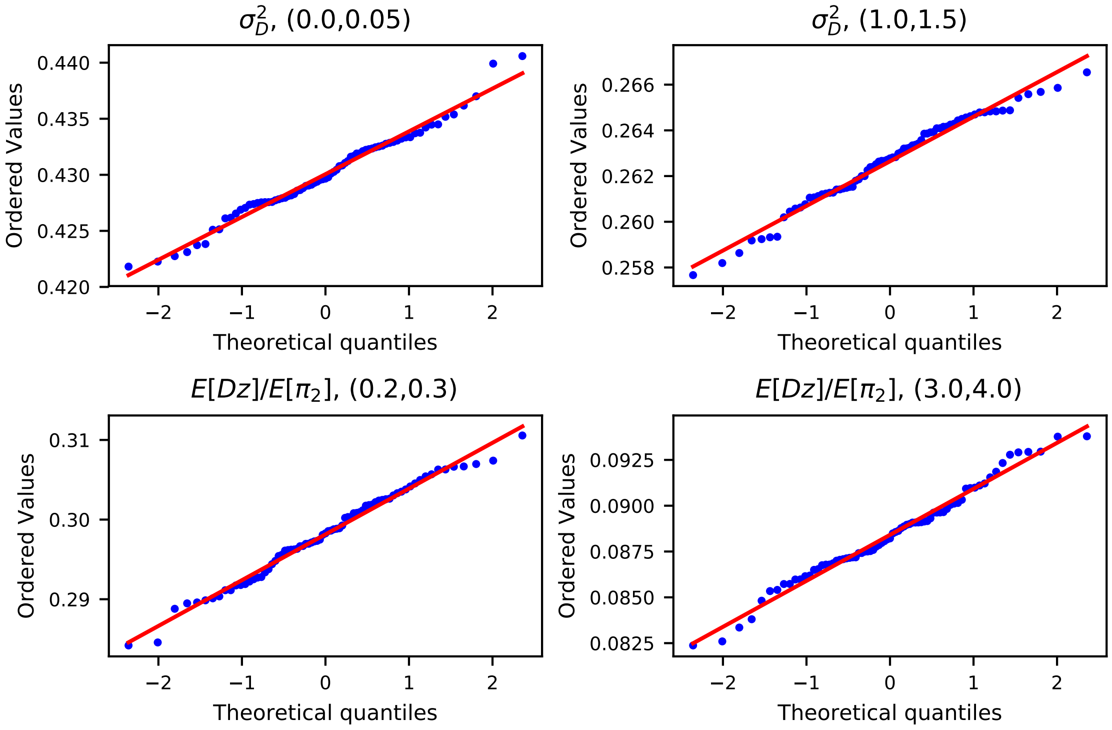

Figure S5: **Hill-Robertson statistics are approximately normally distributed.** Here, we used `msprime` (Kelleher *et al.*, 2016) to simulate 100 replicates of a two-epoch demography with recent growth, and computed $\sigma_d^2$ and $\mathbb{E}[Dz]/\mathbb{E}[\pi_2]$ for multiple recombination distance bins.

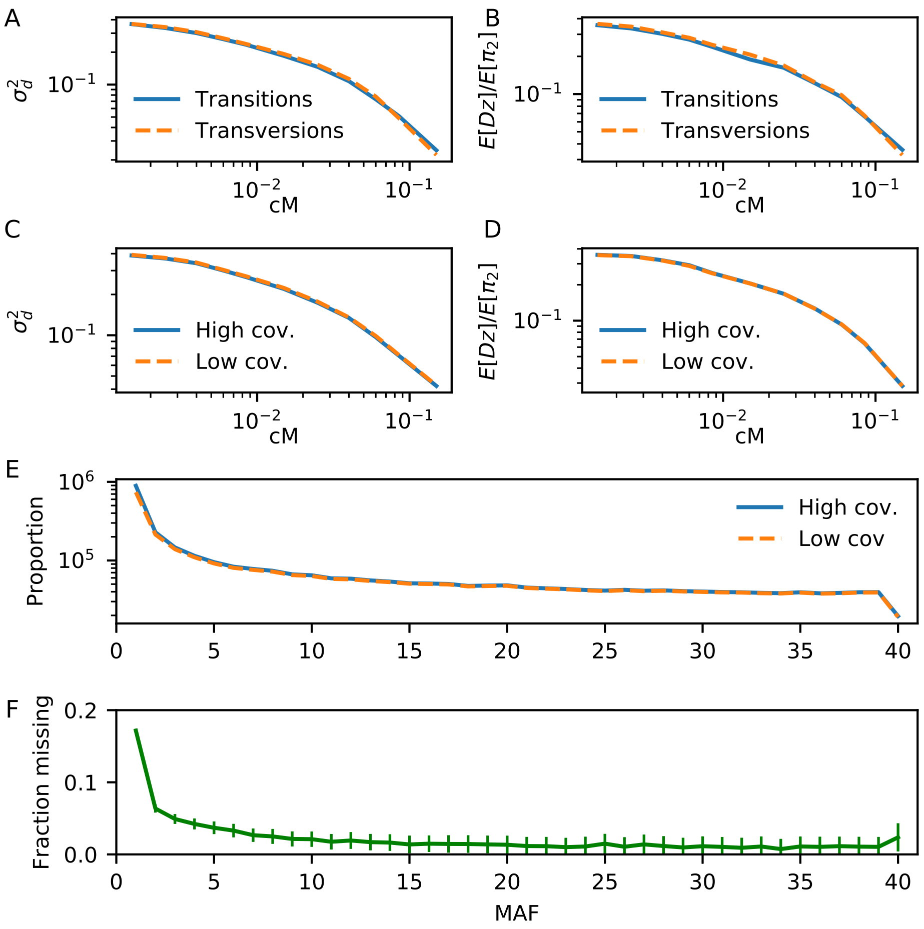

Figure S6: **Effect of mutation types and low coverage on Hill-Robertson statistics.** We used 40 individuals that overlapped between the 1000 Genomes data and the 90 Han Chinese data to compute (A,C) $\sigma_d^2$, (B,D) $\mathbb{E}[Dz]/\mathbb{E}[\pi_2]$ and (E) the folded AFS across intergenic sites. (A-B) In our analyses in the main text, we used all mutations (transitions and transversions). Here, we compare LD curves for statistics estimated from transitions (solid, blue) or transversions (dashed, orange) only. Differences in statistics between the two mutation types are negligible. (C-D) The 90 Han Chinese data was high coverage, while 1000 Genomes data (which we used in our analysis) was low coverage. The LD curves are largely unaffected by the level of low coverage in the 1000 Genomes data. (E-F) For comparison, the allele frequency spectrum is sensitive to coverage, as the singleton bin of the AFS is significantly underestimated in the 1000 Genomes data (F).

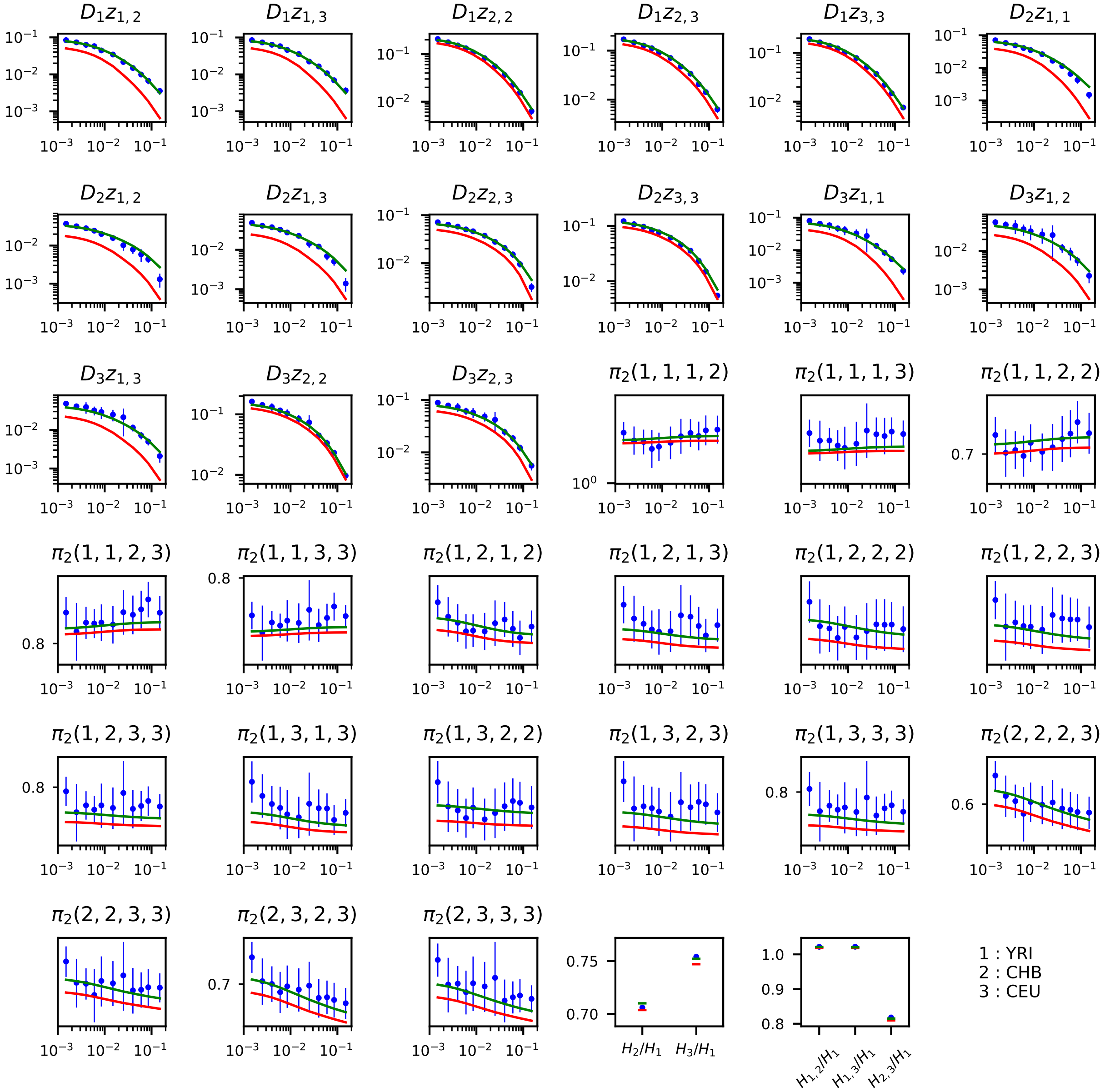

Figure S7: **Additional statistics from model fits.** Figures 3 and 4 compare model predictions to a handful of observed statistics in the multi-population basis. Here, we show comparisons for the remainder of the statistics used in the fits, and each two-locus statistic is normalized by $\pi_2(1,1,1,1) = \pi_2(\text{YRI})$. Single-locus statistics are normalized by $H(\text{YRI})$. The horizontal axis is given in units of cM. Indices in the titles indicate populations: YRI is population 1, CHB is population 2, and CEU is population 3. Red curves: standard OOA model. Green curves: OOA model with archaic branches. Error bars on the data indicate 95% confidence intervals of estimates. Best fit parameters and 95% confidence intervals for each are given in Table S2.

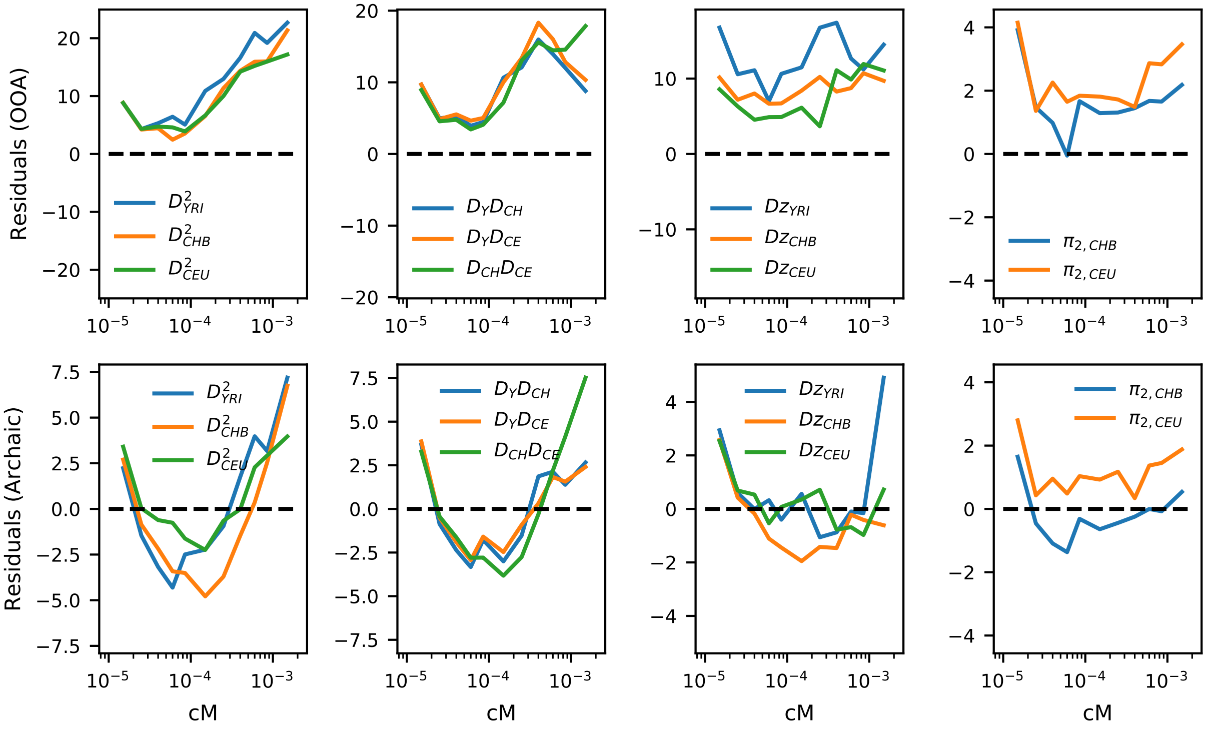

Figure S8: **Residuals from the OOA and archaic models.** Residuals are shown for the best fit OOA model (top, illustrated in Figure 3) and the best fit model with archaic admixture (bottom, from Figure 4). Residuals were computed as $(D_i - E_i)/V_i^{1/2}$, where $D_i$ is the observed data for a given statistic and bin, $E_i$ is the expected statistic from the best fit model, and $V_i$ is the variance of the statistics computed through bootstrap resampling.

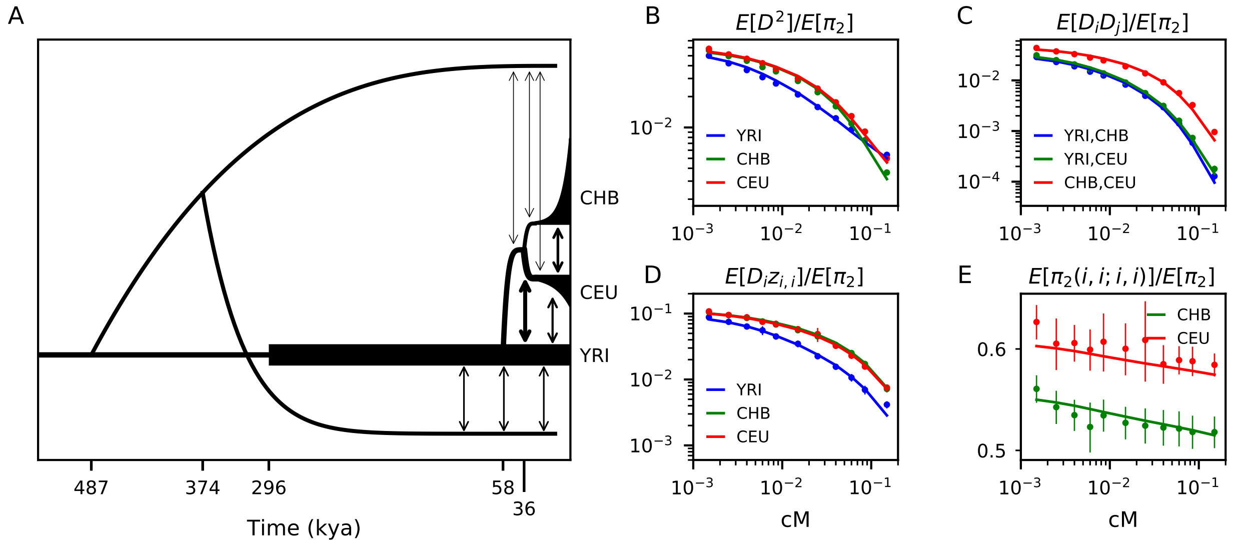

Figure S9: **Alternate topology of archaic branches.** (A) In addition to the scenario where each archaic branch splits independently from the modern human branch (Figure 4), we considered a model where a single archaic lineage splits from modern humans, and then some time later splits into the Eurasian and African archaic branches. Aside from the archaic split times and topology, parameterization was the same between the two models. (B-E) This archaic admixture model provided a good fit the LD data, roughly equal to the archaic admixture model shown in the main text.

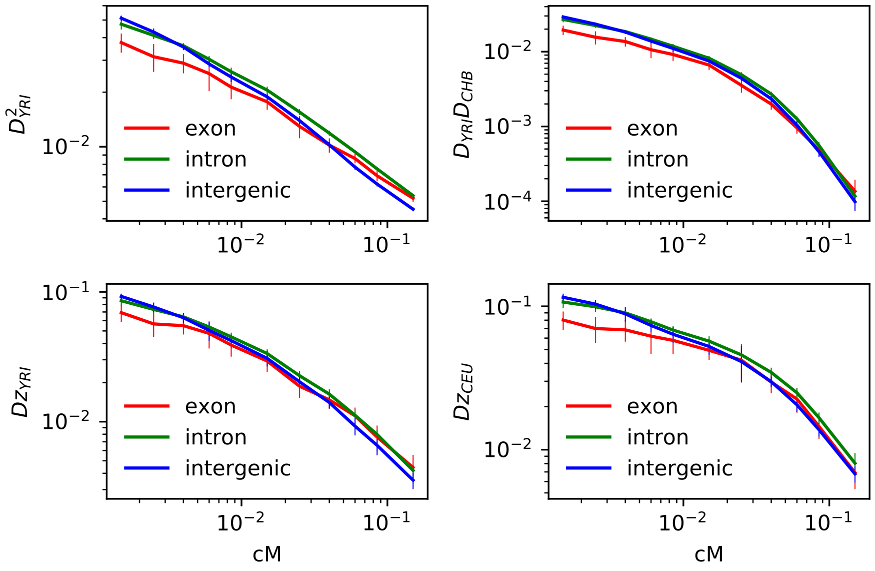

Figure S10: **LD statistics from genome regions.** We compared intergenic data, which we used in our analyses, to LD decay curves from intron and exon regions. Each statistic is normalized by $\pi_2(YRI)$. Exon regions have LD decay curves that differ significantly from intron and intergenic regions, and intron and intergenic regions also differ. Selection is known to affect the expected AFS and LD, so we excluded genic regions from our analyses.

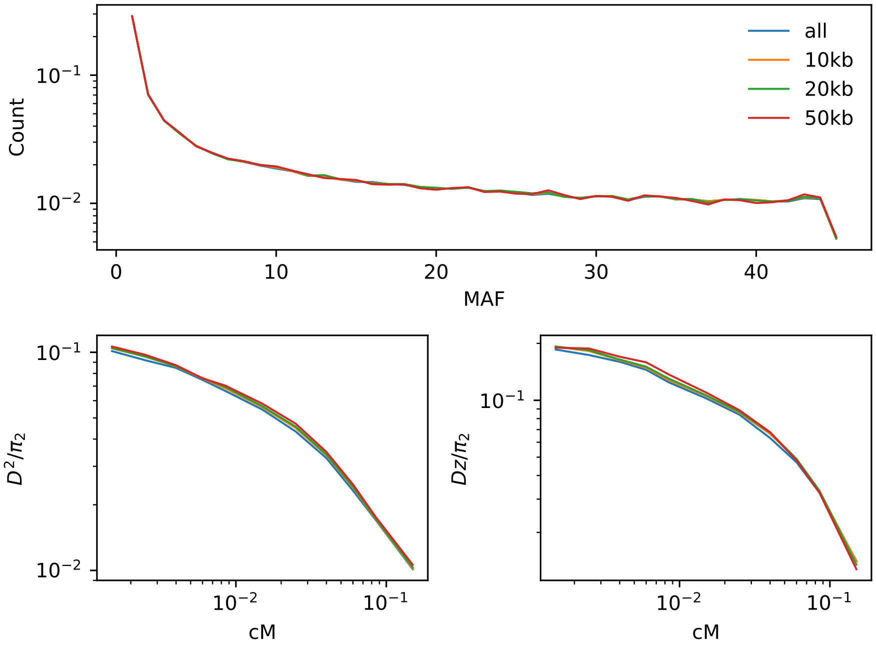

Figure S11: **LD statistics from intergenic regions.** We compared statistics for SNPs across all intergenic regions to SNPs from intergenic regions at least a given distance from the nearest gene. Overall, statistics were similar for each choice. We chose to include all intergenic SNPs in order to increase the number of observed pairs in our data.

| | Relative error($\times 10^{-3}$) | | Comp. time (minutes) | |
|---|---|---|---|---|
| Model | ∂a∂i | moments | ∂a∂i | moments |
| Equilibrium, $n = 30$, $\rho = 0$ | 1.8 | 0.0017 | 0.25 | $< 10^{-3}$ |
| Equilibrium, $n = 30$, $\rho = 10$ | 15.6 | 0.40 | 0.25 | $< 10^{-3}$ |
| Equilibrium, $n = 50$, $\rho = 0$ | 4.6 | 0.0033 | 3.5 | $< 10^{-3}$ |
| Equilibrium, $n = 50$, $\rho = 10$ | 34 | 0.11 | 3.5 | $< 10^{-3}$ |
| Bottleneck, $n = 30$, $\rho = 0$ | 0.41 | 0.031 | 7.8 | 0.037 |
| Bottleneck, $n = 30$, $\rho = 10$ | 38.32 | 0.25 | 7.9 | 0.054 |
| Bottleneck, $n = 50$, $\rho = 0$ | 3.97 | 0.036 | 21.8 | 0.60 |
| Bottleneck, $n = 50$, $\rho = 10$ | 63.53 | 0.058 | 22.3 | 0.94 |

Table S1: **Comparison of `moments.TwoLocus` and `∂a∂i.TwoLocus`.** For the equilibrium distribution, we compared to Hudson's 2001 implementation for $n = 100$ projected down to sample sizes $n = 30$ and $n = 50$. For the bottleneck distribution, we computed a numerical approximation with a larger sample size ($n = 80$) and shorter integration time step using `moments` and then projected to the required size. We measured relative error as $\sum(\Psi_n(\text{model}) - \Psi_n(\text{True}))^2/\Psi_n(\text{True})$. Equilibrium solutions were cached, as is default in both programs, but `∂a∂i` still needs to integrate $\psi$ against the multinomial distribution to obtain $\Psi_n$, accounting for the time differences in the Equilibrium case. With recombination, `moments` improves in accuracy as $n$ increases, since the Jackknife approximation becomes more and more accurate with larger $n$.

| Model | Archaic Admixture B | |
|---|---|---|
| Parameter | ML Estimates | 95% CI |
| $N_0$ | 3700 | 3020 – 4370 |
| $N_{\text{YRI}}$ | 14000 | 11800 – 16100 |
| $N_{\text{B}}$ | 860 | 110 – 1610 |
| $N_{\text{CEU0}}$ | 2300 | 1430 – 3210 |
| $r_{\text{CEU}}(\%)$ | 0.122 | 0.081 – 0.149 |
| $N_{\text{CHB0}}$ | 650 | 340 – 960 |
| $r_{\text{CHB}}(\%)$ | 0.362 | 0 – 0.435 |
| $m_{\text{AF - B}}(\times 10^{-5})$ | 53.4 | 11.8 – 95.0 |
| $m_{\text{YRI - CEU}}(\times 10^{-5})$ | 2.43 | 1.62 – 3.24 |
| $m_{\text{YRI - CHB}}(\times 10^{-5})$ | 0 | — |
| $m_{\text{CEU - CHB}}(\times 10^{-5})$ | 11.6 | 7.27 – 15.9 |
| $T_{\text{AF}}$ (kya) | 296 | 244 – 347 |
| $T_{\text{OOA}}$ (kya) | 57.7 | 35.4 – 80.0 |
| $T_{\text{CEU - CHB}}$ (kya) | 36.3 | 28.8 – 43.8 |
| $T_{\text{Arch. split}}$ (kya) | 487 | 167 – 807 |
| $T_{\text{Arch. Af. - Nean. split}}$ (kya) | 374 | 25.5 – 723 |
| $T_{\text{Arch. Af. mig.}}$ (kya) | 110 | 0 – 476 |
| $m_{\text{AF - Arch. Af.}}(\times 10^{-5})$ ... | 2.43 | 0 – 5.24 |
| $m_{\text{OOA - Nean}}(\times 10^{-5})$ | 1.51 | 0 – 3.19 |
| $T_{\text{Arch. adm. end}}$ (kya) | 20.7 | 16.3 – 25.1 |

Table S2: **Maximum likelihood parameters for alternate archaic topology.** For the most part, estimates were qualitatively similar to the archaic model presented in the main text. In each model, the split between modern humans and the branch leading to the archaic African population occurred about 500 kya. However, here the Neanderthal lineage split from this branch more recently than 500 kya, which is considerably more recent than most estimates or our estimate from the alternative model. This is expected, as the largest discrepancy between the non-archaic model and data occurred for $Dz$ in African populations, so the inference of the split date in this model is primarily driven by the signal in YRI. Without including archaic genomes in this analysis, we did not have statistical power to discriminate between the two proposed topologies.

| Model | OOA (fit w/o $Dz$) | | Archaic Admixture | |
|---|---|---|---|---|
| Parameter | Estimates | 95% CI | Estimates | 95% CI |
| $N_0$ | 2360 | 2140 – 2580 | 2860 | 2460 – 3260 |
| $N_{\text{LWK}}$ | 14600 | 11500 – 17700 | 15300 | 10400 – 20300 |
| $N_{\text{B}}$ | 1130 | 700 – 1570 | 1020 | 730 – 1310 |
| $N_{\text{GBR0}}$ | 1560 | 760 – 2370 | 2210 | 1500 – 2920 |
| $r_{\text{GBR}}(\%)$ | 0.229 | 0.011 – 0.293 | 0.157 | 0.135 – 0.175 |
| $N_{\text{KHV0}}$ | 390 | 0 – 980 | 630 | 480 – 790 |
| $r_{\text{KHV}}(\%)$ | 0.680 | 0 – 0.888 | 0.471 | 0 – 0.545 |
| $m_{\text{AF - B}}(\times 10^{-5})$ | 48.9 | 29.2 – 68.6 | 50.7 | 41.2 – 60.2 |
| $m_{\text{LWK - GBR}}(\times 10^{-5})$ | 2.71 | 0 – 8.92 | 2.87 | 1.45 – 4.29 |
| $m_{\text{LWK - KHV}}(\times 10^{-5})$ | 0 | 0 – 1.32 | 0 | — |
| $m_{\text{GBR - KHV}}(\times 10^{-5})$ | 9.94 | 4.41 – 15.5 | 7.29 | 0 – 16.8 |
| $T_{\text{AF}}$ (kya) | 215 | 139 – 290 | 249 | 219 – 279 |
| $T_{\text{OOA}}$ (kya) | 68.2 | 52.6 – 83.7 | 61.5 | 44.0 – 79.0 |
| $T_{\text{GBR - KHV}}$ (kya) | 28.3 | 21.9 – 34.8 | 30.9 | 26.2 – 35.5 |
| $T_{\text{Arch. Af. split}}$ (kya) | — | | 511 | 456 – 566 |
| $T_{\text{Arch. Af. mig.}}$ (kya) | — | | 250 | 160 – 341 |
| $m_{\text{AF - Arch. Af.}}(\times 10^{-5})$ | — | | 0.752 | 0.288 – 1.22 |
| $T_{\text{Nean.}}$ (kya) | — | | 540 | 381 – 700 |
| $m_{\text{OOA - Nean}}(\times 10^{-5})$ | — | | 0.414 | 0 – 0.993 |
| $T_{\text{Arch. end}}$ (kya) | — | | 13.0 | 4.3 – 21.7 |

Table S3: **Models fits to alternate trio.** We fit the same out-of-Africa model with and without archaic branches to a separate trio in the 1000 Genomes data: (Luhya from Kenya (LWK), Kinh from Vietnam (KHV), and British from England and Scotland (GBR)). Best fit parameters compare qualitatively to those fit to the YRI-CHB-CEU data, although confidence intervals were wider for this trio.
